# Human BioMolecular Atlas Program (HuBMAP): 3D Human Reference Atlas Construction and Usage

**DOI:** 10.1101/2024.03.27.587041

**Authors:** Katy Börner, Philip D. Blood, Jonathan C. Silverstein, Matthew Ruffalo, Rahul Satija, Sarah A. Teichmann, Gloria Pryhuber, Ravi S. Misra, Jeffrey Purkerson, Jean Fan, John W. Hickey, Gesmira Molla, Chuan Xu, Yun Zhang, Griffin Weber, Yashvardhan Jain, Danial Qaurooni, Yongxin Kong, HRA Team, Andreas Bueckle, Bruce W. Herr

**Affiliations:** Department of Intelligent Systems Engineering, Luddy School of Informatics, Computing, and Engineering, Indiana University, Bloomington, IN, USA; CIFAR MacMillan Multiscale Human program, CIFAR, Toronto, ON, Canada; Pittsburgh Supercomputing Center, Carnegie Mellon University, Pittsburgh, PA, USA; Department of Biomedical Informatics, University of Pittsburgh School of Medicine, Pittsburgh, PA, USA; Ray and Stephanie Lane Computational Biology Department, Carnegie Mellon University, Pittsburgh, PA, USA; New York Genome Center, New York, NY, USA; Wellcome Sanger Institute, Wellcome Genome Campus, Hinxton, Cambridge, UK; University of Rochester Medical Center, Rochester, NY, USA; Department of Biomedical Engineering, Johns Hopkins University, Baltimore MD, USA; Department of Biomedical Engineering, Duke University, Durham, NC, USA, New York Genome Center, New York, NY, USA; J. Craig Venter Institute, La Jolla, CA, USA; Department of Biomedical Informatics, Harvard Medical School, Boston, MA, USA

## Abstract

The Human BioMolecular Atlas Program (HuBMAP) aims to construct a reference 3D structural, cellular, and molecular atlas of the healthy adult human body. The HuBMAP Data Portal (https://portal.hubmapconsortium.org) serves experimental datasets and supports data processing, search, filtering, and visualization. The Human Reference Atlas (HRA) Portal (https://humanatlas.io) provides open access to atlas data, code, procedures, and instructional materials. Experts from more than 20 consortia are collaborating to construct the HRA’s Common Coordinate Framework (CCF), knowledge graphs, and tools that describe the multiscale structure of the human body (from organs and tissues down to cells, genes, and biomarkers) and to use the HRA to understand changes that occur at each of these levels with aging, disease, and other perturbations. The 6th release of the HRA v2.0 covers 36 organs with 4,499 unique anatomical structures, 1,195 cell types, and 2,089 biomarkers (e.g., genes, proteins, lipids) linked to ontologies and 2D/3D reference objects. New experimental data can be mapped into the HRA using (1) three cell type annotation tools (e.g., Azimuth) or (2) validated antibody panels (OMAPs), or (3) by registering tissue data spatially. This paper describes the HRA user stories, terminology, data formats, ontology validation, unified analysis workflows, user interfaces, instructional materials, application programming interface (APIs), flexible hybrid cloud infrastructure, and previews atlas usage applications.

## Main

Inaugurated in 2018, the Human BioMolecular Atlas Program (HuBMAP) aims to construct a comprehensive reference model of the healthy (i.e., ’non diseased’) human body across all levels, from organs and tissues down to cells and canonical biomarkers^1,2^. The Human Reference Atlas (HRA)^3^ includes a Common Coordinate Framework (CCF; see **Box 1**) which helps harmonize multimodal data, including 3D organ models, histology images, and omics data from profiling of single cells. HRA data comprise human expert generated information (e.g., anatomical systems, anatomical structures, cell types, biomarker [ASCT+B] tables, and 2D and 3D reference objects), experimental data mapped to the HRA, as well as enriched atlas data in support of different atlas applications. The origin and evolution of HRA and ASCT+B tables are detailed in prior work^3^.

The CCF provides quantitative workflows for integrating new experimental data into the growing atlas, such as histology images, vascular pathways, and single cell analyses. The resulting HRA provides data evidence for common states of cells and anatomical structures in the human body at specific 3D locations, and this can be used as a canonical reference to describe the changes that occur across biological variables (e.g., age, sex, race, body mass) and acute or chronic diseases. It can benefit applications such as drug development by providing a better understanding of perturbations of cell types and states in diseased conditions, which could reveal relevant targets for precision medicine through comparisons of a diseased tissue to a non-diseased, but CCF matched reference.

When HuBMAP was launched, the first unifying concepts that map major organs in the human body across scales were emerging^4,5^. Existing atlases used organ-specific references (e.g., Waxholm Space for the brain^6^ or the Helmsley 1D distance reference system for the colon^7^), but most of these references do not map to a shared human body CCF. To advance CCF development, in March 2020, the National Institutes of Health (NIH) and the Human Cell Atlas (HCA)^8^ consortium organized a joint virtual meeting with a CCF breakout session. This resulted in the formation of the HRA Working Group (WG). Over the last 49 months, WG members jointly developed a definition and key properties for the human reference atlas. These properties are:

1. **The HRA defines a reference 3D multiscale space and shape of anatomical structures and the cell types and biomarkers used to characterize them**. Anatomical structures, cell types, and biomarkers are validated and represented in or are added to existing ontologies (e.g., the uber-anatomy ontology (Uberon)^9^, the Foundational Model of Anatomy Ontology [FMA]^10,11^, Cell Ontology [CL]^12^, Provisional Cell Ontology [PCL]^13^, and the Human Gene Ontology Nomenclature Committee [HGNC, https://www.genenames.org]). As more data are collected, the HRA will increasingly be able to show how body shape and size differ across individuals and change over a person’s lifespan.
2. **The HRA enables adding new experimental datasets and mapping these to existing data through a variety of mechanisms**. For example, the location of tissue specimens can be specified relative to virtual 3D reference organ models in the HRA; single cell genomic data can be mapped to the HRA using annotation tools like Azimuth^14^; and, single cell-resolution spatial proteomics data can be mapped to the HRA using Organ Mapping Antibody Panels (OMAP)^15^. With the development of new technologies and computational methods in the future, additional mappings and linkages, such as integration of multi-omics data, will be possible.
3. **The HRA follows best practices and standards for sharing scientific data**. To do this, the HRA needs to be authoritative (i.e., it should be supported by: peer-reviewed scholarly publications, experimental data evidence, or expert consensus); meet the Transparency, Responsibility, User focus, Sustainability and Technology (TRUST) principles for digital repositories^16^; be representative (i.e., covering all major human demographics and welcoming everyone to contribute to and use the HRA data); be open and adhere to the Findable, Accessible, Interoperable, Reusable (FAIR) principles^6^ (i.e., anyone can use the HRA data and code and these are provided in community standard formats with linked ontologies; be published as Linked Open Data (LOD) connected to ontologies and other LOD; API queries, and user interfaces are supported); detailed protocols and SOPs are provided; and be continuously evolving (e.g., as new technologies, data, and methods become available).

Experts also agreed on key Standard Operating Procedures (SOPs) for atlas construction and usage plus HRA terminology, see **Box 1**, adopted from the HRA SOPs Glossary^17^. Plus, the HRA WG brought together technical leads from the HuBMAP Integration, Visualization & Engagement (HIVE) Collaboratory with experimental teams in HuBMAP and experts from the Genotype-Tissue Expression (GTEx)^18^, GUDMAP: GenitoUrinary Developmental Molecular Anatomy Project^19^, Kidney Precision Medicine Project (KPMP)^20,21^, LungMAP^22^, BRAIN Initiative Cell Census Network Initiative (BICCN)^23,24^, Cellular Senescence Network (SenNet)^25^ and other NIH-funded consortia and with strong support from the Human Cell Atlas effort (HCA)^8,26^ to develop the HRA data, code, and portal infrastructure together.

#### Box 1. Key HRA Terminology

- **3D collision:** The intersection of 2D or 3D bounding box volumes or surface polygon meshes.
- **3D model:** Digital object with a shape and size defined by polygon meshes (vertices and edges) that can be used to represent the real-world form of anatomical structures, cells, or proteins in 3D.
- **3D reference object:** Polygon mesh of 3D spatial objects (e.g., anatomical structures), their object node hierarchy, materials, and surface color and texture.
- **Anatomical Structure (AS):** A distinct biological entity with a 3D volume and shape, e.g., an organ, functional tissue unit (FTU), or cell.
- **Antibody Validation Report (AVR)**: Document providing details on the characterization of individual antibodies for multiplexed antibody-based imaging assays. These details include target protein information (e.g., target name, UniProt accession number), and antibody information (e.g., RRID, host organism, vendor, catalog number, lot number). AVRs also provide details on controls used for antibody characterization and validation (positive and negative tissues, cell lines, isotype controls, etc), exemplar imaging data, and information on other antibodies tested.
- **ASCT+B Tables:** Anatomical Structures, Cell Types and Biomarkers (ASCT+B) Tables are authored by multiple experts across many consortia. Tables capture the partonomy of anatomical structures, cell types, and major biomarkers (e.g., gene, protein, lipid, or metabolic markers) defining cellular identity supported by scientific evidence and are linked to ontologies.
- **Atlas enriched dataset graph:** A graph for highest quality datasets used for HRA construction (i.e., it has an extraction site, cell type population, and publication or provenance on a major atlas portal), enriched with additional metadata, computed by HRApop.
- **Biomarkers (B):** HRA biomarkers are used to *characterize* or identify cell types. They include genes (BG), proteins (BP), metabolites (BM), proteoforms (BF), and lipids (BL).
- **Cell Suspension**: Single cells or nuclei isolated from a tissue (e.g., using enzymes or mechanical means) for single cell assays, e.g., before sc/snRNA-seq assay is run.
- **Cell Types (CT):** Tissue is composed of different (resident and transitory) cell types that are *characterized* or identified via biomarkers.
- **Cell Type Annotation (CTann):** Azimuth and other cell type annotation tools are used to assign cell types to cells from sc/snRNA-seq studies. Manually compiled crosswalks are used to assign ontology IDs to CTann cell types.
- **Cell Type Population (CTpop):** A listing of unique cell types, the number of cells per cell type, and mean biomarker expression values per cell type computed for anatomical structures, extraction sites, and datasets.
- **Common Coordinate Framework (CCF):** The HRA CCF consists of ontologies and reference object libraries, computer software (e.g., user interfaces) and training materials that (1) enable biomedical experts to semantically annotate tissue samples and to precisely describe their locations in the human body (“registration”), (2) align multi-modal tissue data extracted from different individuals to a 3D reference coordinate system (“mapping”), and to (3) provide tools for searching and browsing HuBMAP data at multiple levels from the whole body down to single cells (“exploration”). Alternative CCF definitions do exist^27^.
- **Cosine similarity:** Measures the cosine of the angle between two vectors, with 1 indicating identical vector directions and 0 indicating no similarity.
- **Crosswalk:** An ontological mapping of terms in HRA digital objects (e.g., 2/3D reference objects, OMAPs, Azimuth references) to ontology terms in the ASCT+B tables.
- **Dataset graph:** A JSON-LD file containing a graph of RUI-registration, donor, experimental (e.g., links to cell by biomarker [CxB] H5AD files or CTpop), literature, and provenance data.
- **Digital Object (DO):** Unit of information that includes properties (attributes or characteristics of the object) and may also include methods (means of performing operations on the object).
- **Digital Object Identifier (DOI):** Centrally registered identifier composed of a string of numbers, letters and symbols used to uniquely identify an article or document with a permanent web address or Uniform Resource Locator (URL).
- **Extraction Site**: Digital, 3D representation of a tissue block. If the RUI is used to register tissue, then each site has a unique ID; data on size, location, and rotation in 3D in relation to a HRA reference organ; a listing of all anatomical structures that the cuboid intersects with (bounding-box collision by default); and metadata on who registered it.
- **FAIR principles:** Acronym for Findable, Accessible, Interoperable, and Reusable, which is a way of sharing data to maximize its utility^6^.
- **Functional tissue unit (FTU):** A small tissue organization (i.e., set of cells) that performs a unique physiologic function and is replicated multiple times in an organ. Examples are liver lobule, alveolus of lung, or pancreatic acinus.
- **Hematoxylin and eosin (H&E) stain:** Histology stain that is widely used as a gold standard for pathological evaluation of tissue sections.
- **Human Reference Atlas (HRA)**: The HRA is a comprehensive, high-resolution, three-dimensional atlas of major cells in the healthy human body. The HRA provides standard terminologies and data structures for describing specimens, biological structures, and spatial positions linked to existing ontologies.
- **Human Reference Atlas Literature (HRAlit):** Scholarly publication linked to HRA DOs to provide scholarly evidence.
- **Human Reference Atlas Population (HRApop):** Experimental data linked to HRA DOs to provide data evidence and number of cells per cell type per 3D anatomical structure.
- **Linked Open Data (LOD):** A method for sharing data in standard, non-proprietary formats, and deeply interlinked with other data resources.
- **Millitome:** A device used to hold and slice organs into a grid of equally sized tissue blocks plus a process of generating HRA-aligned digital tissue blocks.
- **Ontology:** A set of subject area concepts (here, anatomical structures, cell types and biomarkers), their properties and the relationships between them. Ontologies used in the HRA include Uberon and Cell Ontology (CL).
- **Organ Mapping Antibody Panel (OMAP):** A comprehensive panel of curated antibodies that identifies the major anatomical structures and cell types in a specific organ. The selected antibodies are optimized for a tissue preservation method (i.e., fixed, frozen) and multiplexed imaging modality (e.g., CODEX, Cell DIVE) through published protocols (protocols.io) and AVR.
- **Partonomy:** A classification hierarchy that represents part–whole relationships.
- **Polygon mesh:** A collection of vertices, edges and faces defining the shape of a polyhedral object, e.g., tissue block cuboids or reference objects denoting anatomical structures.
- **Reference Objects**: A 3D model of anatomical structures created by medical illustrators with the involvement of subject matter experts following standard operating procedures.
- **Registration Set (see also Dataset graph):** Grouping of tissue blocks by the study/paper in which they were published. Each set has a human-readable registration set name, a machine readable Internationalized Resource Identifier (IRI), and it links to a paper DOI and tissue block ID.
- **RUI-registered:** Tissue spatially and semantically registered to the HRA using the Registration User Interface (RUI).
- **Segmentation:** Image processing that predicts boundaries of objects, e.g., anatomical structures such as nuclei, cellular membranes, cells, or FTUs in 2D or 3D.
- **Single cell transcriptomic data, sc/snRNA-seq:** Data from single cell or single nucleus (sc/sn), high-throughput suspension-based studies that measure polyadenylated ribonucleic acid (RNA) molecules in an individual cell.
- **Single cell proteomic data:** Data from single cell studies using CODEX, Cell DIVE, Ibex, CycIF, or other assays that detect proteins in situ in a tissue, consequently enabling protein expression quantification.
- **Tissue Block:** A sample or specimen derived from an organ or tissue including subsections thereof obtained from a donor that has a unique ID and links to donor, organ extraction site, processing, preservation and other metadata. The locations of tissue blocks are registered using the RUI.
- **Tissue Section**: A thin (several microns) section of a tissue block usually obtained using a cryotome or microtome. Tissue sections inherit the location and rotation of their parent tissue block. The thickness and number of tissue sections is captured in an input field inside the RUI.
- **Typology:** A classification that represents general types, e.g., cell types or biomarker types.
- **United File**: A GLB 3D object that contains all the modeled organs in the Human Reference Atlas.
- **Vascular Common Coordinate Framework (VCCF)**: A proposed approach to use vasculature as a coordinate system to map all the cells in the human body.
- **Web Ontology Language (.OWL) file format:** The World Wide Web Consortium (W3C) Web Ontology Language is a Semantic Web language documented at https://www.w3.org/OWL.

An important next step in the HRA effort was collecting user stories in support of atlas construction and usage to encourage dialogue, deliberation, and iteration by designers and users around three key questions: Who are the users involved in a particular user story? What are the outcomes they hope to achieve? What value do they stand to gain? Agreement on answers to these questions helped prioritize user needs, provided context for a proposed user story, and reduced ambiguity.

More than 30 one-on-one interviews were conducted with atlas architects, i.e., experts who serve as principal investigators or are otherwise intimately involved in the construction of the latest generation of human atlases, including BICCN, GTEx, GUDMAP, HCA, HuBMAP, Human Tumor Atlas Network (HTAN)^28^, KPMP, LungMAP, (Re)building the Kidney (RBK)^29^, and SenNet. Given the interdisciplinary nature of this effort, the atlas architects who were interviewed comprise a diverse group of physicians, laboratory and computational biologists, engineers, and computer and data scientists. In addition, six programmers from different human atlas projects were surveyed.

The interview and survey results helped identify three key objectives and seven concrete user stories (US#1-7) for the construction, functionality, and sustainability of the HRA (**Table 1**). The three objectives are:

1. The HRA should facilitate atlas construction by aligning new tissue blocks with existing data. For example, developers of the HRA want to predict cell type populations for new tissue blocks (US#1 in **Table 1**) and predict the spatial origin of tissue samples with known cell type populations (US#2).
2. The HRA should contain functionality that provides insights into changes (e.g., with aging, disease, or other perturbations) that occur at all levels in the body. To do this, researchers and clinicians need to be able to search for and explore biomarker expression values for cell types and functional tissue units (FTUs) (US#3 and US#4) and determine the location and distances between cells (US#5).
3. The HRA should use processes that encourage collaboration and guide future development to ensure long-term sustainability. This includes leveraging architectures with modular, lightweight components that can be easily shared (US#6) with metrics of success provided via an HRA dashboard to funders and the public in order to gain feedback and support (US#7).

**Table 1:** User stories. Feature summary, target user roles, user activities, and added value for seven user stories that drive HRA development.

| Feature | User Role | User Activities | Added Value |
| --- | --- | --- | --- |
| *Facilitate atlas **construction** by aligning new tissue blocks with existing data* |  |  |  |
| **US#1.** Predict cell type populations | Programmers that support<br>Researchers,<br>Clinicians,<br>Pathologists | Predict and explore the likely cell type populations for a RUI-registered tissue block. | Improve cell type annotation through information on what cell type populations exist in what anatomical structures. |
| **US#2.** Predict spatial origin of tissue samples | Programmers that support<br>Researchers,<br>Clinicians | Predict and explore the likely 3D location in the human body for a given tissue block with known cell type population. | Compensate for the absence of spatial origin information in many single cell datasets. |
| *Use the atlas to gain insights into changes that occur at all levels in the body with aging or disease* |  |  |  |
| **US#3.** Compare reference tissue with aging/diseased tissue | Researchers,<br>Clinicians | Compare tissue blocks, cell types, and biomarker expression levels between healthy reference tissue and aging/diseased tissue. | Understand and communicate changes in tissue structure and function with age or disease. |
| **US#4.** Compare reference Functional Tissue Units with aging/diseased FTUs | Researchers,<br>Clinicians | Compare FTUs in terms of cell types and mean biomarker expression levels for healthy reference tissue and aging/diseased tissue. | Understand and communicate changes in FTU structure and function with age or disease |
| **US#5.** Provide cell distance distribution visualizations | Researchers,<br>Pathologists | Compute, visualize, and explore distance distributions between different cells, cell types, and anatomical structures (e.g., FTUs), and cell types and morphological features (e.g., the edge of an organ). | Add granularity to our understanding of how disease develops (e.g., how tumor cells grow or metastasize) in support of targeted therapies. |
| *Ensure atlas **sustainability** with processes that encourage collaboration and guide future development* |  |  |  |
| **US#6.** Develop lightweight atlas components | Programmers that support<br>Researchers and<br>Clinicians | Implement usable and useful HRA components (interfaces and APIs) into other portals in the growing ecosystem of human atlases. | Facilitate collaboration and data/code reuse between the HRA and other portals in support of FAIR data principles. |
| **US#7.** Implement dashboard for HRA | Researchers,<br>Clinicians, Funders | Track the evolution and usage of the HRA using data, code, and portal usage statistics in aggregate and divided by portal (e.g., HuBMAP or SenNet) or PEDP survey results. | Enable evidence-based decision-making by providing insights into the atlas' construction and usage (e.g., gaps in data, application areas, user demographics, equitable access). |

The three key objectives and associated user stories help focus presentations and discussions in the monthly HRA WG; they are driving HRA development and iterative optimization. Every six months, a new HRA release is published. With every release, existing ontologies are expanded plus HRA data structures and algorithms are improved to better serve the needs of the international human atlasing community. **Fig. 1** details major components of the 6th release of the HRA and their interlinkages.

**Figure 1:**
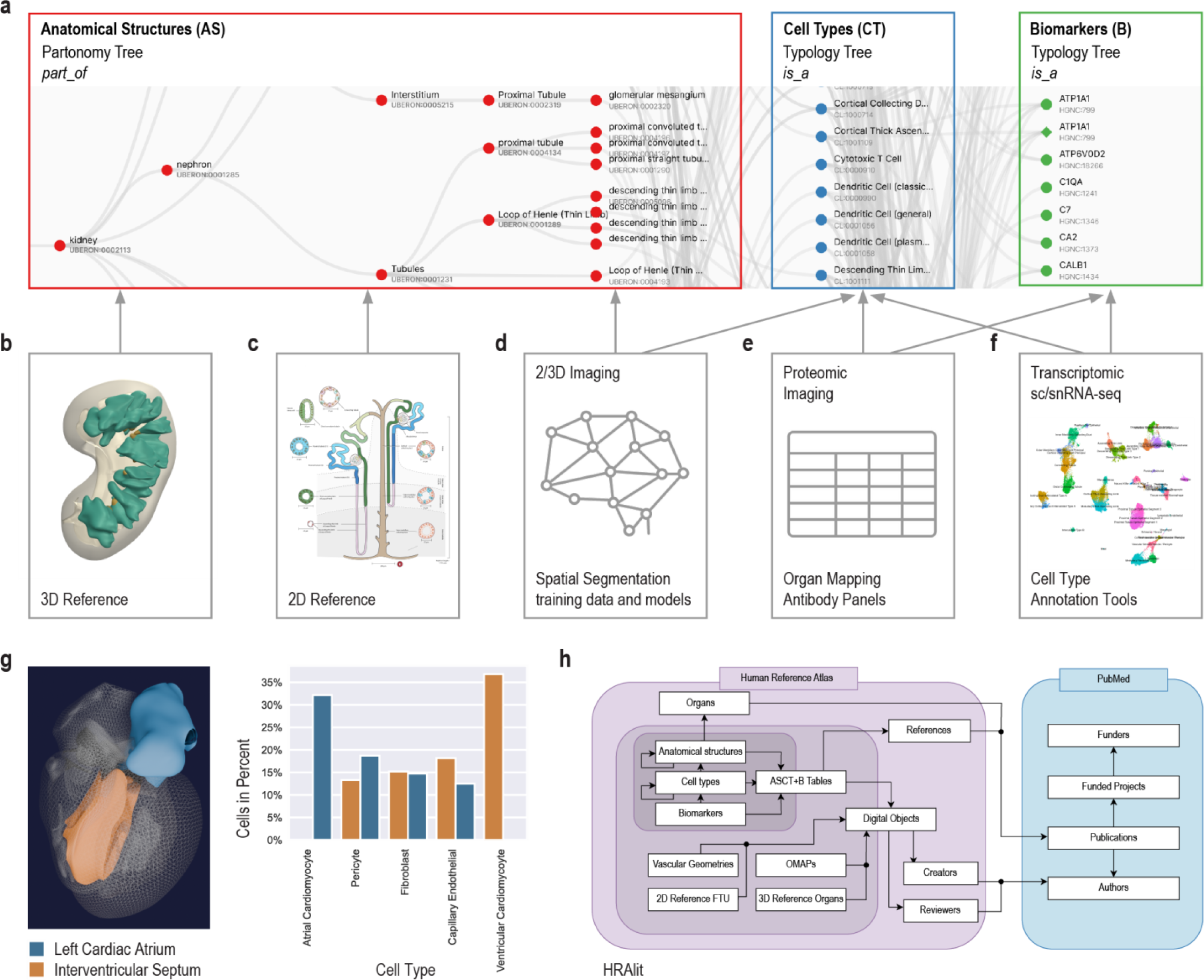
Human Reference Atlas (HRA) components and linkages. **a.** The anatomical structures, cell types and biomarkers (ASCT+B) tables document the nested *part_of* structure of organs (e.g., cells that make up functional tissue units, successively larger anatomical structures, an entire organ such as the kidney, which is *part_of* the body). The cells that make up (are *located_in*) each of the anatomical structures are organized in a multi-level cell type typology with ‘cell’ at the root and more and more specialized child nodes. The biomarkers used to *characterize* cell types might have one of five types: genes, proteins, metabolites, proteoforms, and lipids organized in a biomarker typology. Gray arrows indicate crosswalks that connect other HRA DOs to ASCT+B tables. **b.** The HRA 3D reference objects represent the shape, size, location, and rotation of 1,215 3D anatomical structures of 356 types for 65 organs with crosswalks to ASCT+B tables. Shown are ‘renal papilla’ and ‘renal pyramid’ in the kidney. **c.** 2D reference illustrations document the shape, size, and spatial layout of 3,726 2D cells of 131 types for 22 FTUs in 10 organs with crosswalks to ASCT+B tables. Shown is the kidney nephron. **d.** Labeled training data exist for FTUs in five organs with crosswalks (gray arrows) to anatomical structures and cell types in the ASCT+B tables. **e.** 13 Organ Mapping Antibody Panels (OMAPs) are linked to 197 Antibody Validation Reports (AVRs) and there exist crosswalks to cell types and biomarkers in ASCT+B tables. **f.** 10 Azimuth references for healthy adult organs plus crosswalks to cell types and biomarkers in ASCT+B tables. **g.** HRApop reports cell type populations for anatomical structures compiled from single-cell experimental data. Exemplarily shown is the left cardiac atrium (blue) and the interventricular septum (orange) of the female heart plus a bar graph with the top-five cell types that have the highest percentage across these two anatomical structures. Note that some cell types appear only in one anatomical structure. Annotations were made with Azimuth. **h.** The HRAlit database links HRA DOs to existing ontologies (e.g., Uberon, CL), expert ORCID, publication evidence, funding, and experimental data used for HRApop computation.

## Results

The HIVE Infrastructure and Engagement Component (IEC) developed HuBMAP’s flexible hybrid cloud microservices architecture (**Supplemental Figure 1** and **Methods**) to support data curation, ingestion, integration, access, analysis, exploration, and download via the HuBMAP Data Portal (https://portal.hubmapconsortium.org). HIVE Tools Components (TCs) focused on the HuBMAP Data Portal User Interface, visualization, workflow integration and tool development. HIVE Mapping Components (MCs) developed Azimuth^14^ references and the HRA Portal (https://humanatlas.io) in close collaboration with external experts.

The HuBMAP Consortium Website (**Supplemental Figure 2**) provides easy access to HuBMAP resources, publications, news, internship programs, member services, etc.. It links to the HuBMAP Data Portal and the HRA Portal. The HuBMAP Data Portal provides access to HuBMAP data, APIs, and user interfaces with continuous data and code releases. The HRA Portal serves atlas level data and code created by 18 atlas projects and new HRA releases are published every six months. Both portals use knowledge graphs (KG) to store data and the HRA KG is regularly ingested into the Unified Biomedical Knowledge Graph (UBKG, https://ubkg.docs.xconsortia.org) to link HuBMAP experimental data to existing ontologies and the HRA. The HRA uses HuBMAP and other experimental data to compute cell type populations for anatomical structures (see HRApop in **Methods** and **Supplemental Table 1**). Several HRA user interfaces (section **User interfaces**) are deployed in the HuBMAP Data Portal and other portals to support HRA construction and usage.

Atlas construction is complex and requires community agreement on data formats, APIs, and user interfaces. Previews are used to showcase and optimize new functionality before it is integrated into the HuBMAP or HRA Portal. Primary data repositories are listed in **Supplemental Table 2** and HRA code repositories in **Supplemental Table 3**. Data used in the HuBMAP Data Portal has a blue background, HRA Portal in red, Previews in yellow, and external code in white.

### Flexible hybrid cloud infrastructure supporting HRA and HuBMAP data

Systematic integration of more than 50 open source algorithms developed by more than 30 teams is non trivial. Agreement on metadata and API calls is required to make the output of one algorithm compatible with the input expected by the next (set of) algorithms. Several algorithms crucial to tissue segmentation and annotation were developed by biologists with deep subject matter domain expertise but limited knowledge on how to build production pipelines. The HIVE production development team worked closely with the original algorithm authors to package their algorithms in a way so that they can be run reliably at scale in a hybrid cloud infrastructure that is flexible and extendable to meet evolving needs.

Specifically, the HIVE IEC, composed of members from the Pittsburgh Supercomputing Center (PSC), the University of Pittsburgh (Pitt), and Stanford University, implemented a flexible hybrid cloud infrastructure and community engagement supporting delivery of HuBMAP’s vision in the following key areas: **1) Curation and Ingestion:** Semi-automated data ingestion (https://software.docs.hubmapconsortium.org) currently from HuBMAP data providers, and–in the future–from community partners, and the general research community, to maximize efficiency and usefulness for building the HRA; **2) Integration:** Automated analysis and annotation of ingested data and alignment of these annotations to the HRA via the UBKG; **3) Findability and Accessibility:** Manifestation of backend resources in the modular architecture of APIs and containers, services, and documentation (https://software.docs.hubmapconsortium.org) that minimizes user friction in integrated searching, querying, analyzing, and viewing of HuBMAP data, and in the future of tissue maps at multiple spatial scales and among multiple layers of information; **4) Interoperability:** Use of the HuBMAP deployment of the UBKG with extensions to create the HuBMAP Ontology API (https://smart-api.info/ui/d10ff85265d8b749fbe3ad7b51d0bf0a) to translate HuBMAP data, HRA assets, and community data among one another via ontologies; the HuBMAP Ontology API contains endpoints for querying a UBKG instance with content from the HuBMAP context (https://ubkg.docs.xconsortia.org/contexts/#hubmapsennet-context). **5) Analysis:** Infrastructure support to currently enable users with interactive analyses of HuBMAP data via Jupyter notebooks, and in the future, batch workflows among both HuBMAP and user-contributed data and tools, including integration and mapping against the HRA; and **6) Sustainability:** HuBMAP’s flexible hybrid cloud infrastructure—efficiently leveraging on-premises resources at PSC for services that would incur significantly higher public cloud charges compared to on-prem, such as data storage, processing, analysis, and download (**Supplemental Figure 1** and **Methods**)—will facilitate sustainment of open tools, data, and infrastructure beyond the end of the HuBMAP program.

### Atlas construction and publication

HRA data comprises human-expert-generated data (e.g., ASCT+B Tables, OMAPs, AVRs, 2/3D reference objects), experimental data mapped to the HRA (via RUI location, HRA-aligned CTann, or OMAP/AVR), and enriched atlas data (e.g., HRApop and HRAlit); see **Fig. 1** for an overview of HRA digital object types and their crosswalks, **Box 1** for terminology, and **Methods** for details. HRA data, usage and extension of ontologies, unified data processing workflows, user interfaces, documentation, and instructional materials are detailed here.

#### Data types and status

The 6th release of the HRA v2.0 (December, 2023) includes an anatomical-structure systems graph which groups major organs into organ systems (e.g., digestive system, reproductive system); three ASCT+B tables that represent the *branching* structures for the blood and lymph and the peripheral nervous system; and 29 ASCT+B tables (**Fig. 1.a**) that document the nested *part_of* structure of other organs (e.g., kidney with cells that compose smaller and the subsequently large functional tissue units and organ parts) for a total of 33 ASCT+B tables. The cells that make up each of the anatomical structures are organized in a multi-level cell type typology, with ‘cell’ at the root and successively more specialized child nodes. Cells are mapped to five biomarker types: genes, proteins, metabolites, proteoforms, and lipids organized in a biomarker typology.

Anatomically based 3D Reference Objects (**Fig. 1.b**) in the HRA include the shape, size, location, and rotation of 1,215 3D anatomical structures with 519 unique ontology IDs in 65 organs (ASCT+B Tables list 4,869 anatomical structures). 2D References (**Fig. 1.c**) describe the spatial layout of 3,726 rendered 2D cells of 131 unique cell types for 22 FTUs in 10 organs. Labeled training data for spatial segmentation and machine learning models (**Fig. 1.d**) exist for 5 FTUs in 5 organs. 13 Organ Mapping Antibody Panels (OMAPs, **Fig. 1.e**) are linked to 197 Antibody Validation Reports (AVRs) and aligned with ASCT+B. Cell Type Annotation Tools (**Fig. 1.f**) include Azimuth and other references for healthy adult organs with crosswalks to cell types and biomarkers in the ASCT+B tables.

An important part of HRA processing is data enrichment. One example is HRApop (**Fig. 1.g**), which covers 553 tissue datasets that are used to compute cell type populations for 40 anatomical structures for which 3D reference objects exist, across 23 organs with 13 unique UBERON IDs. The code to reproduce the bar graph with HRApop data (seven datasets) is available^30^. HRAlit^31^ (**Fig. 1.h**) links HRA DOs to 7,103,180 publications, 583,117 authors, 896,680 funded projects, and 1,816 experimental datasets.

#### Data enrichment

This HRA processing step ensures that HRA DOs are high quality, usable, and useful for the user stories listed in **Table 1** and other applications. Normalization ensures that the raw data is well structured and presented in a format that can be readily translated into a knowledge graph via LinkML (https://linkml.io). During enrichment, certain implicit relationships are made explicit using OWL reasoning (e.g., transitive relationships like *subclass* and *part_of* are made explicit); external metadata is added from ontologies via APIs to enhance the graph’s usefulness (e.g., via queries to the scicrunch API to lookup antibody information for OMAPs); queries are used to add data from related graphs (e.g., extracting additional metadata and hierarchies related to anatomical structures, cell types, and biomarkers from popular biomedical ontologies like Uberon and Cell Ontology); and finalizes conversion from LinkML to knowledge graph (e.g., converting and combining all into an RDF- formatted graph in Turtle format).

#### Data publication

A new revised and extended version of the HRA digital objects (DOs) together with updated user interfaces and APIsare published every six months via the HRA Portal (https://humanatlas.io). The three HRA core ontologies (specimen, biological structure, and spatial ontologies)^7^ are shared as FAIR, versioned Linked Open Data (LOD) at https://lod.humanatlas.io. Select data is also provided in a relational database and as comma-separated value (CSV) files. RUI data is published via the HuBMAP, SenNet, GUDMAP, GTEx, and other portals. For instance, the HuBMAP Search API is queried by the HRA API to generate dataset graphs from HuBMAP data. The public graph with all donor, tissue block, tissue section, RUI data, and experimental dataset information can be accessed via the HRA dataset graph at https://lod.humanatlas.io/ds-graph.

The HRA Digital Object Processor (https://github.com/hubmapconsortium/hra-do-processor) supports automated processing of HRA data, including data normalization, validation, graph transformation, enrichment, and publishing. The end product is the HRA Knowledge Graph (KG, https://github.com/hubmapconsortium/hra-kg) and a set of flat files suitable for hosting all data as Linked Open Data. HRA infrastructure is optimized for deployment to Amazon S3, AWS AppRunner, and AWS CloudFront, but could be adapted to other file hosting platforms.

The HRA provenance graph keeps track of all HRA DOs (described using standard terminology from DCAT^32^ for organizing catalogs of data and W3C-Prov^33^ for describing the provenance of any particular piece of data) and code versions (via GitHub) so HRA KG provenance can be accessed and the HRA KG can be recomputed every six months.

**Supplemental Table 2** lists all data used in HuBMAP Data Portal (with blue background), HRA Portal (red), and demonstration Previews (yellow). Note that HRA data is mirrored by the European Bioinformatics Institute’s (EBI’s) Ontology Lookup Service (OLS), Stanford University’s NCBO BioPortal, and University of Michigan Medical School Ontobee. Publishing the HRA via widely used repositories for biomedical ontologies makes the HRA findable, accessible, interoperable, and reusable (FAIR)—users can browse the HRA data online or access it programmatically via APIs.

#### Usage and extension of ontologies

Data and workflows are linked to existing ontologies whenever possible, see **Table 2**. The 6th release of the HRA v2.0 uses biological structure ontologies Uberon 2023-10-27^9^ and FMA v5.0.0^10,11^ for anatomical structures; Cell Ontology (CL) v2023-10-19^12^ and PCL 2023-02-27^34^ for cell types; HGNC v2023-09-18^35^, Ensembl Release 111^36^, GeneCards Version 5.19: 15 January 2024^37,38^, and UniProt Release 2024_1^39^ for biomarkers. The Human Genome HGNC v2023-09-18 is used for the FTU Explorer. Spatial data is annotated using Dublin Core terms (DCTERMS) v2020-01-20^40^. Specimen data uses LOINC v2022-07-11 (v2022AB)^41^ for standardized representation of sex, race, and ethnicity data. Meta-ontologies such as DCTERMS and Relation Ontology^42^ (RO) are used to capture relationships among concepts within the HRA data. Assay type names come from BAO v2023-01-31^43^ and EFO v2023-02-15^44^. The use of these ontologies is strongly encouraged to maintain consistency among ASCT+B tables, Azimuth and other CTann tools, and OMAP data in support of atlas construction and usage.

**Table 2:** Ontologies used and extended.

| DO Type | Ontology Name/Version | #Terms added/ updated | #Links added/updated |
| --- | --- | --- | --- |
| Specimen |  |  |  |
|  | LOINC v2022-07-11 (v2022AB) |  |  |
| Biological structure |  |  |  |
| Anatomical structures | Uberon v2023-10-27 | 85 new terms added to Uberon through Feb 2024 | 333 relationships added to Uberon through Feb 2024<br>294 Uberon terms assigned 3D reference objects* |
| Cell types | CL v2023-10-19<br>PCL 2023-02-27 | 129** new terms added to CL through Feb 2024<br>468 new terms added to | 249 relationships added to CL through Feb 2024<br>468 relationships added to PCL |
|  |  | PCL through Feb 2024 | through Feb 2024 |
| Biomarkers | HGNC v2023-09-18<br>Ensembl Release 111<br>UniProt Release 2024_1<br>GeneCards Version 5.19: 15<br>January 2024 | None | None |
| Human Genome | HGNC v2023-09-18 for iFTU |  |  |
| Spatial |  |  |  |
|  | Dublin Core terms DCTERMS<br>v2020-01-20 |  |  |
|  | Relation Ontology used by<br>OBO |  |  |
| Assay Types |  |  |  |
|  | BAO v2023-01-31<br>EFO v2023-02-15 |  |  |

\* The foaf:depiction annotation is used to link 294 unique Uberon terms to 586 GLB files.
\*\* 25 out of 129 terms and 37 out of 249 links were added to CL supported by projects other than HuBMAP

A major contribution of the cross-consortium HRA effort is the extension of cross-species ontologies such as Uberon and CL to cover healthy human terms. Between 2021 and Oct 2023, 61 anatomical structure terms have been added to Uberon, 125 cell types were added to Cell Ontology. In January 2024, 468 cell types were added to the Provisional Cell Ontology (PCL), 461 for the human brain^45^—in support of HRA construction and usage. PCL uses computationally-derived marker genes from NS-Forest^46^ to define sc/snRNA-seq-derived cell types in the brain. The 461 human brain cell types were added to the ASCT+B tables. All PCL cell type terms are associated with biomarker genes using a “has_characterizing_marker_set” relation in the ontology. In the 6th release of the HRA, there are 962 anatomical structure terms that are either missing from Uberon or not yet crosswalked to Uberon terms in the ASCT+B table. The majority of missing terms are for blood and lymph vasculature, skeleton or skeletal muscle systems and are typically more specific than currently represented in Uberon (e.g., "dorsal branch of lateral proper palmar digital artery of fifth digit of hand"). Work is ongoing to improve mappings (∼100 mappings were recently added and will be published in 7th HRA release). 119 cell types are either unmapped or not yet in CL or PCL—an initial assessment suggests 60-70% of these are genuinely new terms for CL. These 387 biomarkers have Ensembl IDs or GeneCards IDs or have not been mapped rather than HGNC IDs—all of these terms have ASCTB-TEMP IDs. There are GitHub issues to add new terms to existing ontologies to properly represent data in the ASCT+B tables, including requests for 128 anatomical structures in Uberon. There now exists a formal operating procedure to include new cell types into CL via Minimal Information Reporting About a CelL (MIRACL) sheets^47^.

#### Unified processing workflows

The HRA SOPs^48^ detail the human expert and algorithmic steps needed to construct the HRA and to use it properly. Protocols published on protocols.io and other places are used to compile experimental data in a reproducible manner. In January 2024, there existed 235 HuBMAP protocols^49^—many of these document the reproducible workflows required to generate data used in HRA construction. **Figs. 1** and **2** provide an overview of the numerous steps required to construct the HRA and to map new experimental data to it.

**Figure 2:**
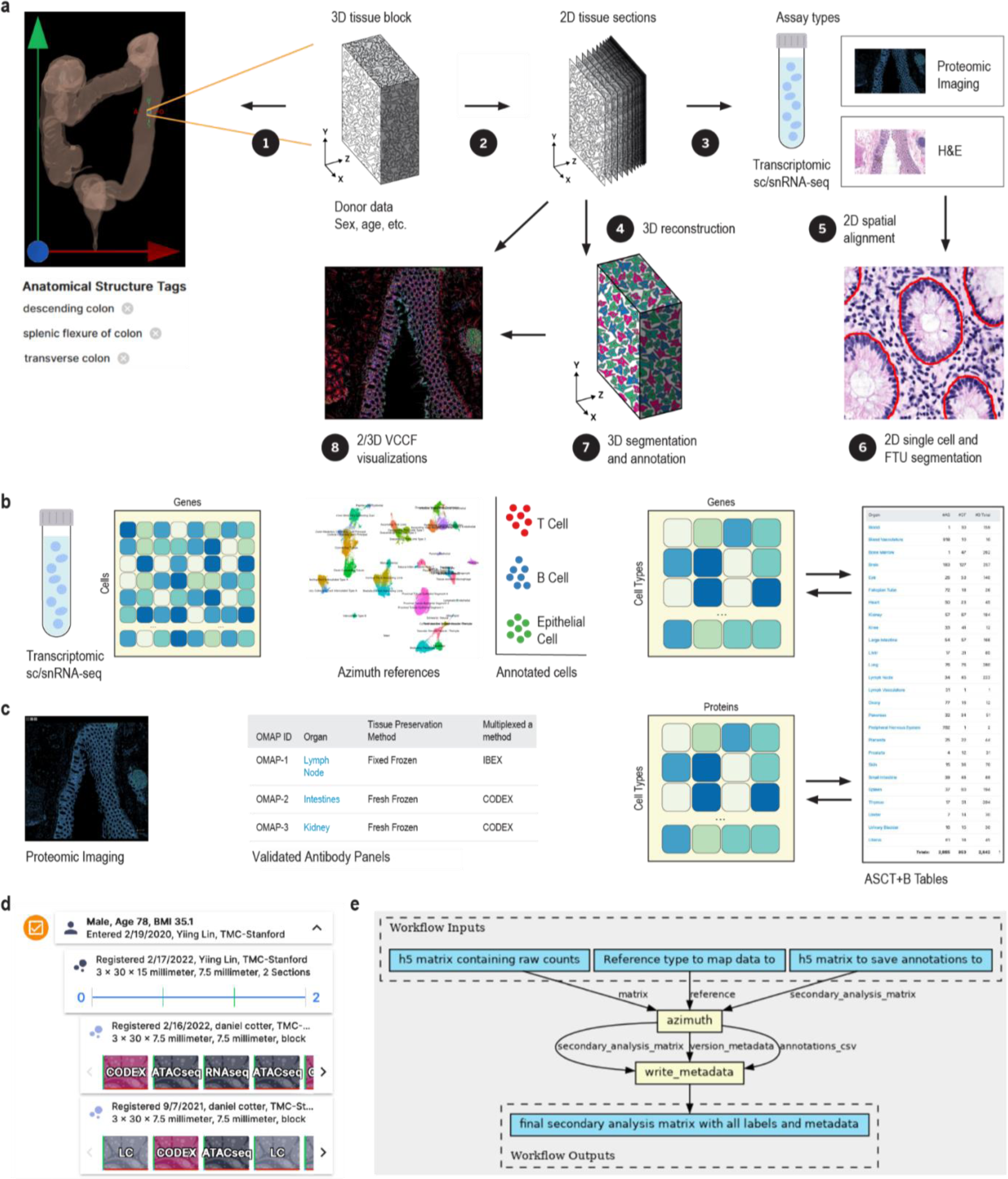
Mapping experimental data to the HRA. **a.** A tissue block is 3D spatially registered and semantically annotated using the Registration User Interface or the Millitome, see (1). A smaller part of the tissue block might be used for sc/snRNA-seq analysis (not shown) or cut into tissue sections (2). Tissue sections are analyzed using one or more assay types (3). Shown are single cell transcriptomics (e.g., sc/snRNA-seq), OMAP-aligned spatial proteomics (e.g., CODEX, Cell DIVE), and high resolution hematoxylin and eosin (H&E) stained histology images. Spatial alignment of different assay types for the very same or different tissue sections is non-trivial (5). H&E data is used to segment functional tissue units (FTUs) using trained machine learning models (6). 3D reconstruction of tissue volumes is accomplished by aligning data from multiple serial tissue sections computationally (4) followed by 3D segmentation and annotation (7). 2D or 3D data is analyzed to identify the distance of different cell types to the vasculature (VCCF Visualizations) as a multiscale common coordinate framework from which no other cell is very distant (8). **b.** Single cell/nucleus data (sc/snRNA-seq) is stored as a cell by gene matrix; cell types are annotated using Azimuth or other cell type annotation tools; results are aggregated to cell type by gene biomarker expression value matrices that are aligned with the ASCT+B tables; and are used in diverse HRA user interfaces (e.g., Exploration User Interface and FTU Explorer). **c.** OMAP-aligned spatial data generated using validated antibody panels linked to AVRs are analyzed to compute cell type by protein biomarker expression value matrices that are aligned with the ASCT+B tables using semi-automated workflows. **d.** The Exploration User Interface provides full provenance for donors (sex, age, BMI), data providers (upload date, contact name, affiliation), tissue blocks and sections (size, number, date, and contact info for RUI registration), and datasets (assay type) with links to raw data in the HuBMAP Data Portal, other data portals, or publications. **e.** CWL workflows detail what tools (yellow) are run on which input/output data (blue). Shown is the Azimuth cell type annotation workflow.

The HuBMAP Consortium has developed uniform computational processing pipelines for multiple data types: single-cell/single-nucleus RNA-seq, single-cell/single-nucleus ATAC-seq, multiplexed antibody-based spatial proteomics (CODEX [recently renamed to PhenoCycler] and Cell DIVE), multiplexed ion beam imaging (MIBI), Slide-seq, and Visium sequencing spatial transcriptomics, fluorescent *in situ* hybridization (FISH) spatial transcriptomics, among others. HuBMAP computational pipelines are all open source, published on GitHub as CWL workflows wrapping tools in Docker images (also executable via Singularity), with supplementary data (genome indexes/annotations, deep learning models) built into the published Docker images for portability and reproducibility.

The HuBMAP sc/snRNA-seq pipeline (https://github.com/hubmapconsortium/salmon-rnaseq, also used for sequencing spatial transcriptomics such as Slide-seq and Visium), is built on the Salmon quasi-mapping method^50^ and performs gene expression quantification for intronic and exonic sequences, with downstream analysis using Scanpy^51^ and RNA velocity computation via scVelo^52^. Outputs of the sc/snRNA-seq pipeline are annotated with an automated version of the Azimuth cell type annotation tool for supported tissues; these currently include heart, lung, and kidney, with additional annotations computed as new Azimuth references are integrated into HuBMAP processing infrastructure.

HuBMAP imaging pipelines (see **Methods**) are end-to-end analysis methods that accept raw images, perform illumination correction, background subtraction, and tile stitching if necessary, then perform cell and nucleus segmentation, writing expression and segmentation mask images as multichannel OME-TIFF files. The expression and mask images are further processed via Spatial Process & Relationship Modeling (SPRM)^53^, which computes image and segmentation quality metrics using the CellSegmentationEvaluator tool^54,55^, creates cell adjacency maps, computes features for each cell and nucleus, performs unsupervised clustering of cells, nuclei, and image pixels, computes biomarkers differentiating one cluster versus the rest for each clustering type, and writes results to CSV and HDF5 format for use by end users and in the HuBMAP Data Portal.

For HRApop (see **Fig. 1.g**), 445 public datasets from HuBMAP^2,56^, two datasets from SenNet^57^, 91 healthy datasets from two collections from CZ CELLxGENE^58,59^ (“Cells of the adult human heart” and “LungMAP — Human data from a broad age healthy donor group”), and 15 single cell datasets from GTEx^60,61^ were mapped to the HRA (see **Methods**). As a result, cell type population data exists for 40 anatomical structures in 23 organs with 13 unique UBERON IDs, separated by single cell transcriptomics (e.g., sc/snRNA-seq) and OMAP-aligned spatial proteomics (e.g., CODEX, Cell DIVE). Three organs (large intestine, small intestine, and skin) have cell type populations computed from transcriptomics and proteomics data.

For HRAlit^31^ (see **Fig. 1.h**), 583,117 experts, 7,103,180 publications, 896,680 funded projects, and 1,816 experimental datasets were mapped to the digital objects in the HRA (**Methods**).

#### User interfaces

The HuBMAP Portal (https://hubmapconsortium.org, see **Supplemental Figure 2**) introduces HuBMAP goals and links to experimental and atlas data, tools, and training materials. The HuBMAP Data Portal (https://portal.hubmapconsortium.org) supports ingest, search, exploration, and download of experimental data. The HRA Portal (https://humanatlas.io, see **Supplemental Figure 3**) supports the construction, access, exploration, usage, and download of HRA data.

The ASCT+B Reporter^3^ (https://humanatlas.io/asctb-reporter, see **Supplemental Figure 4**) supports the authoring and review of ASCT+B Tables and OMAPs by human organ experts. Detailed SOPs^48^ and video tutorials^62,63^ exist and more than 170 unique experts have contributed to the HRA as authors and/or reviewers using this tool as measured by the number of unique ORCID IDs listed in relevant digital objects of the 6th release HRA.

Azimuth^14^ (https://azimuth.hubmapconsortium.org, see **Supplemental Figure 5**) was developed by HuBMAP to automate the processing, analysis, and interpretation of single-cell/nucleus RNA-seq and ATAC-seq data. Its reference-based mapping pipeline reads a cell by gene matrix and performs normalization, visualization, cell annotation, and differential expression (biomarker discovery) analyses (see **Figs. 1.f** and **2.b**). Results can be explored within the app or downloaded for additional analysis. In HuBMAP, Azimuth is used in production mode to automatically annotate sc/snRNA-seq datasets. Crosswalks exist to associate Azimuth cell types to ASCT+B table terms and ontology IDs.

The Registration User Interface^64^ (https://apps.humanatlas.io/rui, see **Supplemental Figure 6** and SOP^65^) supports the registration of human tissue blocks into the 3D CCF with automatic assignment of anatomical structure annotations that are linked to the UBERON and FMA ontologies based on surface mesh-level collision events. The anatomical structure annotations in combination with ASCT+B table and experimental data make it possible to predict cell types that are commonly found in anatomical structures and colliding tissue blocks. RUI output in JSON format records registration data (e.g., tissue block UUID and 3D size, location, and rotation plus anatomical structure annotation based on bounding box) together with provenance data (e.g., operator name, date). The RUI is available as a stand-alone tool for anyone to use to contribute HRA-aligned spatial data. It is fully integrated in the HuBMAP, SenNet, and GUDMAP data ingest portals but requires authentication.

The Exploration User Interface^64^ (https://apps.humanatlas.io/eui, see **Supplemental Figure 7**) supports visual browsing of tissue samples and metadata at the whole body, organ, tissue, and cell levels, see **Table 1, US#3**. In January 2024, 901 human tissue blocks with 4,221 datasets from 351 donors and 19 consortia/studies were RUI-registered into the HRA 3D CCF. Users can filter by donor demographics (e.g., sex, age) or data source (e.g., consortium/study, technology). They can search for specific anatomical structures, cell types, or biomarkers to explore the number of tissue blocks that collide with an anatomical structure but also the cell types located in these anatomical structures or their characterizing biomarkers—according to the ASCT+B tables. Users can also run a 3D spatial search using an adjustable probing sphere, explore details on demand on the right with links to Vitessce^66,67^ visualizations in the HuBMAP Data Portal and links to data and tools in other data portals. The EUI with all HRA data is available as a stand-alone tool that supports exploration of all experimental data that has been mapped to the HRA. The EUI was customized, branded, and fully integrated in the HuBMAP, SenNet, and GTEx data portals to support exploration of consortia specific data, see **Supplemental Figure 8**.

Vitessce^66,67^ (http://vitessce.io) is a tool used to visually explore experimental data, Azimuth references (see **Supplemental Figure 5**), HRA segmentations and annotations, or cell-cell distance distribution visualizations (see **Supplemental Figure 9**), see previews in **Atlas usage** section.

The Interactive FTU Explorer^68^ (https://apps.humanatlas.io/ftu-explorer, see **Supplemental Figure 10**) supports the exploration of cell types in their 2D spatial context together with mean biomarker expression matrices, see **Table 1, US#4**. For example, tissue data (gene or protein expression levels, as available) can be compared against healthy HRA reference data to determine differences in the number of cells, cell types, or mean biomarker expression values to inform clinical decision making.

The HRA Organ Gallery^69,70^ (https://github.com/cns-iu/hra-organ-gallery-in-vr, see **Supplemental Figure 11**) supports the multi-scale exploration of 1,215 anatomical structures in the 65 3D Reference Objects of the HRA 2.0. Using a Meta Quest VR device, users select the male or female reference body; they can then select a specific organ and explore it with both hands. To achieve view update rates of 60 frames per second, lower level-of-detail models are used that were derived from the original HRA 3D Reference Objects.

The HRA API (https://humanatlas.io/api/, see **Supplemental Figure 12-14**) supports programmatic access to all HRA digital objects and the experimental HRApop data mapped into it. Users first select an API server and route, input query parameters, then view the query response.

The HRA Dashboard compares HRA, publication, and experimental data to world population data. **Supplemental Figure 15.a** shows population pyramids by age group of HRA survey respondents and tissue data donors in comparison to world population plus population pyramids by career age for HRA experts and publication authors. **Supplemental Figure 15.b** features the ethnic composition of survey respondents, HRA tissue donors, HRA experts, paper authors, world population in percentages. The choropleth map in **Supplemental Figure 15.c** shows the number of paper authors overlaid on a world map. CCF-HRA Data Dashboards help understand what HuBMAP data has been RUI registered (https://hubmapconsortium.github.io/hra-data-dashboard).

#### Documentation and instructional material

In January 2024, the HuBMAP Data Portal provides access to 8 publications and associated datasets, 50+ technical documents (https://software.docs.hubmapconsortium.org/technical), and links to 235 experimental protocols on protocols.io; the HRA Portal links to 20 Standard Operating Procedures (SOPs, https://zenodo.org/communities/hra) and to the Visible Human Massive Open Online Course (VHMOOC, https://expand.iu.edu/browse/sice/cns/courses/hubmap-visible-human-mooc) with 39 videos, 4 self-tests and 3 quizzes, 2 hands-on tutorials, plus entrance and exit surveys (see **Supplemental Figure 16**).

### Previews of Atlas Usage

Two exemplary previews demonstrate the usage of atlas data and code developed in HuBMAP for gaining insights into pathology, see user stories that drive HRA construction and usage (**Table 1**). All data and code are publicly available on GitHub^71,72^ and Dryad^73^. The cell distance distribution code is available via the HRA Portal^74^ in support of **Table 1, US#5**. Cell type annotations for the CODEX multiplexed imaging dataset of the human intestine are published via DRYAD at https://doi.org/10.5061/dryad.pk0p2ngrf. Full data and code integration into the HuBMAP Data Portal workflows are planned for future releases.

### Perivascular immune cells in lung

Normal lung function depends on careful matching of airflow to blood flow to achieve normal gas exchange. The abnormal presence and activity of immune cells results in leaky vascular membranes and edema that thickens the gas exchanging membrane, and accumulation of mucus and cellular debris in the airspace can cause a mismatch between flow of air and blood. Persistent inflammation results ultimately in fibrosis. Prior work using scRNA-seq data and the CellTypist common reference dataset discovered previously under- appreciated organ-specific features and aggregates of T cells and B cells^75^. Recent publications in the field of mucosal immunology illustrate the segregation of immune cells within aggregates in human lung tissue and their role in abnormal regulation of vascular function^76,77,78,79^. Molecular and cellular changes, including fibrotic and immune-cell rich regions were recently imaged in the lungs of children with bronchopulmonary dysplasia (BPD), a chronic lung disease following premature birth^80^. The Vitessce tool is applied in **Fig. 3.a** to comparatively visualize and quantify cellularity of specific regions of healthy adult and BPD lung to demonstrate an assessment of multiple cell types relative to nearest vascular endothelial cell nuclei using single-cell spatial protein biomarkers.

**Figure 3:**
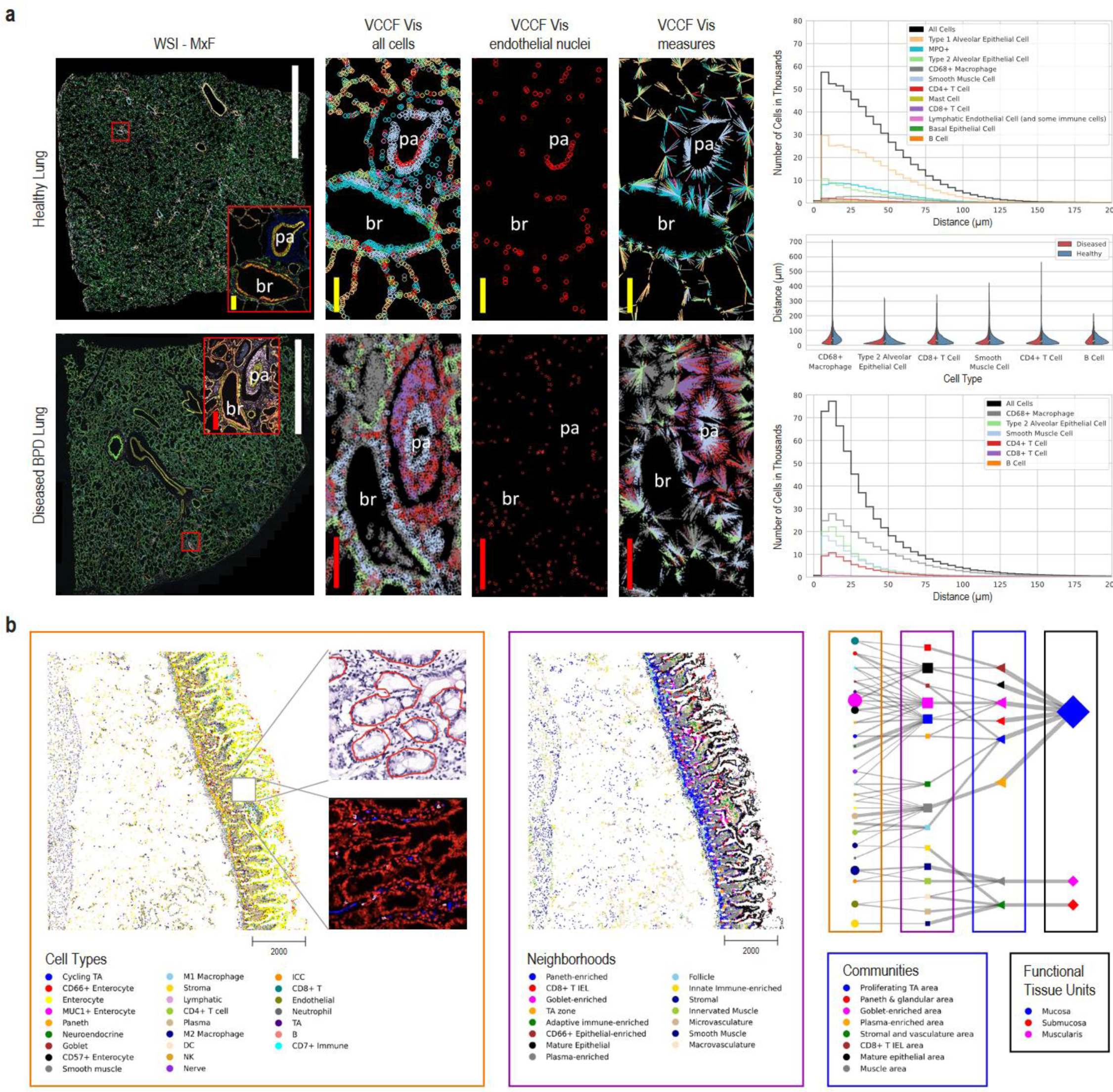
Human Reference Atlas Usage. **a.** HRA can be used to compare the distribution of parenchymal cells including endothelial, epithelial, and muscle that compose the blood vessels, airways and gas exchanging functional lung structures, and resident immune cells including macrophages, to local vasculature (VCCF Visualizations) in healthy (top) and diseased (bottom) lung using multiplexed immunofluorescence microscopy images with bronchiole (br) and an accompanying small pulmonary artery (pa). Scale bar legend: white: 5 mm, red: 200 µm, yellow: 100 µm. The graphs on the right show distance distributions for cell types present in the healthy lung (top) and diseased BPD lung (bottom); the violin plot (middle) shows a comparison between distance distributions for cell types common in both datasets. Datasets are on GitHub^81^. **b.** Multi-level cell neighborhoods can be computed to analyze and communicate the structure and function of FTUs; tissue image with cell type annotations and zoom into H&E with FTU segmentations (red outlines) and zoom into the multiplexed image (CODEX) is shown in left, neighborhoods are given in the middle; hierarchy of FTUs, neighborhoods, communities, and cell types are shown on the right. Datasets are on GitHub^81^.

Demonstrated are whole slide photomicroscopy images of PhenoCycler^R^ multiplexed immunofluorescence assays (WSI-MxF) applied to examples of healthy lung tissue (top row, 28 antibody panel) and BPD tissue (bottom row, 25 antibody panel). Digital zoom is used to highlight similar regions of interest (red box, MxF-ROI) focused on a bronchiole (br) and an accompanying small pulmonary artery (pa). Immune cell aggregates, primarily CD3+ lymphocytes, are noted around both structures in the BPD lung. To assess a vascular common coordinate framework (VCCF) for the localization of the immune and other lung cells, the cell types are masked in cell specific colors (see key) and distances measured to nearest endothelial cell nucleus (red circles). The cell-to-nearest endothelial cell measurements are seen as spokes colored by the cell type. In brief, the graphical representations quantitatively demonstrate increased cellularity with a predominance of CD4+, but not CD8+, lymphocytes, as well as myeloid immune cells, positioned in close proximity to the lung vasculature in the diseased lung. The VCCF visualizations suggest endothelial cells embedded within the lymphocytic aggregates that congregate around the pulmonary artery (pa) in the diseased as compared to the healthy lung. Analyses of cell populations (number of cells per cell type and their mean biomarker expression values), as well as cell to cell and cell to functional tissue unit spatial distribution patterns are valuable to understand tissue and cellular disruptions that account for organ failure in lung disease. In this example, the diseased tissue has gas exchange membranes thickened in part by extravascular immune cell aggregates such that distribution of cell distance to nearest endothelial cells is compressed and exaggerated (see graphs in right of **Fig. 3.a**).

Measurements on 2D images, as demonstrated, provide novel insights however cell segmentation and determination of relative locations in the lung is particularly challenging given the complex airway and vascular branching system and the very thin, highly multicellular gas exchange membranes of the alveoli. It is anticipated that application of similar segmentation, cell-cell and cell-FTU measures to 3D lung tissue volumes will identify currently under-appreciated relationships in health and disease, see **Table 1, US#5**. HuBMAP code can be used on human lung tissue to rapidly understand how the spatial organization of specific immune cell infiltrates relate to disease pathophysiology revealing potentials for targeted therapy to ameliorate human disease. Code and data are available via https://cns-iu.github.io/hra-construction-usage-supporting-information.

#### Hierarchical cell type populations within FTUs

Functional tissue unit segmentation algorithms for histology data^82,83^ and hierarchical cell community analysis^73^ for paired spatial data (see **Methods** for both) can be combined to analyze and communicate the structure and function of FTUs across scales. FTU segmentation algorithms are run as part of the standard HuBMAP workflows (currently limited to glomeruli in kidneys, soon to be expanded to crypts in the large intestine and white pulp in spleen).

Exemplarily, we feature an example hierarchical cell neighborhood analysis previously developed for analyzing cell type neighborhoods across scales and applied within the healthy human intestine^73^ (see Jupyter Notebook on Github^71^). We have named some of these scales: cellular “neighborhoods”, “communities”, and “functional tissue units.” The calculation of similar cellular neighborhoods, communities, and tissue units across different scales is analogous to how we might think that people form neighborhoods, cities, and states.

Currently, the HuBMAP Data Portal supports cell segmentation of antibody-based multiplexed imaging data but lacks the ability to annotate the cell types for such datasets. This functionality is under active development (see **Methods**). Consequently, to demonstrate this user story, a separately processed version of the intestine data^84^ (containing cell type annotations) was used. The cell type predictions for the same dataset, using the current development version of the cell type model by the van Valen lab (see **Methods**) are also made available via https://cns-iu.github.io/hra-construction-usage-supporting-information. This version of the model was trained on several multiplexed datasets spanning tissue types and multiplexed imaging modalities. Cell segmentation masks, generated using Mesmer, are also included with the prediction data.

The Jupyter Notebook on GitHub^71^ demonstrates how to read a previously published CODEX multiplexed imaging dataset of the human intestine^73^, identify how cell types correspond to larger multicellular structures, and support exploration of the relationships between these higher order cellular neighborhoods. By visualizing the data in tissue coordinates, we can observe potential layering or consistent functional tissue unit (FTU) structures, such as the repeat structure of the intestinal crypt in the proximal jejunum in the small intestine (see **Fig. 3.b**, left). Furthermore, we can quantify these relationships across different scales of cellular neighborhoods and represent them as a network graph (see **Fig. 3.b**, right) in which line thickness indicates the percentage of cells in the next level. Note that the tissue samples and the graph use the very same node color coding and naming.

### Usage Statistics

Between January 2021 and December 2023, 87,310 unique users visited the HuBMAP Data Portal. These users visited 382,384 pages; the top-5 most frequent referrers are nature.com, hubmapconsortium.org, humancellatlas.org, azimuth.hubmapconsortium.org, and humanatlas.io. Azimuth supported the upload and cell type annotation of 27,000 datasets with more than 366,000,000 cells. Since Jan 2023, the new HRA Portal served 5,700 unique visitors accessing 89 pages; the top-5 referrers are the HuBMAP Data Portal, nature.com, google.com, gtexportal.org, and github.com. The 3D reference objects were accessed 3,065 times via the NIH3D website. The HRA OWL file was accessed 1,325 times via the NCBI BioPortal Ontology Browser (https://bioportal.bioontology.org/ontologies/CCF) and 11,531 times via the EBI OLS Ontology Browser (https://www.ebi.ac.uk/ols/ontologies/ccf). 310 students registered for the VHMOOC and spent 5,652 hours reviewing materials, taking self tests, and engaging in a community of practice.

## Discussion

This Resource paper describes data, code, and tools that are of broad utility, interest, and significance to the construction and usage of a multiscale human reference atlas. The HRA effort and evolving data and code infrastructure is novel and unique in several ways: (1) The HRA integrates many assay types across scales, from whole human body to single cell level. (2) It provides standard operating procedures and tools to spatially and semantically register human tissue from 65 organs into one common coordinate framework. (3) It links anatomical structures, cell types, and biomarkers to ontologies and extends existing ontologies when needed. (4) The HRA comes with diverse interfaces that allow users to explore and inspect diverse HRA Digital Objects (3D Reference Objects, ASCT+B Tables, OMAPs, etc.), experimental data, and documentation from the participating consortia, as well as HuBMAP data in particular (HRA Portal, ASCT+B Reporter, RUI, EUI, Cell-Cell Distance Distribution Visualizations, Interactive FTU Explorer, and HRA Organ Gallery in VR). For each user interface, we provide a **Supplemental Figure (3, 4, 6, 7, 9, 10, 11)** with high-resolution screenshots and detailed annotations. (5) HRA development is community driven and collaborative; monthly working group meetings inform strategic decision making; 50+ open source algorithms developed by 30+ teams have been systematically integrated into a flexible and adaptive system architecture that adds value to many atlasing projects; new HRA data and code are publicly released every six months via the HRA Portal and ontology services. The resulting HRA is a multiscale, multimodal, three-dimensional digital product that unifies biomedical knowledge across organs, anatomical structures scales, demographic markers, assay types, and links them to ontologies and makes human reference data computable.

The 6th release HRA has several known limitations that will be addressed in future iterations. Starting with the 8th release (published in mid December 2024), all HRA DOs and their complete provenance are covered in the HRA KG. Cell states are not currently captured in CL nor are ever specific cell types emerging from single cell technologies; however, the HRA started to use cell type annotation that have the format “CL ontology term:cell state or specificity terminology” (e.g., pancreatic stellate cell:quiescent mapped to pancreatic stellate cell CL:0002410 which is in CL or enterocyte:MUC1 positive mapped to enterocyte CL:0000584) along with a confidence term to the CL match (e.g., skos:narrowMatch (for cell states or new cell types) or skos:exactMatch (those that exactly match a cell type in CL) to allow it to be represented in the HRA KG and UBKG and updated at a later date when the computational community has settled on a method to ontologically represent such cells. Terms in the HRA KG will be given a ASCTB-TEMP ID and individual cells annotated by a cell type annotation model will be given a cell type ID to facilitate future updates when they become available.

In addition, existing workflows for mapping new experimental data to the HRA will be expanded in three main ways: (1) HuBMAP plans to add several new Azimuth references (e.g., for large and small intestine) and update existing references (e.g., for kidney and lung) to capture new data with additional/revised cell type annotations and CL terms plus CL IDs via crosswalks. (2) Eight new OMAPs were published in the 7th HRA release and several more are in progress for the 8th HRA release, substantially increasing the number of spatial datasets that can be mapped to the HRA. (3) Starting with the 8th HRA release, new 3D organs will be added to the RUI: Quadriceps femoris and triceps surae skeletal muscles, esophagus, and lymphatic vasculature.

Currently, the HRA knowledge graph and API drive different 2D and 3D user interfaces in the HuBMAP, SenNet, GUDMAP, GTEx data portals, and the CZ CellGuide. In line with US#6 (see **Table 1**), we will develop additional lightweight web components that make it easy to access HRA data and feature HRA functionality in other websites. Plus, we are implementing diverse HRA dashboards (US#7) to communicate what HRA DOs exist; what experimental data is used for HRA construction (full provenance); how existing ontologies are expanded to capture healthy human terms and linkages; who is using HRA data, tools and APIs; and how representative the atlas is.

Last but not least, we will expand the interlinking of the HuBMAP Data Portal and the HRA Portal. Specifically, we will ingest new releases of the HRA into the HuBMAP UBKG so that anatomical structures, cell types, and biomarkers supported in the HuBMAP Data Portal are aligned with existing ontologies and the 3D spatial reference framework. As HuBMAP teams start to compile 3D datasets, there is a need to compare existing algorithms for spatial alignment of multiple subsequent tissue sections in support of 3D tissue block reconstruction, as was done for 2D cell segmentation methods^54,55^. This is expected to considerably improve HRA construction and usage.

Community input to HRA user stories, data, code, user interfaces, and training materials is welcome and experts interested to learn more about or contribute to the HRA effort are encouraged to register for the monthly working group events online^85^.

## Methods

Human expert generated data and experimental tissue data are used to construct the HRA, see **Fig. 1**. New experimental data is mapped to the HRA via (1) 3D spatial registration, (2) using suspension based (e.g., sc/snRNA-seq), or (3) spatial (e.g., CODEX^86^, Cell DIVE^87^, IBEX^88,89^, Imaging Mass Cytometry^90^, and other multiplexed, antibody-based protein imaging platforms) assay types that are aligned with the HRA, see **Fig. 2**.

### Expert generated data

#### ASCT+B tables

Anatomical structure, cell type, and biomarker (ASCT+B) tables (https://humanatlas.io/asctb-tables, **Fig. 1.a**) are compiled by experts using the ASCT+B Reporter User Interface (see **Supplemental Figure 4**) following SOPs^91^. Note that the brain ASCT+B table is unique in that it was computationally derived using the common cell type nomenclature approach^92^ which chains together critical cell type features (e.g., brain region, cortical layer), broad cell type class, and gene biomarker information into the annotation.

Starting with the 6th release of the HRA, new and revised tables list cell type parents present in Cell Ontology (CL) for the about 600 cell types that currently have ASCTB-TEMP IDs (i.e., temporary ontology terms and IDs) as they do not yet exist but are systematically added into CL via the HRA effort. This makes it possible to show the complete cell typology in CellGuide (https://cellxgene.cziscience.com/cellguide) and other tools. For example, **Supplemental Figure 17** depicts a CZ CellGuide visualization for Neuron (CL:0000540) showing the CL ontology typology with the ‘neuron’ cell type highlighted in green together with its parent (‘neural cell’, which is_a ‘somatic cell’, which is_a ‘animal cell’) and children nodes (e.g., ‘GABAergic neuron’ and ‘glutamatergic neuron’). The interactive visualization is at https://cellxgene.cziscience.com/cellguide/CL_0000540.

A special focus within HuBMAP has been the development of a detailed blood vasculature ASCT+B table in support of a Vasculature Common Coordinate Framework (VCCF)^93–95^ (https://humanatlas.io/vccf). Relevant data captured in the VCCF include blood vessels and their branching relationships, as well as associated cell types and biomarkers, the vessel type, anastomoses, portal systems, microvasculature, functional tissue units, links to 3D reference objects, vessel geometries (length, diameter), and mappings to anatomical structures the vessels supply or drain.

### 2D and 3D reference objects

Professional medical illustrators follow SOPs^96,97^ to generate 2D reference FTU illustrations and 3D reference anatomical structures, see **Fig. 1.b, c**. Most 3D reference organs were modeled using the male and female datasets from the Visible Human Project provided by the National Library of Medicine^3^.

The ASCT+B Tables in the 6th HRA release feature ontology-aligned terminology for 4,499 unique anatomical structures and 1,195 cell types. For some of these anatomical structures and cell types there exist anatomically-aligned, spatially explicit reference objects. Specifically, there exist two-dimensional illustrations of 22 FTUs in 10 organs with 3,726 cells of 131 cell types plus three-dimensional reference objects for 1,215 anatomical structures with 519 unique Uberon IDs in 65 3D reference objects (male and female, left and right organs) with 36 unique Uberon IDs. A crosswalk associates each of the 2/3D anatomical structures and cell types with their corresponding terms in the ASCT+B tables, see SOP^98^.

### Segmentation masks

Different tools are used to support manual segmentation of images by human experts, i.e., to assign each pixel in an image to an object such as a single cell, FTU, or anatomical structure. Within the HRA effort, the QuPath^99^ tool is used by organ experts to generate 2D segmentation masks for FTUs and vasculature (see SOP^100^) and DeepCell^101^ Label (https://label.deepcell.org) is used to get 2D segmentation masks for single cells. Resulting “gold standard” segmentation and annotation data (see **Fig. 1.d**) is needed to train machine learning algorithms so that experimental datasets can be automatically segmented, see **Fig. 2.a, 6**.

#### OMAPs and AVRs

Organ Mapping Antibody Panels (OMAPs, https://humanatlas.io/omap) are collections of antibodies designed for a particular sample preservation method and multiplexed imaging technology to allow spatial mapping of the anatomical structures and cell types present in the tissues for which they were validated^15,102^ (see **Fig. 1.e**). OMAPs are wet bench validated antibodies which experts initially identify as candidates for their multiplexed antibody based imaging experiments by using literature, available antibody search engines, and potentially also the ASCT+B Reporter User Interface (see **Supplemental Figure 4** and SOP^103^). Antibodies in OMAPs link to expert generated HuBMAP Antibody Validation Reports (AVRs, https://avr.hubmapconsortium.org and SOP^104^) that provide details on the characterization of individual antibodies for multiplexed antibody-based imaging assays. Antibody validation is expensive and time consuming, so these resources are designed to jump start other researchers to be successful and reduce the time required for multiplexed antibody based imaging studies.

#### Cell annotation references

A large majority of single cell data is single cell or single nucleus RNA-seq data. Cell type annotation tools (see **Fig. 1.f**) such as Azimuth^14^, CellTypist^75,105^, and Popular Vote (popV)^106^ are commonly used to cluster cells based on their gene expression profiles, followed by assigning those UMAP (Uniform Manifold Approximation and Projection)^107^ clusters to cell types based on published gene expression profiles. **Supplemental Table 1** shows the number of cell types that these three tools can assign per organ (rightmost columns)—compared to the number of cell types in the ASCT+B tables and 3D reference object library (middle columns); the second column shows the number of datasets available via the HuBMAP, SenNet, GTEx, and CZ CELLxGENE data portals. Note that there exist no datasets for some organs (e.g., urinary bladder).

Human expertise is required to compile crosswalks that associate cell labels assigned by these three tools to terms in CL. Mapping cell labels to CL can be partially automated; however, this is more effective if the labels researchers provide are written out rather than listed as abbreviations since different research groups do not use standardized abbreviations for cell types. Automated mapping to CL is further hindered when the cell type is not yet present in CL, in this case, often a parent cell type is used as a placeholder until the exact cell type can be added to the ontology. For these reasons, it is desirable to construct crosswalks that use the most specific cell type supported by experimental data. Depending upon the number of active editors/curators available for adding the new cell types that single cell RNA sequencing is discovering, prioritization of new terms and collecting supporting literature takes time. Resulting crosswalks are organ specific and they are published as cell type annotation specific crosswalks that associate any cell type assigned by the three tools with the corresponding term in the ASCT+B tables, see examples^108^.

### Experimental data

The HuBMAP Data Portal (https://portal.hubmapconsortium.org) uses a microservices architecture (see **Supplemental Figure 1**) to serve data and code via a hybrid on-premises and cloud approach using federated identity management, Universal Unique Identifiers (UUIDs), and full provenance for data management plus data security. Workflow and container support exist for diverse unified analysis pipelines and interactive exploration tools. This architecture makes it possible to ingest data at scale, adjust metadata formats as needed, add new algorithms and workflows as they become available, and ensure production phase speed and scalability for all services. On January 20, 2024, the HuBMAP Data Portal provided open access to 2,332 datasets from 213 donors. 360 of these datasets are sc/snRNA-seq and 79 are spatial OMAP-aligned datasets.

#### Tissue collection and RUI registration

The Registration User Interface^64^ (RUI, https://apps.humanatlas.io/rui) was implemented to support the spatial registration of tissue blocks into the HRA CCF, see **Supplemental Figure 6**. It collects sample ID, donor metadata, plus provenance information (who registered the data and when) in the process. Subject matter experts with knowledge of the spatial and donor data for the tissue samples use the RUI to register their tissue samples—supported by a designated HRA registration coordinator as needed, see SOP^65^. Alternatively, a more collaborative workflow is available in which the registration coordinator plays a more active role in making the registration with guidance from a subject matter expert. These workflows are explained in more detail in two dedicated SOPs detailing how the RUI can be used^65^ and the responsibilities of the registration coordinator^109^. Next, the registration coordinator uses a location processor tool^110^ to combine tissue sample metadata with de-identified donor metadata (sex, age, body-mass index, race, etc.) and publication metadata (DOI, authors, publication year, etc.). Once the samples are registered and the metadata has been enriched, the registration coordinator contacts the subject matter expert to check accuracy and completeness. The registration coordinator then publishes the validated registration set, making it accessible through the Exploration User Interface (EUI, https://apps.humanatlas.io/eui), see **Supplemental Figure 7**.

Tissue block registration can be streamlined and made more reproducible through the use of the "millitome," (https://humanatlas.io/millitome) a device that aids wet bench scientists to cut and register multiple tissue blocks from a single organ in a reproducible manner. This 3D-printable apparatus is designed to secure a freshly procured organ and is fitted with cutting grooves that can direct a carbon steel cutting knife for uniform slicing, see HRA’s millitome catalog (https://hubmapconsortium.github.io/hra-millitome) to access and customize organ millitomes based on donor sex, organ laterality, organ size, and cutting intervals. Each millitome package contains an STL file for 3D printing the millitome’s reproducible surface geometry plus a lookup sheet correlating millitome locations with tissue sample IDs assigned by the research team. After slicing the organ using the millitome, scientists document the samples on the lookup sheet and submit this data for review by the HRA millitome facilitator. Once the package is complete, data is added to the EUI for review by scientists to verify registration accuracy in terms of tissue size, placement, and orientation. SOPs detail millitome construction^111^ and usage^112^.

#### Single cell/nucleus RNA-seq transcriptomic data annotation

Single cell/nucleus RNA-seq transcriptomic datasets are downloaded from four data portals using the hra- workflows-runner (https://github.com/hubmapconsortium/hra-workflows-runner). For data from HuBMAP and SenNet (each dataset comes from exactly one donor), Search APIs (HuBMAP: https://search.api.hubmapconsortium.org/v3, SenNet: https://search.api.sennetconsortium.org) are used to obtain a list of dataset IDs for all existing cell-by-gene matrices in H5AD format and to download these files plus donor metadata. For GTEx, a single H5AD file is downloaded from https://gtexportal.org/home/singleCellOverviewPage. For CZ CELLxGENE, datasets are stored in collections, and one collection can contain multiple datasets and donors; the workflow runner reads in an index of all healthy adult human collections compiled using the CZI Science CELLxGENE Python API (https://chanzuckerberg.github.io/cellxgene-census/python-api.html); it splits the collection into unique donor- dataset pairs; and runs all H5AD files through the three cell type annotation tools: Azimuth^14^, CellTypist^75^, and popV^113^, see **Supplemental Table 1**. Azimuth (https://azimuth.hubmapconsortium.org) serves organ-specific human adult references for 10 unique organs (lung and tonsil have a revised v2 which is used here); for Azimuth, there exists HRA crosswalks^108^ for 194 unique cell types in 7 organs (3D spatial reference organs do not exist for blood, adipose tissue, and bone marrow). For CellTypist (https://www.celltypist.org), there exists crosswalks for 13 organs and a total of 214 unique cell types. For popV (https://github.com/YosefLab/PopV), we provide crosswalks for 22 organs and 134 unique cell types. There are 542 unique cell types across all three tools. The workflow runner outputs four files: (1) cell summaries for all sc-transcriptomics datasets, subset by cell type annotation tool and (2) a corresponding metadata file with donor and publication information; (3) cell summaries for all sc-proteomics datasets and (4) a corresponding metadata file with donor and publication information^114^. All four files are used during the enrichment phase to construct the atlas-level HRApop data.

#### Cell and FTU segmentation for spatial data

Antibody-based multiplexed imaging datasets, once uploaded to the HuBMAP Data Portal via the Ingest portal, are processed using a unified CWL workflow for cell and nuclei segmentation. Whole cell segmentation for CODEX datasets (https://github.com/hubmapconsortium/codex-pipeline) is done using Cytokit^115^, and Cell DIVE (https://github.com/hubmapconsortium/celldive-pipeline) and MIBI (https://github.com/hubmapconsortium/mibi-pipeline) datasets are processed using Deepcell’s Mesmer model^116^. Resulting cell segmentations are assigned a segmentation quality score using CellSegmentationEvaluator^54^. Cell segmentation for forthcoming 3D spatial proteomics datasets is provided by 3DCellComposer^117^ in combination with trained 2D segmenters.

FTU segmentation on PAS/H&E stained histology datasets is done using code developed via two Kaggle competitions^82,83^. The current production pipeline includes support for FTUs in the kidney, with large intestine and spleen that will be run when histology datasets become available.

#### Cell type annotation for spatial proteomic data

After cell segmentation, spatial cell type annotation is performed using the antibody metadata for marker channels for CODEX datasets, soon expanding to MIBI and Cell DIVE. OMAPs link marker panels in the datasets to cell types in the ASCT+B tables. The SPRM package (https://github.com/hubmapconsortium/sprm) computes various statistical analyses, including mean marker expression for all cells. The Van Valen lab is working on a foundation model to classify cell types across tissue types and imaging technologies; this model will cover 30+ cell types and new cell types will be added as new multiplexed imaging data become available. Since the model is under active development, it has not been released publicly yet, but is available as a private GitHub repository to HuBMAP members (https://github.com/hubmapconsortium/deepcelltypes-hubmap). In addition to this model, various teams have been annotating cell types with different approaches such as manual labeling with clustering or graph-based networks such as STELLAR^118^. The intestine datasets by Hickey et al.^118,119^ were annotated using a combination of manual and STELLAR approaches.

#### Spatial alignment for multi-omics data

Spatial structural alignment of different segmentation masks, see **Fig. 2.a, 5**, in support of multi-omics assay data analysis and/or alignment of spatial transcriptomics data to H&E imaging data can be performed using STalign^120^. Segmented cellular spatial positions are rasterized into an image representation to be aligned with structurally matched H&E images. Because tissues may be rotated, stretched, or otherwise warped during data collection, both affine and diffeomorphic alignments are performed. Such an alignment is achieved by optimizing an objective function that seeks to minimize the image intensity differences between a target (rasterized cell positions) and source (H&E) image subject to regularization penalties. The resulting learned transformation is applied to the original segmented cellular spatial positions to move the points into an aligned coordinate space. Such 2D spatial alignment facilitates downstream molecular and cell-type compositional comparisons within matched structures as well as integration across technologies.

#### Spatial data 3D reconstruction

Spatial alignment of multiple subsequent tissue sections in support of 3D tissue block reconstruction (see **Fig. 2.a, 4**) has been performed using MATRICS-A^121^ for skin data. Additional tools for 3D tissue block reconstruction have been developed and include SectionAligner, 3DCellComposer^117^, and CellSegmentationEvaluator. SectionAligner takes as input a series of images of 2D tissue sections, segments each piece of tissue in each section, and aligns the slices of each piece into a 3D image. 3DCellComposer uses one of various trained 2D cell segmenters (such as Mesmer), to segment each 3D image into individual cells using CellSegmentationEvaluator to automatically optimize parameter settings.

### Atlas enriched data

#### Mesh-Level collision detection

Extraction sites are post-processed via code specifically developed for efficient spatial registration using mesh surfaces^122^. To improve performance during tissue registration, the RUI uses bounding-box collision detection to determine (approximate but fast) intersections at runtime. To optimize accuracy, surface mesh collision detection is used during the enrichment phase to determine exact intersection volumes between a given RUI location and any anatomical structures it intersects with based on mesh-level colliders. The “3D Geometry-Based Tissue Block Annotation: Collision Detection between Tissue Blocks and Anatomical Structures” code is available on GitHub^123^ and the API is deployed to AWS^124^.

#### HRAlit

The HRA digital objects from the 6th release (36 organs with 4,499 anatomical structures, 1,295 cell types, and 2,098 biomarkers) were linked to 7,103,180 publications, which are associated with 583,117 authors, 896,680 funded projects, and 1,816 experimental datasets^31^. The resulting HRAlit database represents 21,704,001 records as a network with 8,694,233 nodes and 14,096,735 links. It has been mined to identify leading experts, major papers, funding trends, or alignment with existing ontologies in support of systematic HRA construction and usage. All data and code is at https://github.com/cns-iu/hra-literature.

#### HRApop

HRA cell type populations (HRApop) provide experimental data evidence for the existence of specific cell types and biomarkers (e.g., gene mean expression values) in the anatomical structures for which 3D reference models exist. In the 6th release HRA, there are 1,215 anatomical structures of 519 types (i.e., unique UBERON IDs) for 65 organs (including male/female, left/right).

There are three criteria that experimental datasets have to meet to be used in HRApop construction: (1) they must be spatially registered using the RUI; (2) have cell population data (e.g., a H5AD file that can be annotated via CTann tools [see **Supplemental Table 1**] or via proteomics workflows); (3) are from a data portal with quality assurance/quality control (QA/QC) or are published in a peer-reviewed paper.

To construct HRApop v0.10.2, we downloaded 9,613 datasets from four data portals (including 5,118 H5AD single cell transcriptomics datasets see **Supplemental Table 1** and 74 single cell proteomics datasets from HuBMAP published in two papers^73,121^). Exactly 553 datasets (371 sc/snRNA-seq transcriptomics and 74 spatial proteomics from HuBMAP, two from SenNet, 91 from CZ CELLxGENE, and 15 from GTEx) satisfied the three requirements for HRApop construction and were used to compute the reference atlas.

HRApop v0.10.2 is used in US#1-2 (see **Atlas Usage**) to make predictions for “test” datasets for which either a RUI-registered extraction site or a cell type population exist. Specifically, we can use HRApop atlas-level data to predict cell type annotation or spatial origin for 2,004 HuBMAP, 166 SenNet, 4,789 CZ CELLxGENE, and 50 GTEx datasets.

Out of the 9,060 datasets that did not satisfy the requirements, 2,065 datasets have a 3D extraction site but no cell summary, 4,598 datasets have a cell summary but no extraction site, and 2,397 datasets have neither. A total of 17 extraction sites have been registered with the RUI but have no mesh collisions with any anatomical structure; we are working with the experimental teams that performed the registration to rectify this.

HRApop reproducible workflows are at https://github.com/x-atlas-consortia/hra-pop and https://github.com/hubmapconsortium/hra-workflows-runner. HRApop data is published as a versioned, HRA- enriched dataset graph, see https://purl.humanatlas.io/graph/hra-pop.

#### VCCF distances & Vitessce visualizations in 2D

In support of constructing a vasculature based CCF (also called VCCF)^93–95^, code that measures and graphs the distance of different cell types to blood vessel cell types in two and three dimensions (see SOP^125^) has been developed^121,126^. Distance plots can be overlaid on the tissue section using Vitessce^66,67^ for 2D data and custom code^121,126^ for 3D data, see **Fig. 2.a, 8** and **Fig. 3.a** for examples.

#### Hierarchical community analysis of cell types

Hierarchical community analysis of cell types makes it possible to automatically detect multi-level functional tissue units^73^. The approach uses the single cell labels and x, y coordinates from spatial datasets. For the preview example featured in this paper, the dataset is a CODEX multiplexed imaging^86,119,127^ dataset of the healthy human intestine^73^. The original multiplexed imaging data was segmented and normalized and clustered using z-normalization of the antibody markers used and using Leiden unsupervised clustering^102^. Cell types were propagated to additional samples using the deep learning algorithm (STELLAR) for cell type label transfer in spatial single-cell data^118^.

Once cell type labels have been assigned, cell neighborhoods are calculated by clustering nearest neighbor (n=10) vectors surrounding each cell. A similar approach was taken to identify larger structures (termed communities^73^) using neighborhoods as the labels and taking a larger window for the nearest neighbors (n=100). Similarly, to identify major tissue units, community labels were used and an even larger window for nearest neighbors (n=300) prior to clustering of the vectors. Once all tissue structures have been identified, the connections in terms of primary components from various levels of tissue structures can be connected and visualized via a network graph. Currently, each node is organized per level and connected to the next spatial layer (e.g., cell type to neighborhood, neighborhood to community). This code is deposited on GitHub^71^.

### Atlas validation

Each digital object in the Human Reference Atlas is validated either by human expert review or using algorithmic means. HRA DO data formats depending on the type: ASCT+B is in CSV format, 3D Reference Organs in GL transmission binary (GLB) format, 2D FTUs in scalable vector graphics (SVG), etc. When this data is normalized to LinkML format, the source data is processed and structural errors in the raw data are identified. Once in its normalized form, LinkML is used to validate the structure of the transformed data, including ensuring data types and URLs fall within acceptable parameters. This step catches basic errors, including malformed URLs, missing data, and incorrect data types that can be a problem downstream. Beyond this, certain digital object types go through more advanced semantic checks to be sure that ontology terms used actually exist and that assertions from the digital object also appear in trusted ontologies like Uberon and Cell Ontology. Validation of the ASCT+B tables is most rigorous and includes detailed validation and reporting from the EBI team. While these tables are being authored, new/updated terms and links are related to the latest ontology versions available in Ubergraph (Uberon 2024-01-18 and CL 2024-01-05 for the 6th HRA) and weekly reports are generated at https://hubmapconsortium.github.io/ccf-validation-tools/ to aid table authors in getting the highest quality data for this critical piece of the HRA.

### Flexible hybrid cloud microservices architecture

#### Hybrid cloud

The IEC developed a hybrid cloud infrastructure that leverages the unique strengths of both on-premises and public cloud resources—each co-locating robust and scalable storage with robust and scalable computing— providing the flexibility to proactively adapt to evolving technologies and respond to the needs of the HuBMAP consortium and the broader atlasing community. As a key piece of this strategy, the HIVE IEC ingested, processed, and archived HuBMAP data at PSC. This approach provides flexible access, since the primary copy of HuBMAP data can be stored on-premises at a low cost, but then made available on any public or local resource without incurring substantial industry standard data egress charges, as well as free, low-friction access, since researchers can run basic analyses without having to create a public cloud account, or larger analyses by accessing the full HuBMAP data repository co-located with PSC’s national supercomputing infrastructure made available without charge to the research community.

#### Microservices architecture

The HuBMAP microservices architecture (**Supplemental Figure 1**) is built via agile development practices based on user-centered design, with microservices that communicate using REST APIs^128^ via Docker orchestration on AWS and on-premises resources. Each microservice is focused to serve specific functionality. Services are packaged into individual Docker containers. This orchestration of Docker containers is routinely built and rebuilt in development, test, and production instances which allows for independent operation and monitoring. This microservices architecture supports the plug-and-play of a continuously evolving set of algorithms required for experimental data ingestion, annotation, segmentation, search, filter, and visualization, as well as for atlas construction and usage. **Supplemental Figure 1** shows the resource, API, and application layers with exemplary modules, see **Supplemental Information** website for interactive version that lets users click on any module to access details. The core service that others are dependent on is the Entity API, backed by a Neo4j graph database, which provides the storage (creation, retrieval, update, and deletion) of all provenance and metadata information associated with HuBMAP data. The Search API allows for search of all provenance and metadata via the AWS hosted OpenSearch search engine, which holds a copy of all information maintained by the Entity API. The HuBMAP authentication and authorization model makes use of the Globus Auth service (https://globus.org) with login services in compliance with the OAuth2 standard (https://oauth.net/2), which provides user tokens that are passed among the services where they can be centrally validated and provide user authorization with linkage to defined groups via the Globus Groups service. The remaining services provide application specific functionality for support of data ingest and management (Ingest API), and unique entity tracking and identifier (ID) generation (UUID API).

#### HRA Cloud Infrastructure

Human Reference Atlas applications, including the HRA Portal, HRA Knowledge Graph, EUI, and RUI are all deployed to the web and hosted via Amazon Web Services (AWS) or GitHub Pages. For applications requiring server-side logic, Docker containers are created, tested, and built automatically with continuous integration / continuous deployment (CI/CD) via GitHub Actions, published to Amazon Elastic Container Registry (ECR), and then deployed via AWS AppRunner or Amazon Elastic Container Service (ECS). For applications which are served primarily as static files, they are tested and built automatically with CI/CD via GitHub Actions and then copied to Amazon S3 for serving or pushed to a branch for GitHub Pages deployment. Except for GitHub Pages, both static and server driven applications have Amazon CloudFront act as the front-end, providing a service mesh that allows for serving web requests, tracking usage, proxying requests to services running in AWS AppRunner or ECS, and caching frequently used files and responses. While Amazon Web Services are used extensively in the HRA Cloud Infrastructure, the technology is well suited to be adapted to other platforms.

## Data & Code Availability

All HuBMAP data is available via the HuBMAP Data Portal (https://portal.hubmapconsortium.org). Azimuth references can be accessed at https://azimuth.hubmapconsortium.org. HRA data and code are available at the HRA Portal (https://humanatlas.io).

Code is available on three different GitHub organizations: (1) https://github.com/hubmapconsortium is the HuBMAP organization where HRA started, (2) https://github.com/cns-iu is the organization owned by the Cyberinfrastructure for Network Science Center at Indiana University, and initial experimental HRA code starts here, (3) https://github.com/x-atlas-consortia was created recently to host cross-consortia code, including hra-kg, hra-pop, hra-apps, and hra-api.

HuBMAP and HRA primary and secondary data repositories are listed in **Supplemental Table 2** and HRA code repositories in **Supplemental Table 3**.

Supporting information is at https://cns-iu.github.io/hra-construction-usage-supporting-information.

## Full list of HRA Team authors

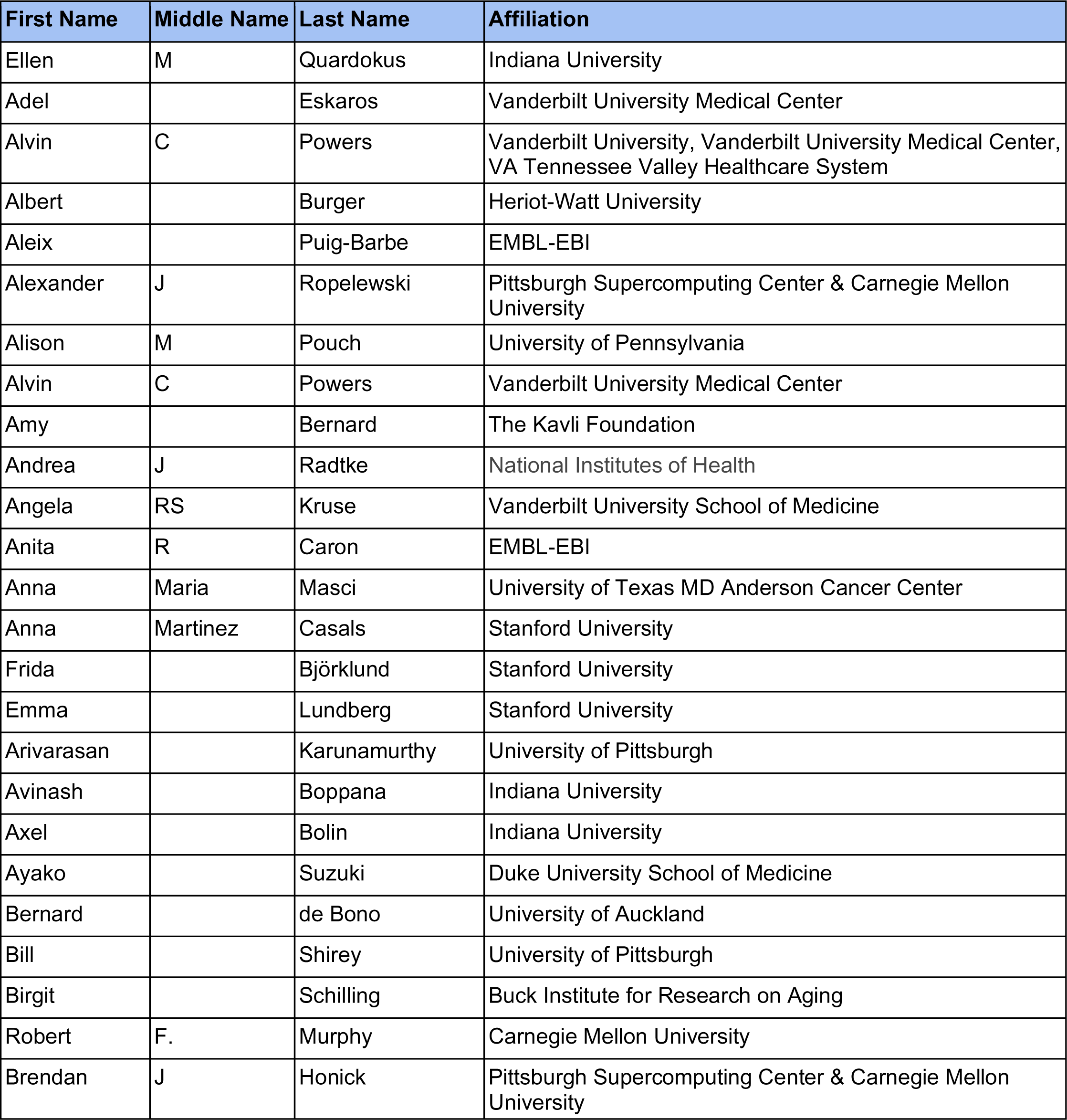
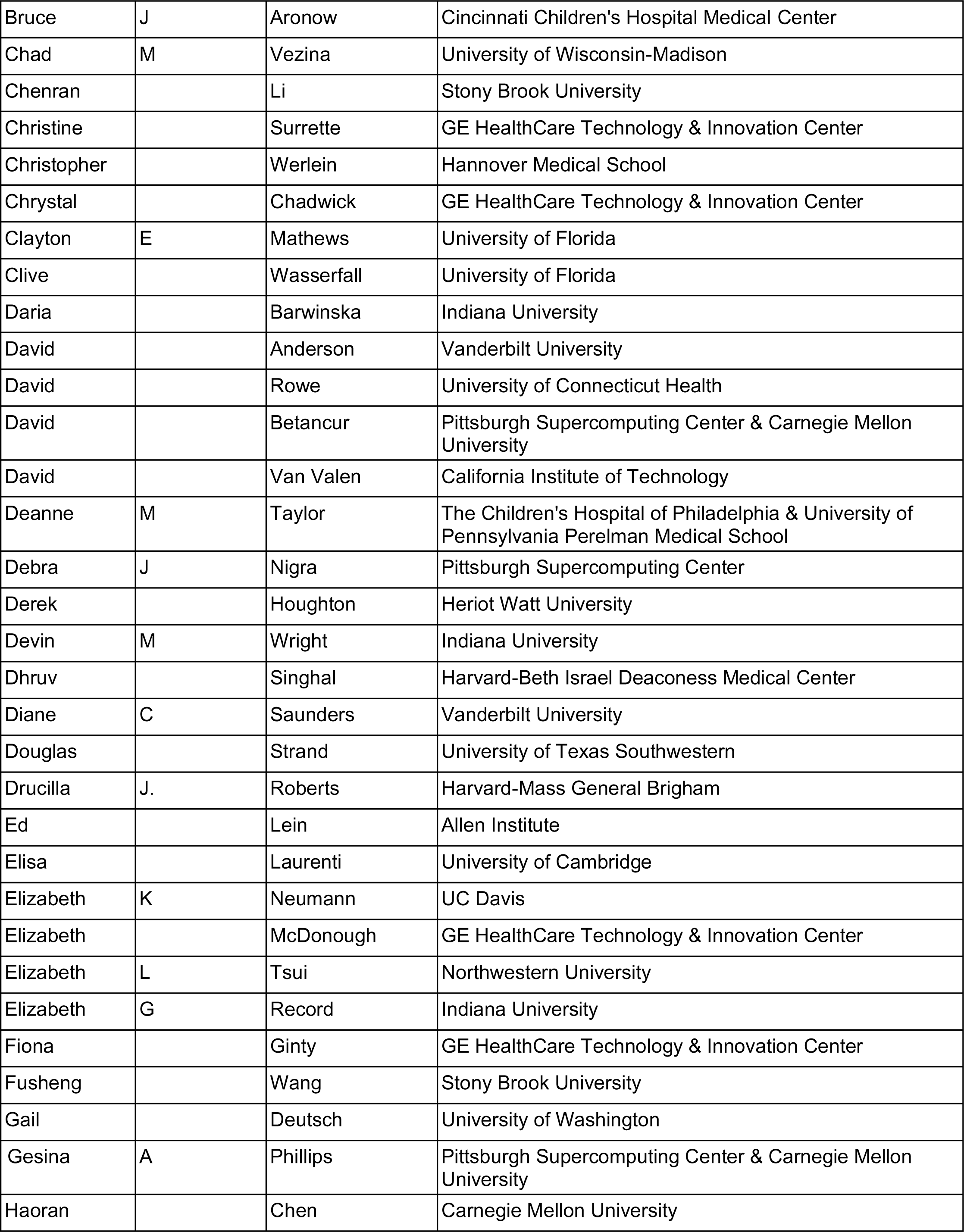
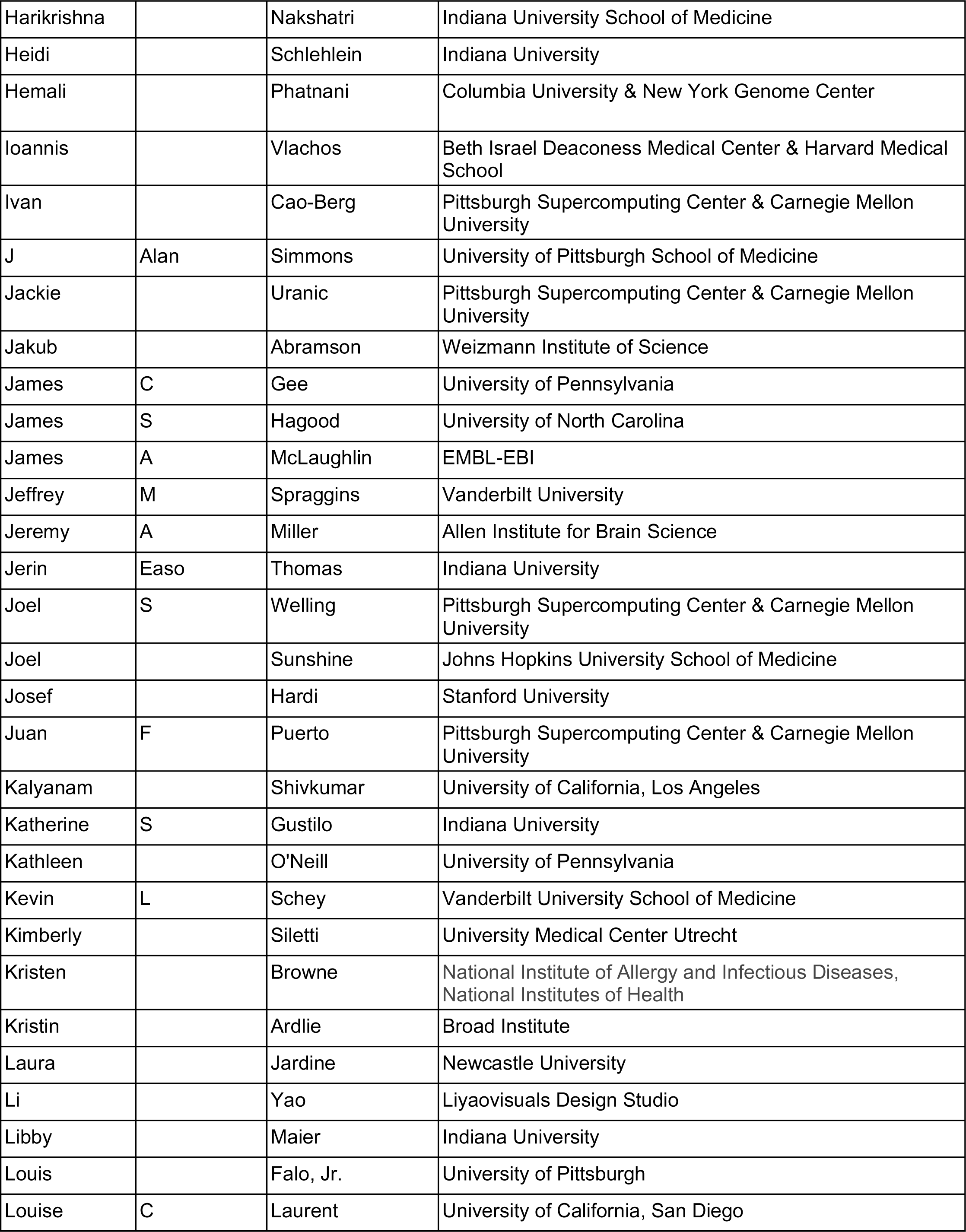
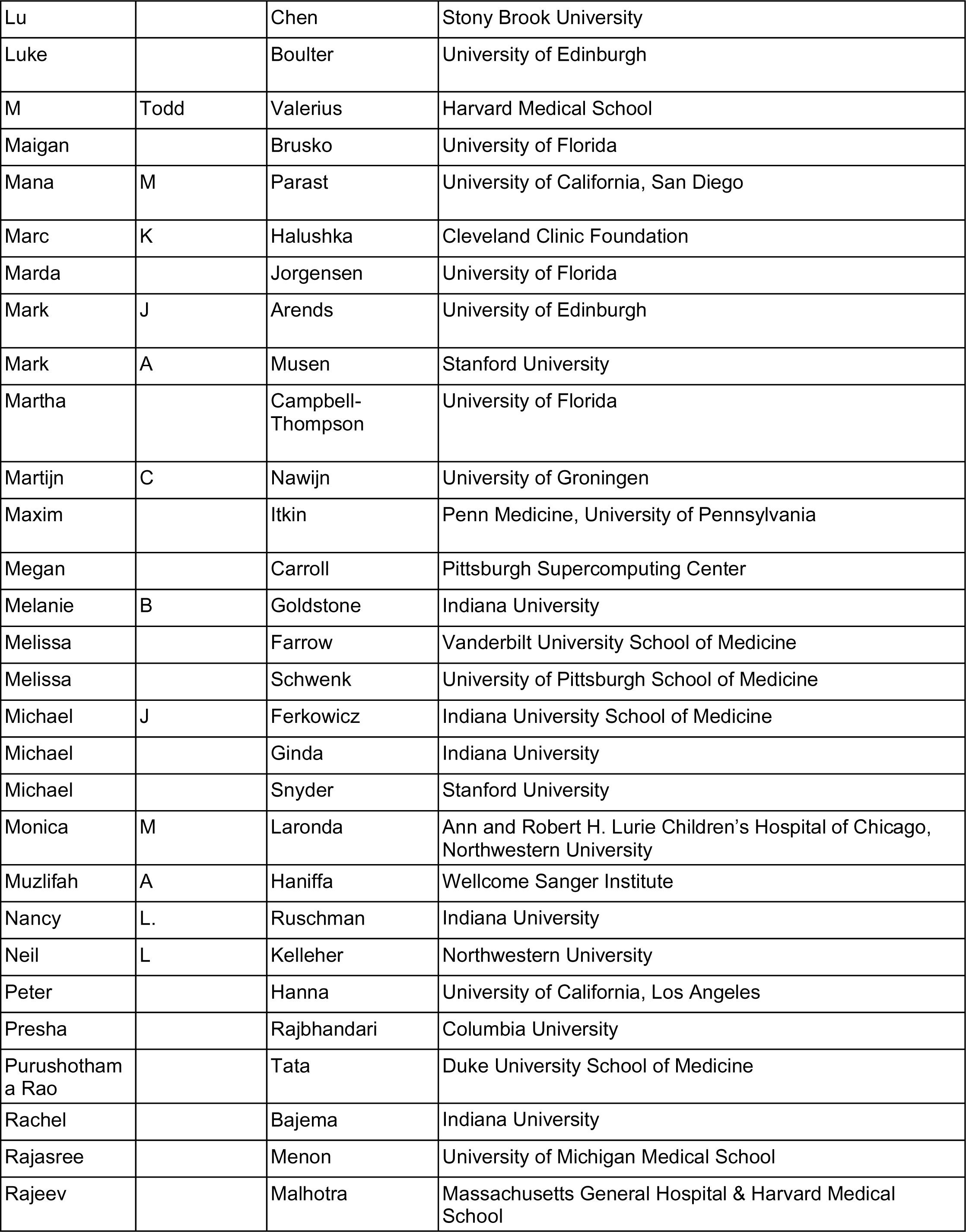
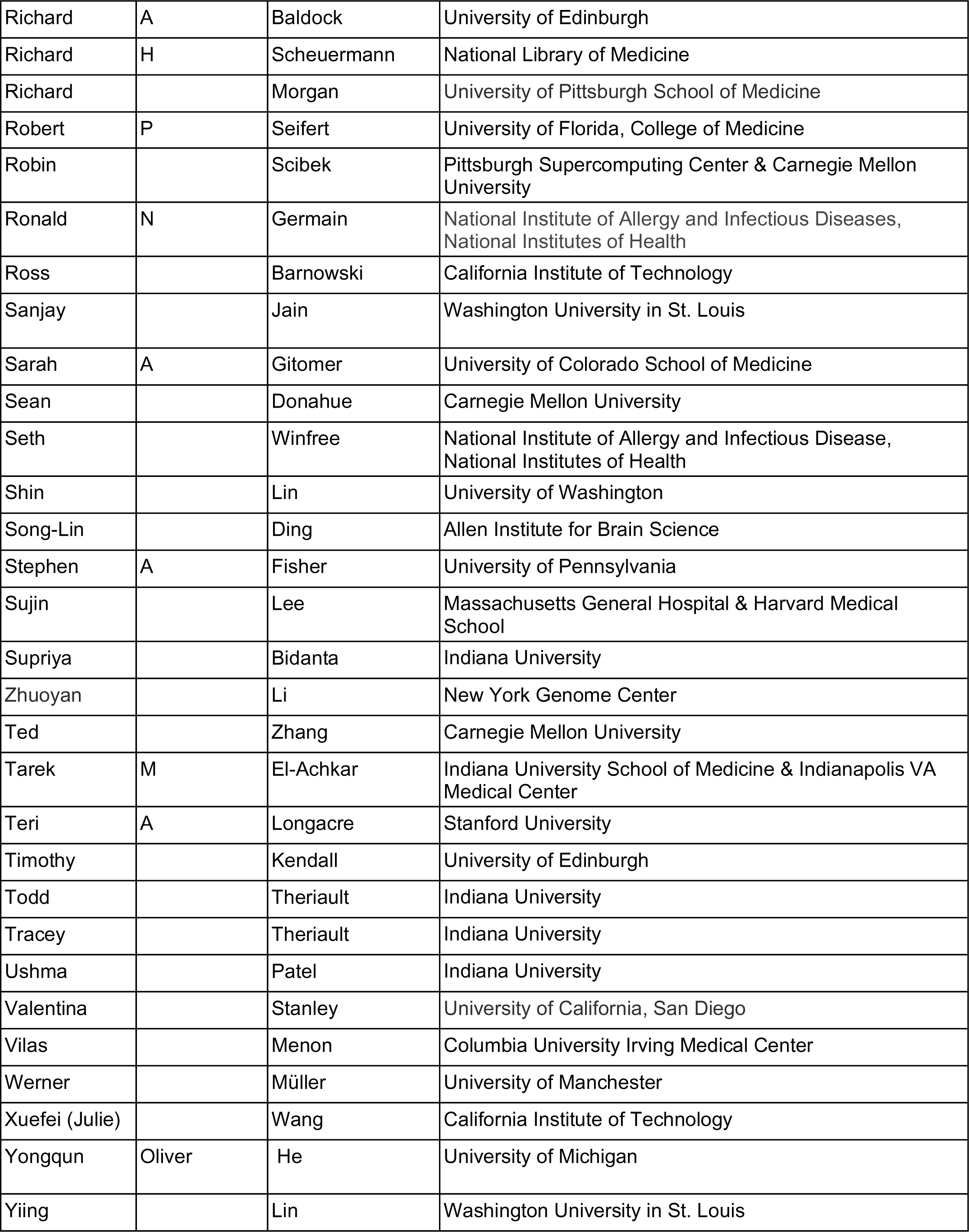
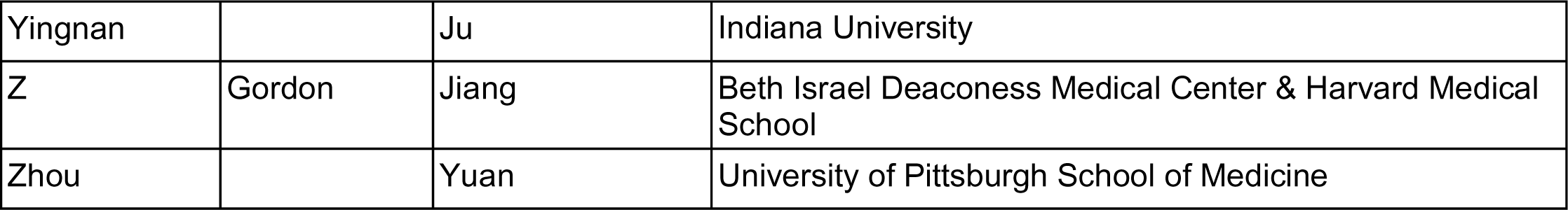

| First Name | Middle Name | Last Name | Affiliation |
| --- | --- | --- | --- |
| Ellen | M | Quardokus | Indiana University |
| Adel |  | Eskaros | Vanderbilt University Medical Center |
| Alvin | C | Powers | Vanderbilt University, Vanderbilt University Medical Center, VA Tennessee Valley Healthcare System |
| Albert |  | Burger | Heriot-Watt University |
| Aleix |  | Puig-Barbe | EMBL-EBI |
| Alexander | J | Ropelewski | Pittsburgh Supercomputing Center & Carnegie Mellon University |
| Alison | M | Pouch | University of Pennsylvania |
| Alvin | C | Powers | Vanderbilt University Medical Center |
| Amy |  | Bernard | The Kavli Foundation |
| Andrea | J | Radtke | National Institutes of Health |
| Angela | RS | Kruse | Vanderbilt University School of Medicine |
| Anita | R | Caron | EMBL-EBI |
| Anna | Maria | Masci | University of Texas MD Anderson Cancer Center |
| Anna | Martinez | Casals | Stanford University |
| Frida |  | Björklund | Stanford University |
| Emma |  | Lundberg | Stanford University |
| Arivarasan |  | Karunamurthy | University of Pittsburgh |
| Avinash |  | Boppana | Indiana University |
| Axel |  | Bolin | Indiana University |
| Ayako |  | Suzuki | Duke University School of Medicine |
| Bernard |  | de Bono | University of Auckland |
| Bill |  | Shirey | University of Pittsburgh |
| Birgit |  | Schilling | Buck Institute for Research on Aging |
| Robert | F. | Murphy | Carnegie Mellon University |
| Brendan | J | Honick | Pittsburgh Supercomputing Center & Carnegie Mellon University |
| Bruce | J | Aronow | Cincinnati Children's Hospital Medical Center |
| Chad | M | Vezina | University of Wisconsin-Madison |
| Chenran |  | Li | Stony Brook University |
| Christine |  | Surette | GE HealthCare Technology & Innovation Center |
| Christopher |  | Werlein | Hannover Medical School |
| Chrystal |  | Chadwick | GE HealthCare Technology & Innovation Center |
| Clayton | E | Mathews | University of Florida |
| Clive |  | Wasserfall | University of Florida |
| Daria |  | Barwinska | Indiana University |
| David |  | Anderson | Vanderbilt University |
| David |  | Rowe | University of Connecticut Health |
| David |  | Betancur | Pittsburgh Supercomputing Center & Carnegie Mellon University |
| David |  | Van Valen | California Institute of Technology |
| Deanne | M | Taylor | The Children's Hospital of Philadelphia & University of Pennsylvania Perelman Medical School |
| Debra | J | Nigra | Pittsburgh Supercomputing Center |
| Derek |  | Houghton | Heriot Watt University |
| Devin | M | Wright | Indiana University |
| Dhruv |  | Singhal | Harvard-Beth Israel Deaconess Medical Center |
| Diane | C | Saunders | Vanderbilt University |
| Douglas |  | Strand | University of Texas Southwestern |
| Drucilla | J. | Roberts | Harvard-Mass General Brigham |
| Ed |  | Lein | Allen Institute |
| Elisa |  | Laurenti | University of Cambridge |
| Elizabeth | K | Neumann | UC Davis |
| Elizabeth |  | McDonough | GE HealthCare Technology & Innovation Center |
| Elizabeth | L | Tsui | Northwestern University |
| Elizabeth | G | Record | Indiana University |
| Fiona |  | Ginty | GE HealthCare Technology & Innovation Center |
| Fusheng |  | Wang | Stony Brook University |
| Gail |  | Deutsch | University of Washington |
| Gesina | A | Phillips | Pittsburgh Supercomputing Center & Carnegie Mellon University |
| Haoran |  | Chen | Carnegie Mellon University |
| Harikrishna |  | Nakshatri | Indiana University School of Medicine |
| Heidi |  | Schlehlein | Indiana University |
| Hemali |  | Phatnani | Columbia University & New York Genome Center |
| Ioannis |  | Vlachos | Beth Israel Deaconess Medical Center & Harvard Medical School |
| Ivan |  | Cao-Berg | Pittsburgh Supercomputing Center & Carnegie Mellon University |
| J | Alan | Simmons | University of Pittsburgh School of Medicine |
| Jackie |  | Uranic | Pittsburgh Supercomputing Center & Carnegie Mellon University |
| Jakub |  | Abramson | Weizmann Institute of Science |
| James | C | Gee | University of Pennsylvania |
| James | S | Hagood | University of North Carolina |
| James | A | McLaughlin | EMBL-EBI |
| Jeffrey | M | Spraggins | Vanderbilt University |
| Jeremy | A | Miller | Allen Institute for Brain Science |
| Jerin | Easo | Thomas | Indiana University |
| Joel | S | Welling | Pittsburgh Supercomputing Center & Carnegie Mellon University |
| Joel |  | Sunshine | Johns Hopkins University School of Medicine |
| Josef |  | Hardi | Stanford University |
| Juan | F | Puerto | Pittsburgh Supercomputing Center & Carnegie Mellon University |
| Kalyanam |  | Shivkumar | University of California, Los Angeles |
| Katherine | S | Gustilo | Indiana University |
| Kathleen |  | O'Neill | University of Pennsylvania |
| Kevin | L | Schey | Vanderbilt University School of Medicine |
| Kimberly |  | Siletti | University Medical Center Utrecht |
| Kristen |  | Browne | National Institute of Allergy and Infectious Diseases, National Institutes of Health |
| Kristin |  | Ardlie | Broad Institute |
| Laura |  | Jardine | Newcastle University |
| Li |  | Yao | Liyaovisuals Design Studio |
| Libby |  | Maier | Indiana University |
| Louis |  | Falo, Jr. | University of Pittsburgh |
| Louise | C | Laurent | University of California, San Diego |
| Lu |  | Chen | Stony Brook University |
| Luke |  | Boulter | University of Edinburgh |
| M | Todd | Valerius | Harvard Medical School |
| Maigan |  | Brusko | University of Florida |
| Mana | M | Parast | University of California, San Diego |
| Marc | K | Halushka | Cleveland Clinic Foundation |
| Marda |  | Jorgensen | University of Florida |
| Mark | J | Arends | University of Edinburgh |
| Mark | A | Musen | Stanford University |
| Martha |  | Campbell-Thompson | University of Florida |
| Martijn | C | Nawijn | University of Groningen |
| Maxim |  | Itkin | Penn Medicine, University of Pennsylvania |
| Megan |  | Carroll | Pittsburgh Supercomputing Center |
| Melanie | B | Goldstone | Indiana University |
| Melissa |  | Farrow | Vanderbilt University School of Medicine |
| Melissa |  | Schwenk | University of Pittsburgh School of Medicine |
| Michael | J | Ferkowicz | Indiana University School of Medicine |
| Michael |  | Ginda | Indiana University |
| Michael |  | Snyder | Stanford University |
| Monica | M | Laronda | Ann and Robert H. Lurie Children's Hospital of Chicago, Northwestern University |
| Muzlifah | A | Haniffa | Wellcome Sanger Institute |
| Nancy | L. | Ruschman | Indiana University |
| Neil | L | Kelleher | Northwestern University |
| Peter |  | Hanna | University of California, Los Angeles |
| Presha |  | Rajbhandari | Columbia University |
| Purushotham a Rao |  | Tata | Duke University School of Medicine |
| Rachel |  | Bajema | Indiana University |
| Rajasree |  | Menon | University of Michigan Medical School |
| Rajeev |  | Malhotra | Massachusetts General Hospital & Harvard Medical School |
| Richard | A | Baldock | University of Edinburgh |
| Richard | H | Scheuermann | National Library of Medicine |
| Richard |  | Morgan | University of Pittsburgh School of Medicine |
| Robert | P | Seifert | University of Florida, College of Medicine |
| Robin |  | Scibek | Pittsburgh Supercomputing Center & Carnegie Mellon University |
| Ronald | N | Germain | National Institute of Allergy and Infectious Diseases, National Institutes of Health |
| Ross |  | Barnowski | California Institute of Technology |
| Sanjay |  | Jain | Washington University in St. Louis |
| Sarah | A | Gitomer | University of Colorado School of Medicine |
| Sean |  | Donahue | Carnegie Mellon University |
| Seth |  | Winfree | National Institute of Allergy and Infectious Disease, National Institutes of Health |
| Shin |  | Lin | University of Washington |
| Song-Lin |  | Ding | Allen Institute for Brain Science |
| Stephen | A | Fisher | University of Pennsylvania |
| Sujin |  | Lee | Massachusetts General Hospital & Harvard Medical School |
| Supriya |  | Bidanta | Indiana University |
| Zhuoyan |  | Li | New York Genome Center |
| Ted |  | Zhang | Carnegie Mellon University |
| Tarek | M | El-Achkar | Indiana University School of Medicine & Indianapolis VA Medical Center |
| Teri | A | Longacre | Stanford University |
| Timothy |  | Kendall | University of Edinburgh |
| Todd |  | Theriault | Indiana University |
| Tracey |  | Theriault | Indiana University |
| Ushma |  | Patel | Indiana University |
| Valentina |  | Stanley | University of California, San Diego |
| Vilas |  | Menon | Columbia University Irving Medical Center |
| Werner |  | Müller | University of Manchester |
| Xuefei (Julie) |  | Wang | California Institute of Technology |
| Yongqun | Oliver | He | University of Michigan |
| Yiing |  | Lin | Washington University in St. Louis |
| Yingnan |  | Ju | Indiana University |
| Z | Gordon | Jiang | Beth Israel Deaconess Medical Center & Harvard Medical School |
| Zhou |  | Yuan | University of Pittsburgh School of Medicine |

## Supporting information

ZIP file with 17 supplemental figures

## Acknowledgements

Thanks go to Adam Taylor, Ajay Pillai, Avi Ma’ayan, David Osumi-Sutherland, Gary Bader, Rafael Gonçalves, Rahul Satija, Sebastian Lobentanzer, and Zorina Galis for their expert comments and suggestions on an earlier version of this paper. We thank Can Ergen for sharing popV code and an initial crosswalk to CL, plus for validating popV results.

The Human Reference Atlas (HRA) is under active development by HuBMAP, the Cellular Senescence Network (SenNet) Consortium, the Kidney Precision Medicine Project (KPMP), the National Institute of Diabetes and Digestive and Kidney Diseases (NIDDK), and the GenitoUrinary Development Molecular Anatomy Project (GUDMAP) projects with expert input by the HRA Editorial Board and in close collaboration with experts from more than 18 other consortia. K.B. and S.A.T. are co-directors of and are funded by the CIFAR MacMillan Multiscale Human program.

This research has been supported by the NIH Common Fund through the Office of Strategic Coordination/Office of the NIH Director under awards:

● OT2OD033756 (K.B., Y.Z., G.M.W., Y.J., D.Q., A.B., B.W.H.) and OT2OD026671 (K.B., G.M.W., Y.J., D.Q., A.B., B.W.H.);
● U54 HL165443 and HLU01148861 (G.P., R.M., J.P.);
● 1R03OD036499 (Y.Z.);
● OT2OD026673 (J.F.) and 3U54AG075936 (J.W.H.);
● OT2OD026675 and OT2OD033759 (P.B., J.C.S., A.B.);
● 3OT2OD033760 (R.S., G.M.);
● as well as 3OT2OD026682 and 1OT2OD033761 (M.R., S.A.T., C.X.).

Further, this work was supported by:

● the SenNet CODCC under award number U24CA268108 (K.B, J.C.S., Y.J., D.Q., A.B., B.W.H.);
● by the NIDDK under award U24DK135157 (K.B., D.Q., B.W.H.);
● by the KPMP grant U2CDK114886 (K.B., Y.J., D.Q., A.B., B.W.H.);
● by the National Human Genome Research Institute RM1HG011014 (R.S.);
● and the NIH National Institute of Allergy and Infectious Diseases (NIAID), Department of Health and Human Services under BCBB Support Services Contract HHSN316201300006W/HHSN27200002.

This research was supported in part by the Intramural Research Program of the U.S. National Institutes of Health. The funders had no role in study design, data collection and analysis, decision to publish, or preparation of the manuscript.

## Supplemental Figures

**Supplemental Figure 1.**
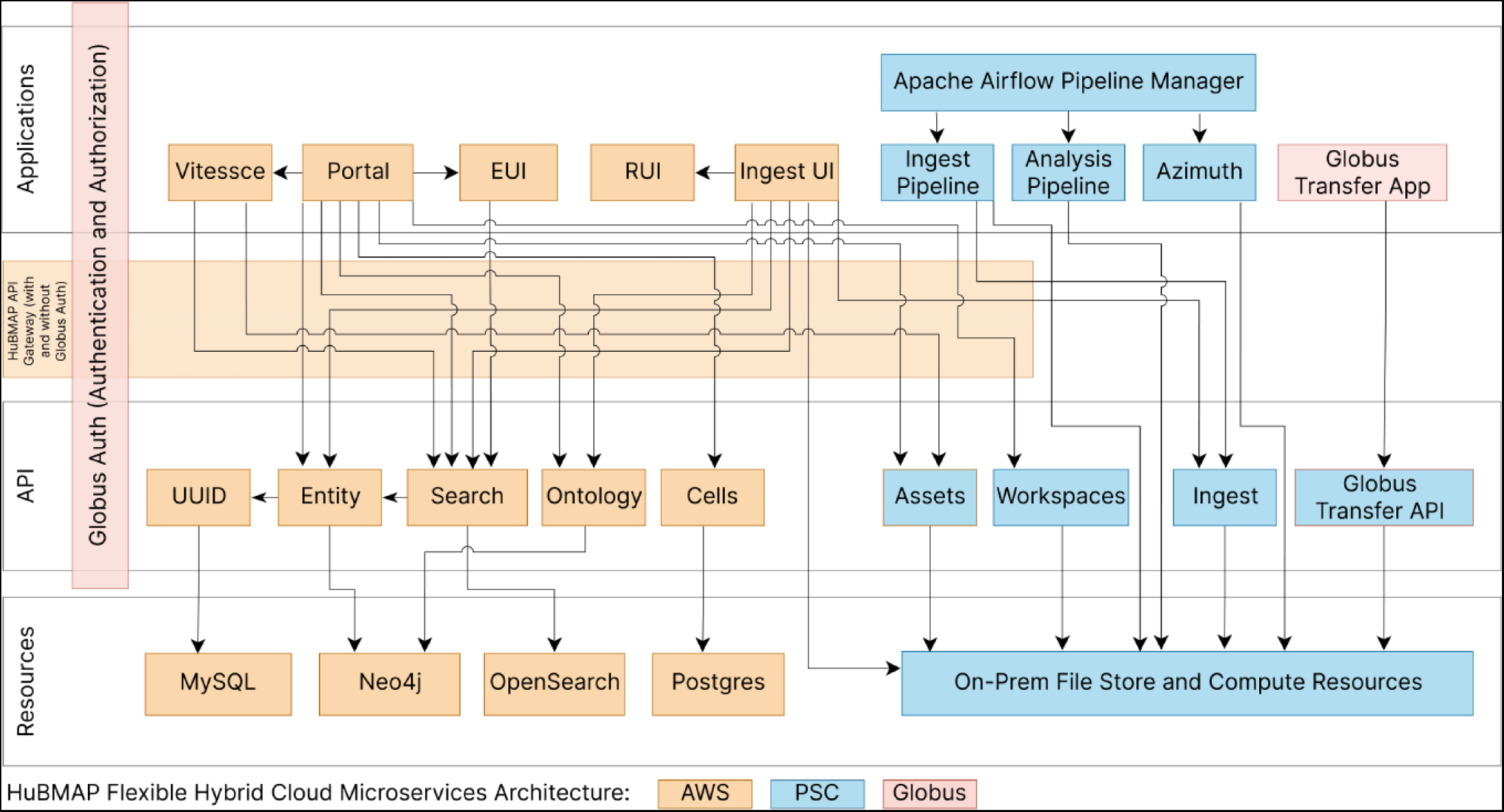
**Hybrid Cloud Microservices System Architecture**

**Supplemental Figure 2.**
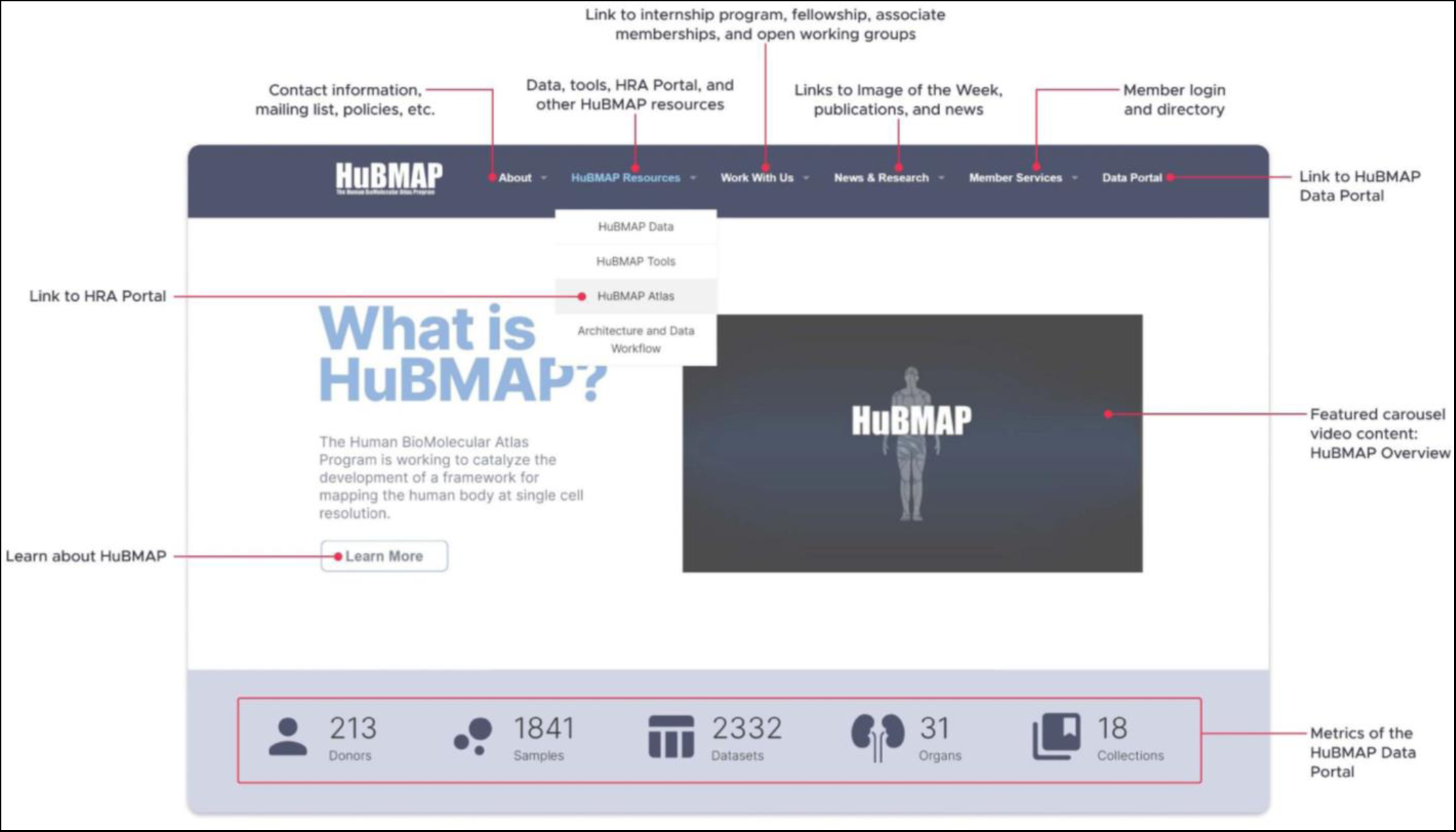
**HuBMAP Consortium Website**

**Supplemental Figure 3.**
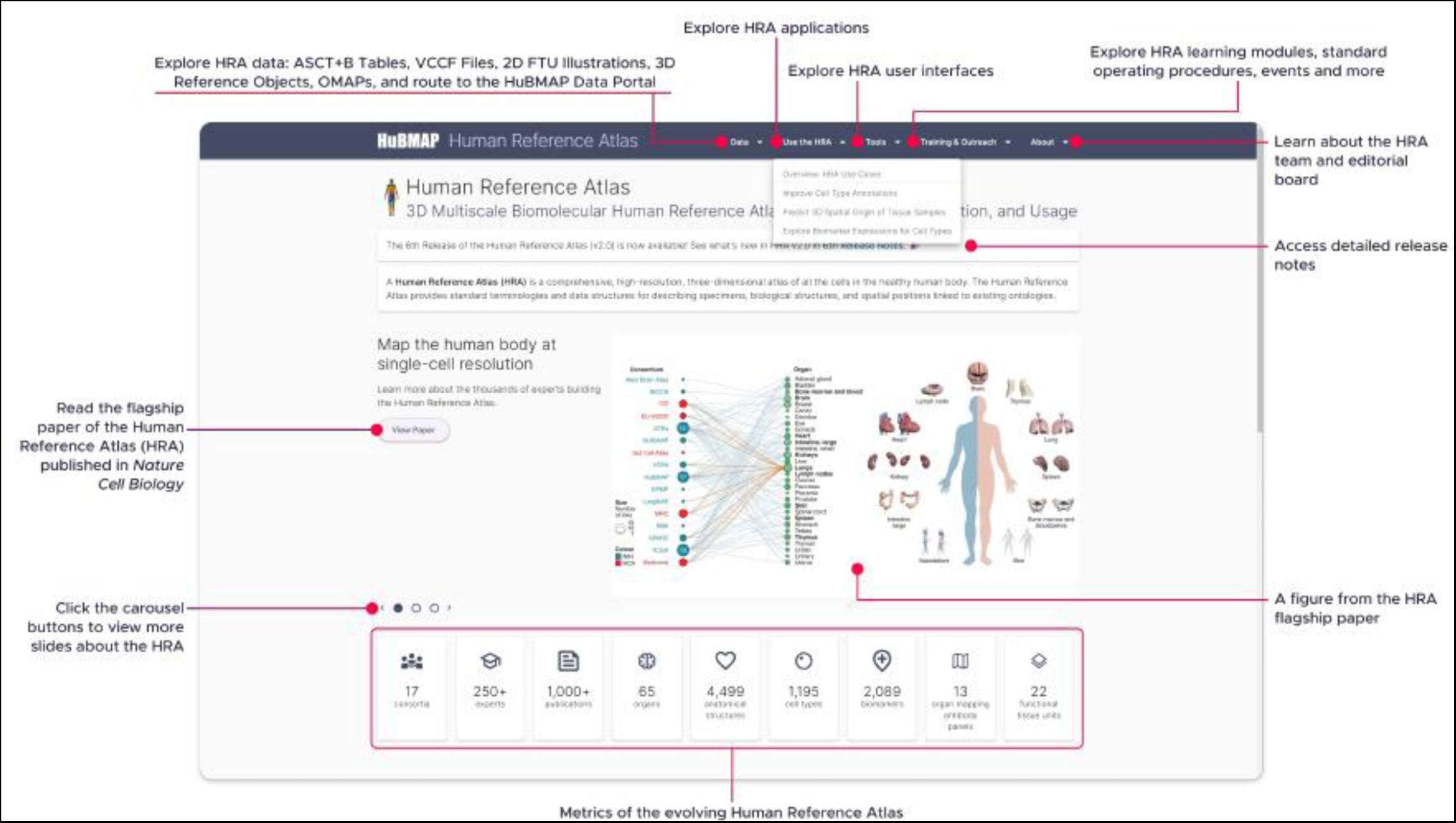
**Human Reference Atlas Portal**

**Supplemental Figure 4:**
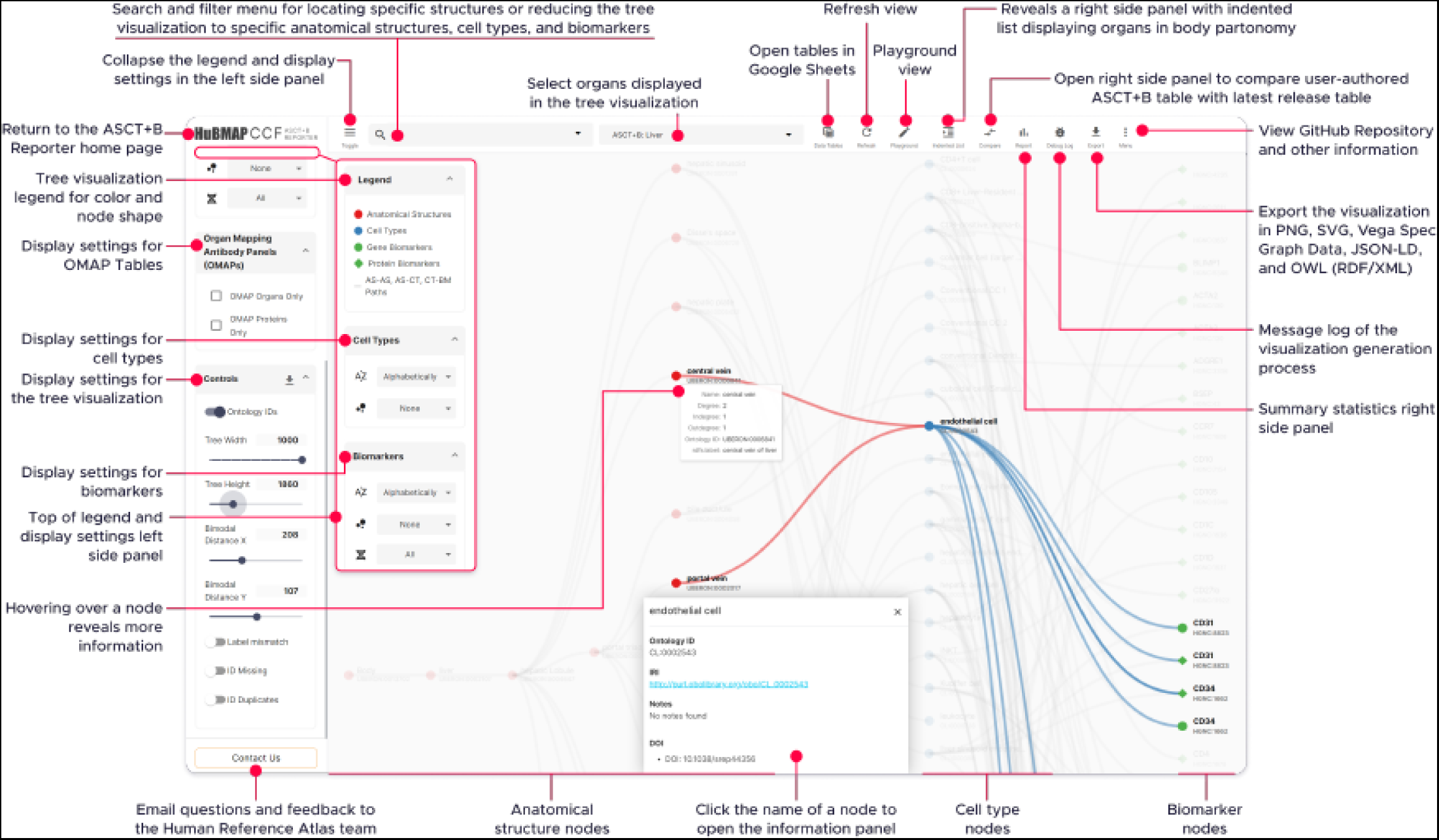
**ASCT+B Reporter User Interface**

**Supplemental Figure 5.**
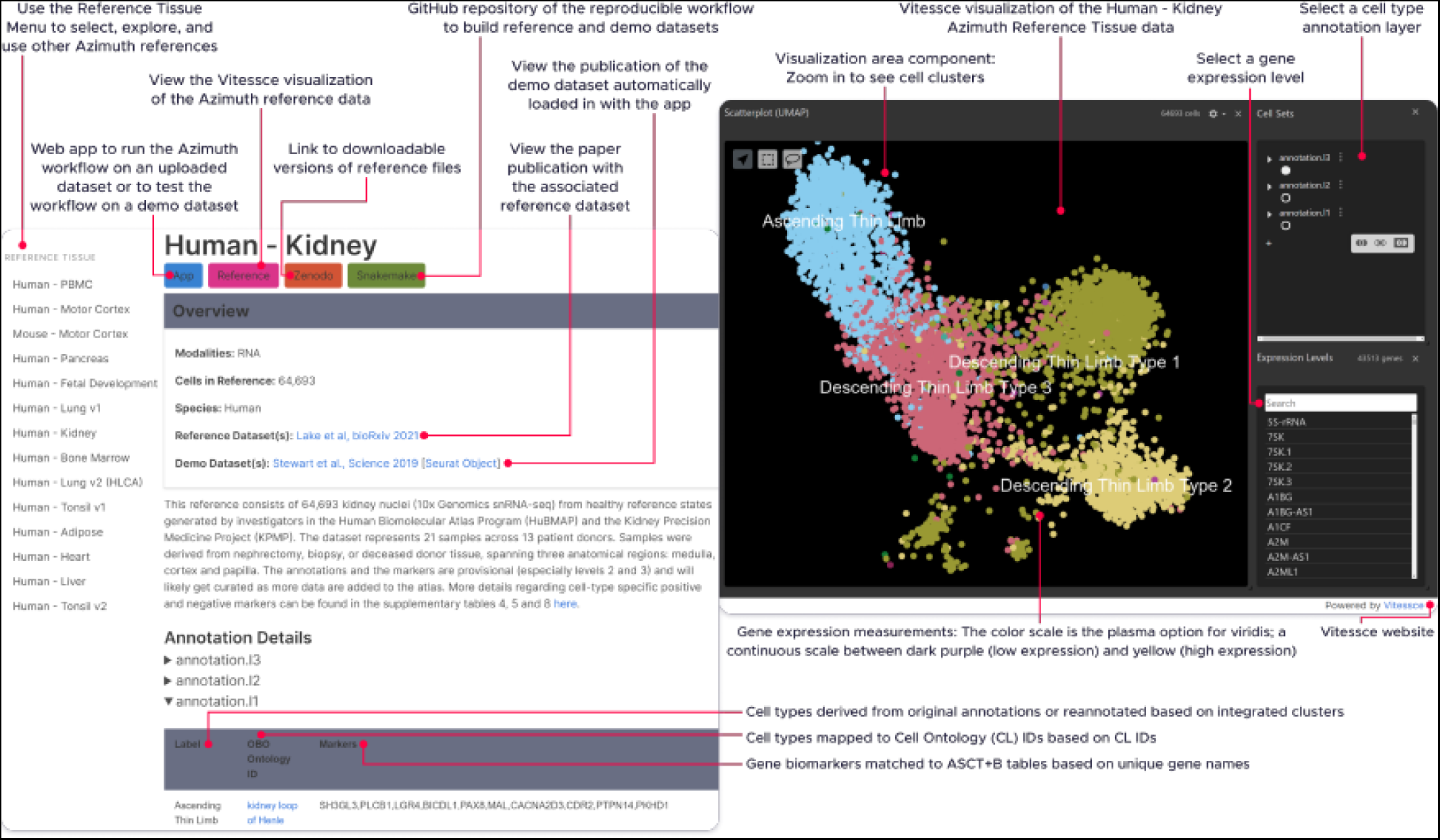
**Azimuth Portal and Reference Explorer User Interface**

**Supplemental Figure 6:**
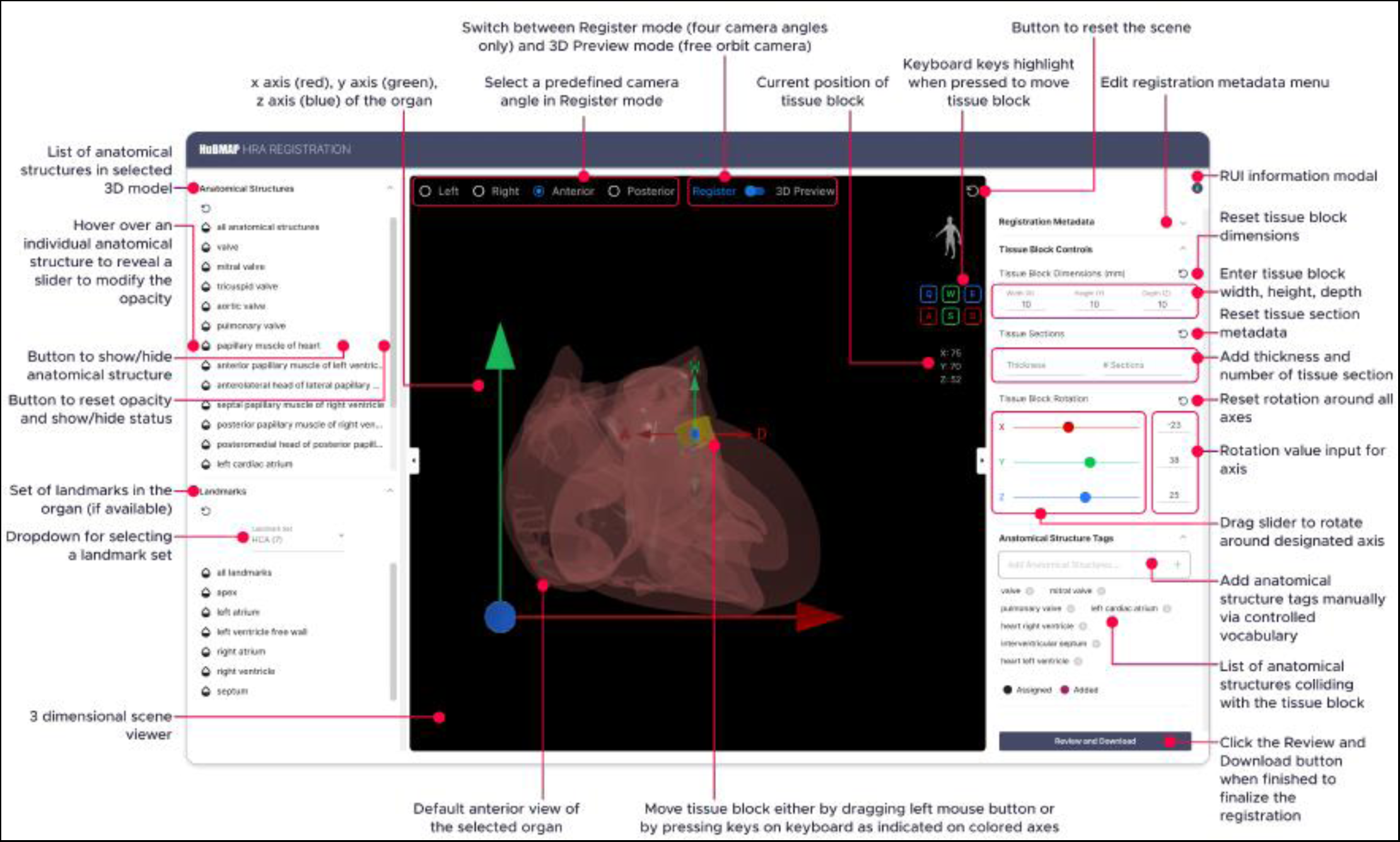
**Registration User Interface (RUI)**

**Supplemental Figure 7:**
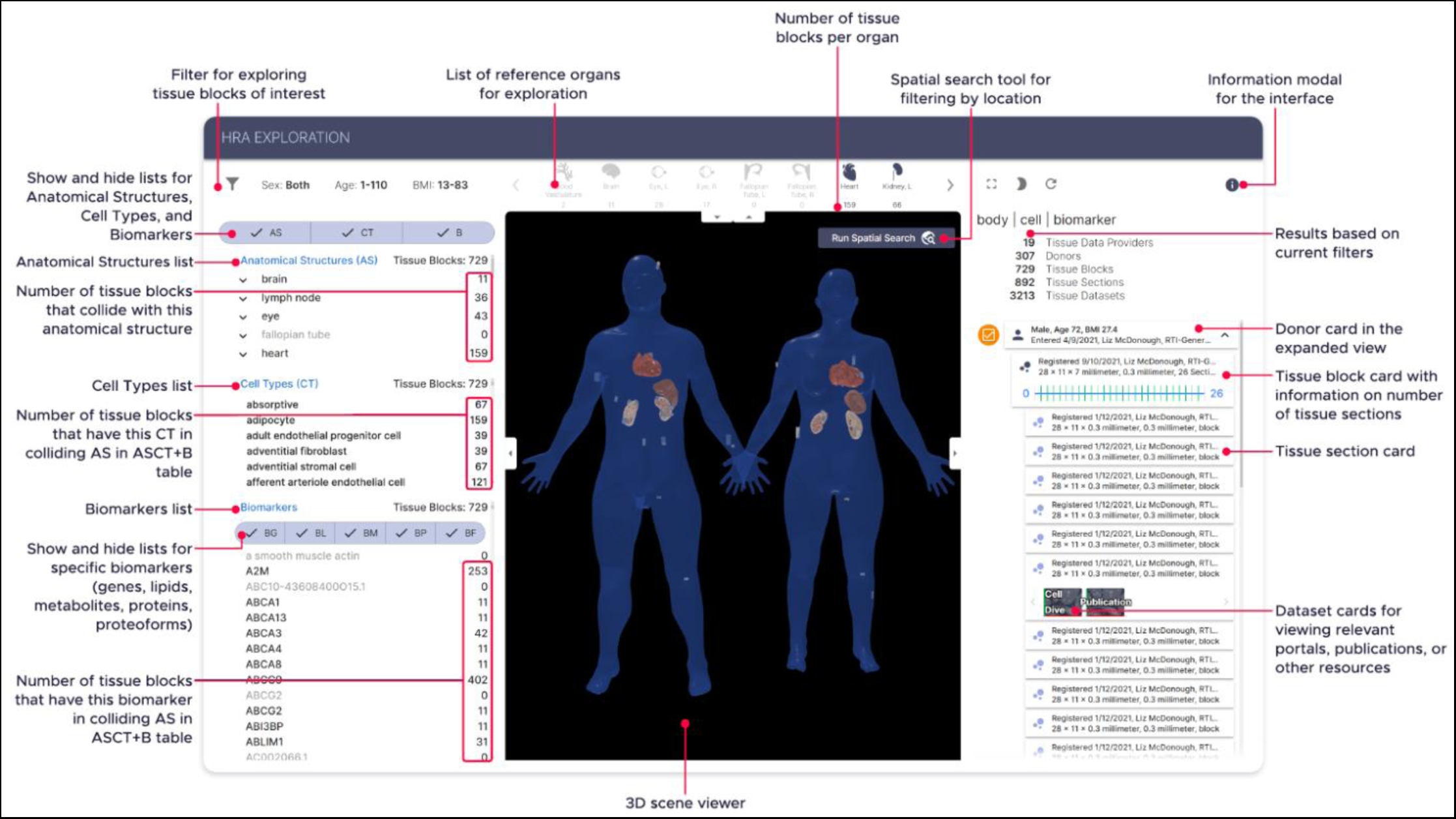
**Exploration User Interface (EUI)**

**Supplemental Figure 8:**
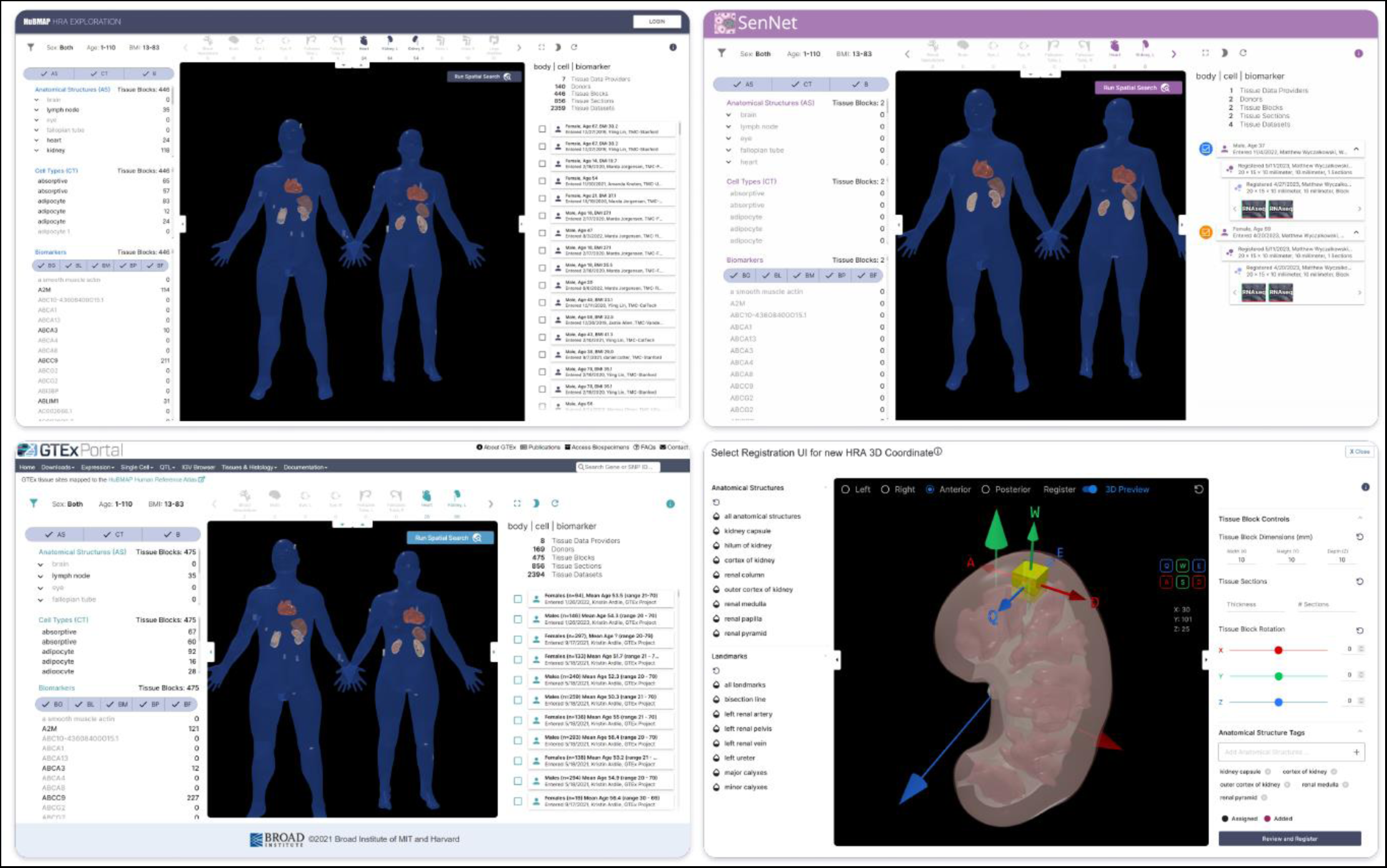
**Customized, Branded Deployment of EUI in HuBMAP, SenNet, GTEx, and RUI in GUDMAP**

**Supplemental Figure 9:**
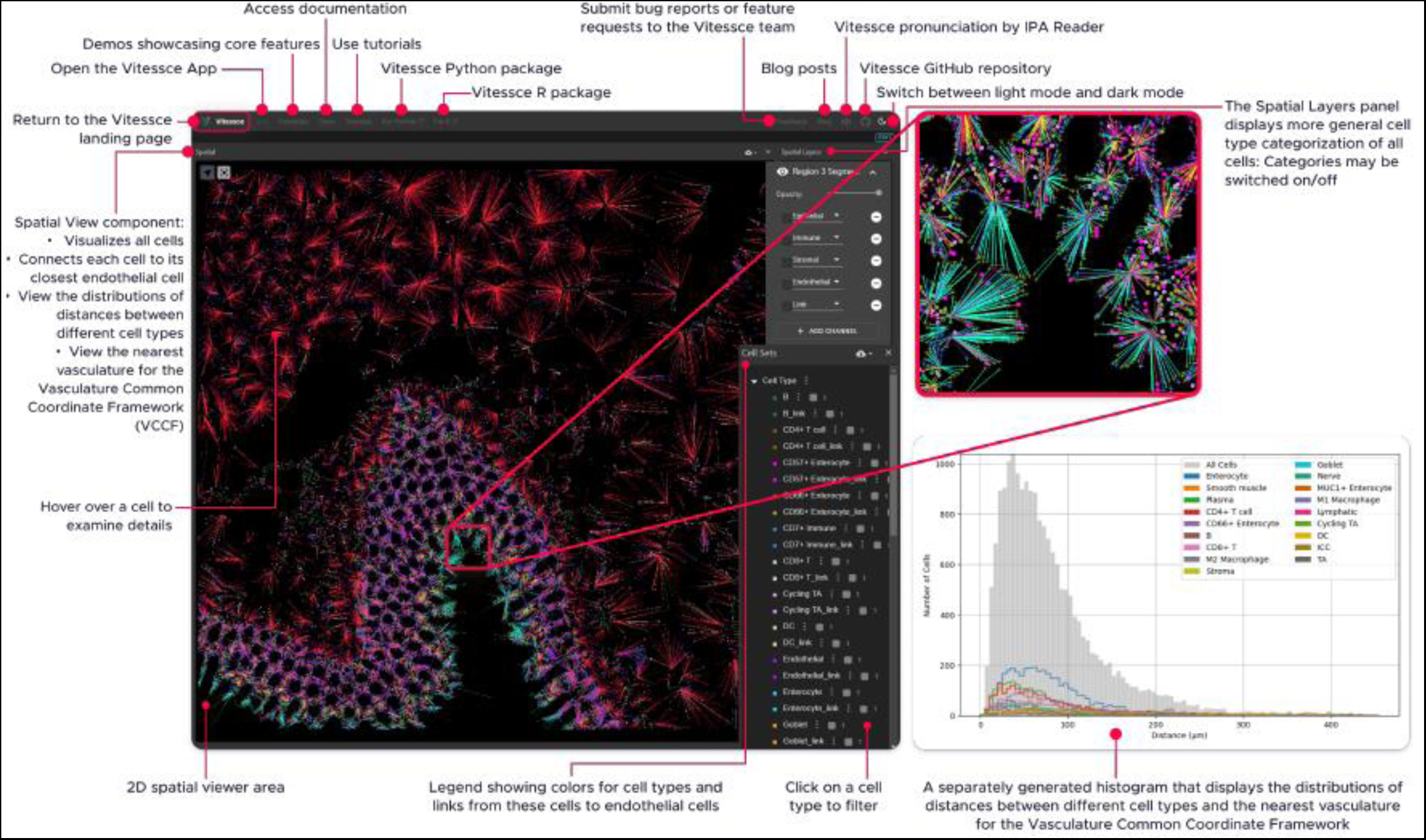
**Cell-Cell Distance Distribution Visualizations**

**Supplemental Figure 10:**
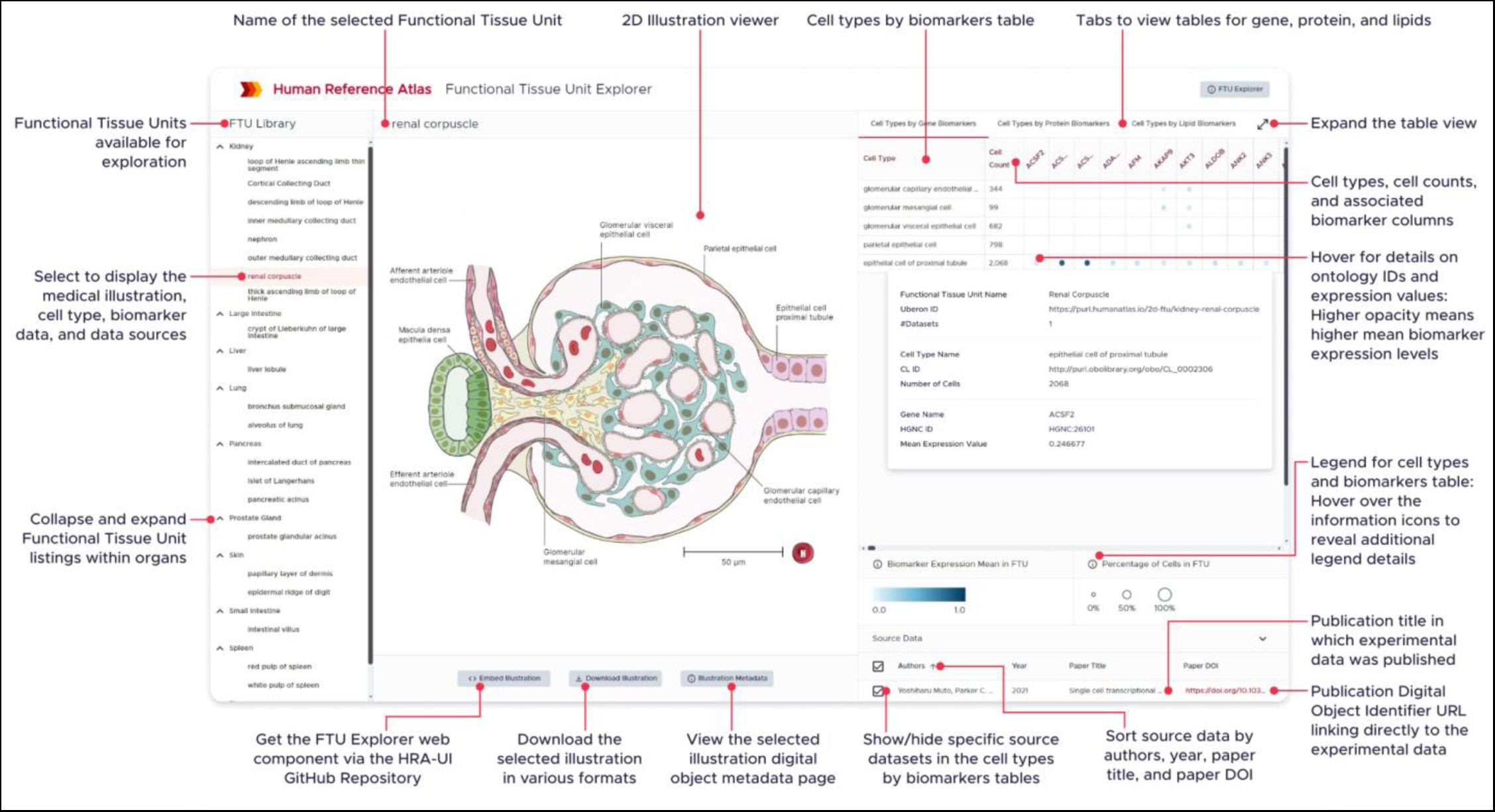
**Interactive FTU Explorer**

**Supplemental Figure 11:**
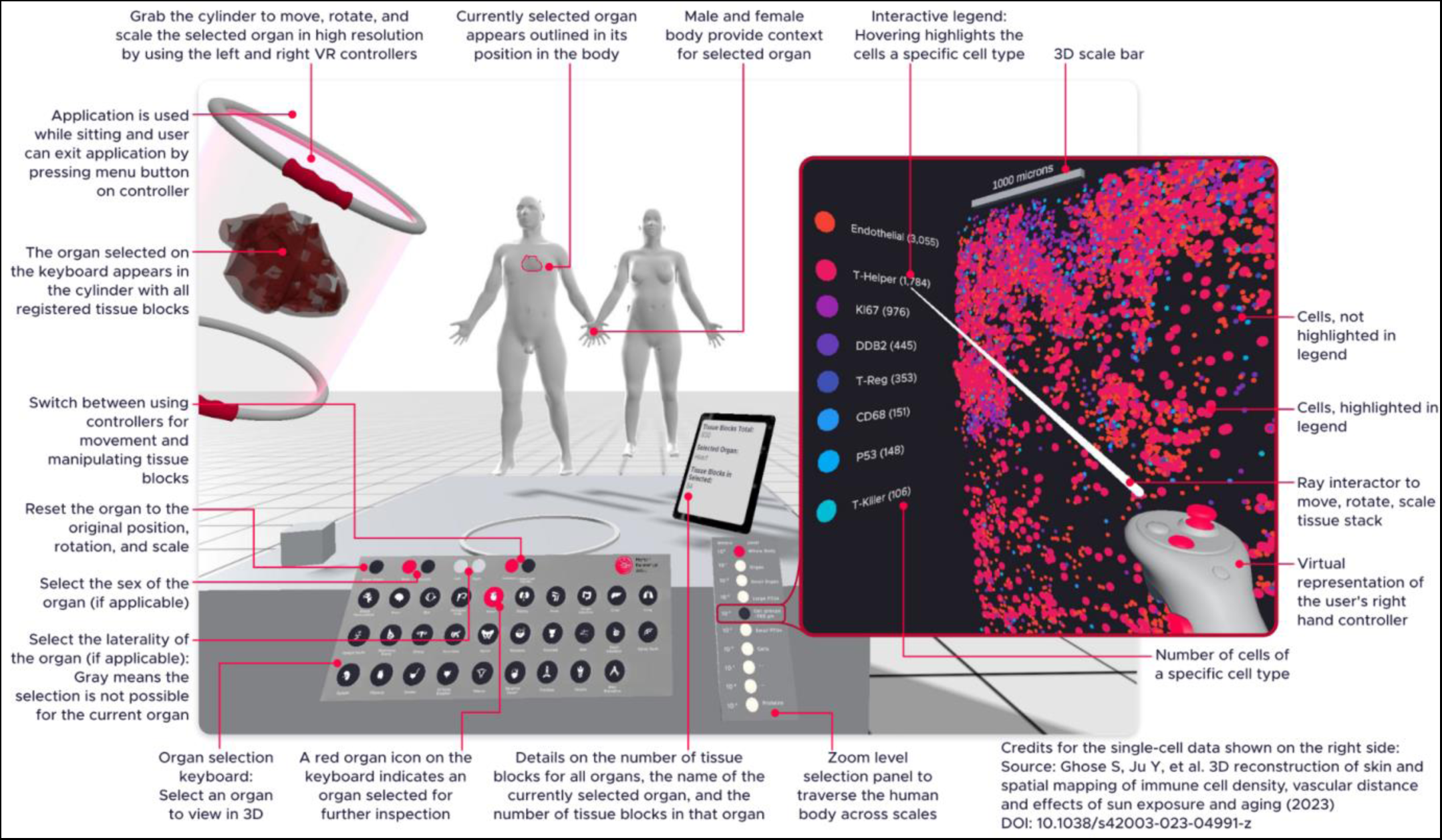
**HRA Organ Gallery in VR**

**Supplemental Figure 12:**
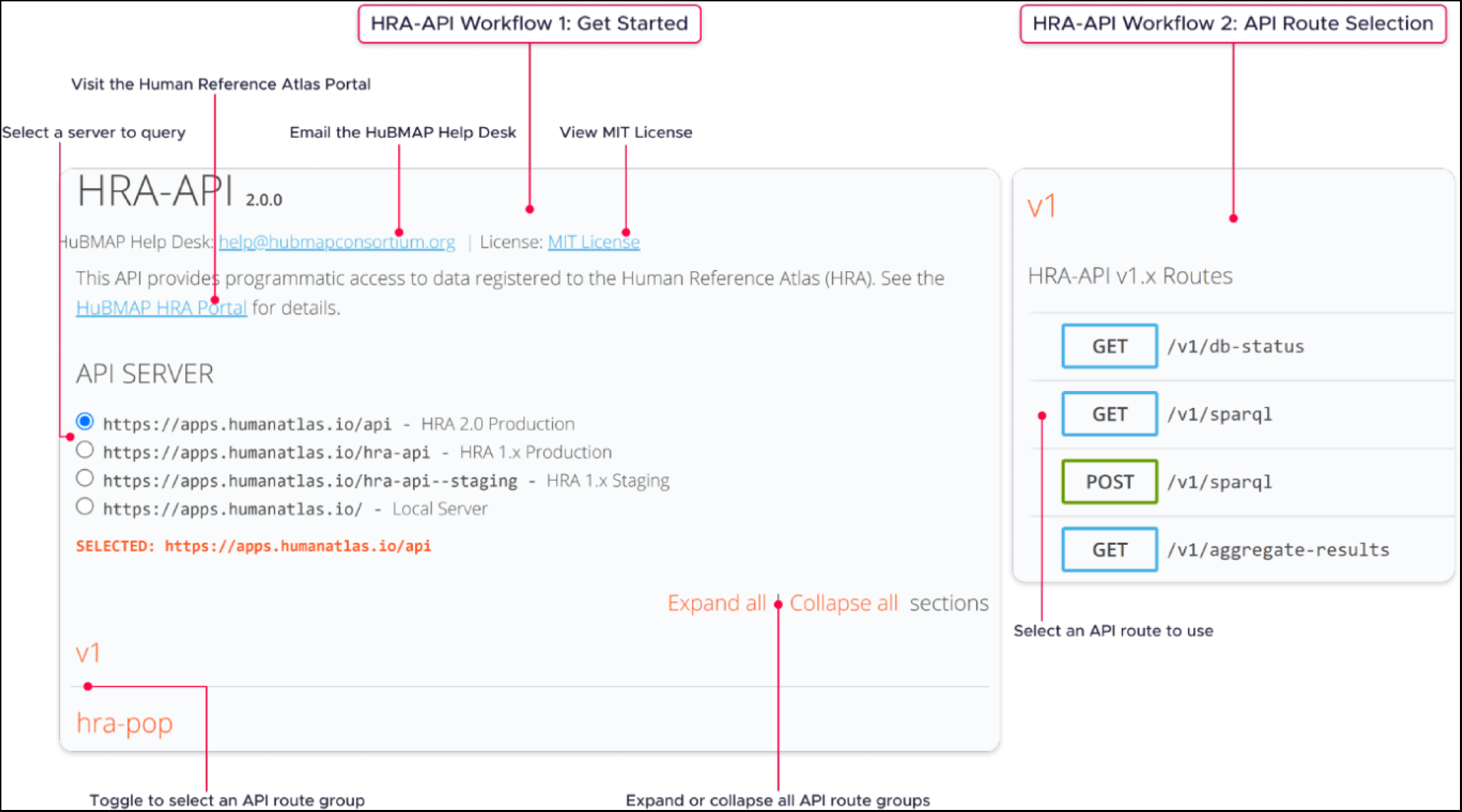
**Human Reference Atlas Application Programming Interface: Get Started and API Route Selection**

**Supplemental Figure 13:**
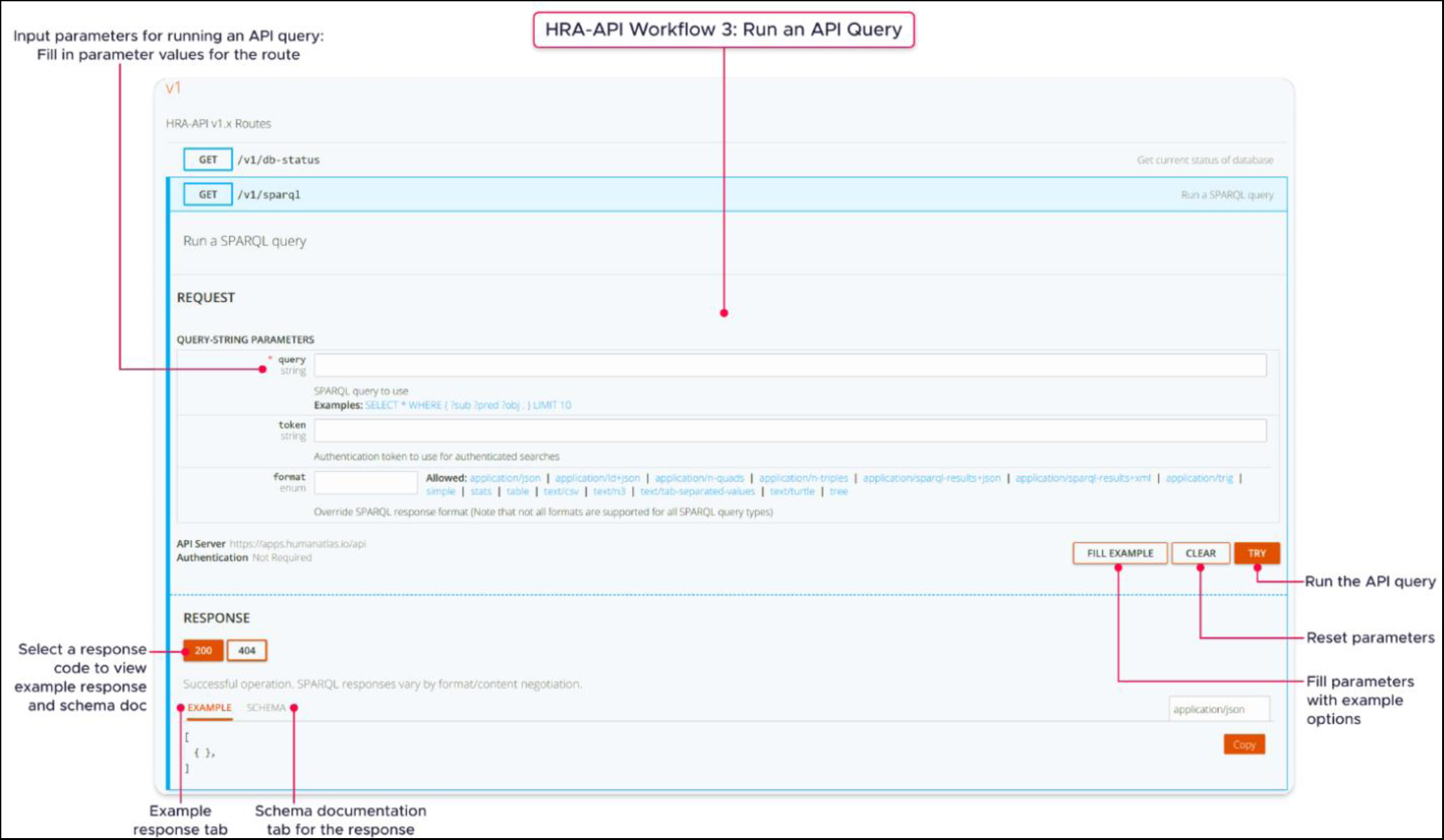
**Human Reference Atlas Application Programming Interface: Run an API Query**

**Supplemental Figure 14:**
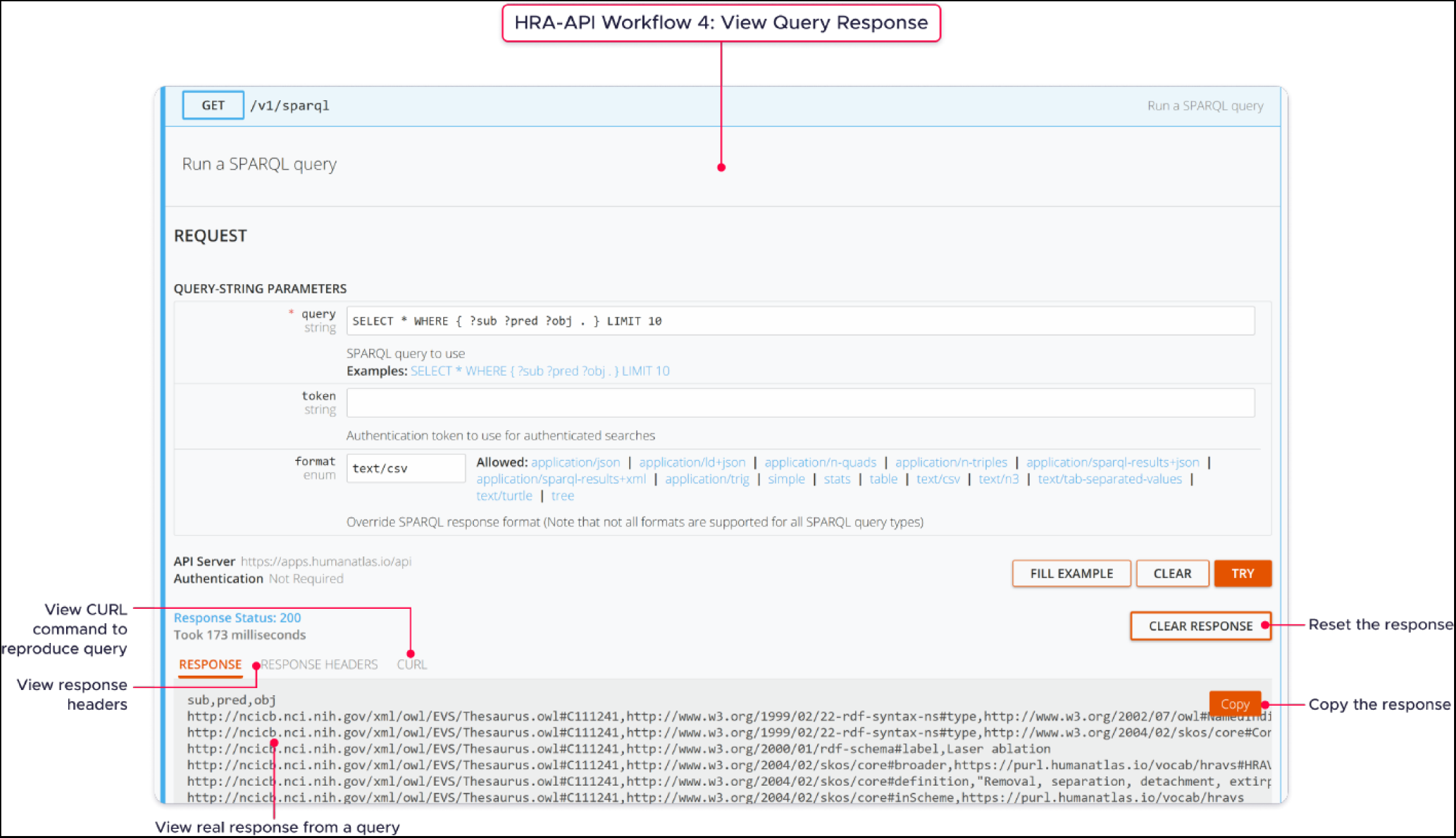
**Human Reference Atlas Application Programming Interface: View Query Response**

**Supplemental Figure 15.**
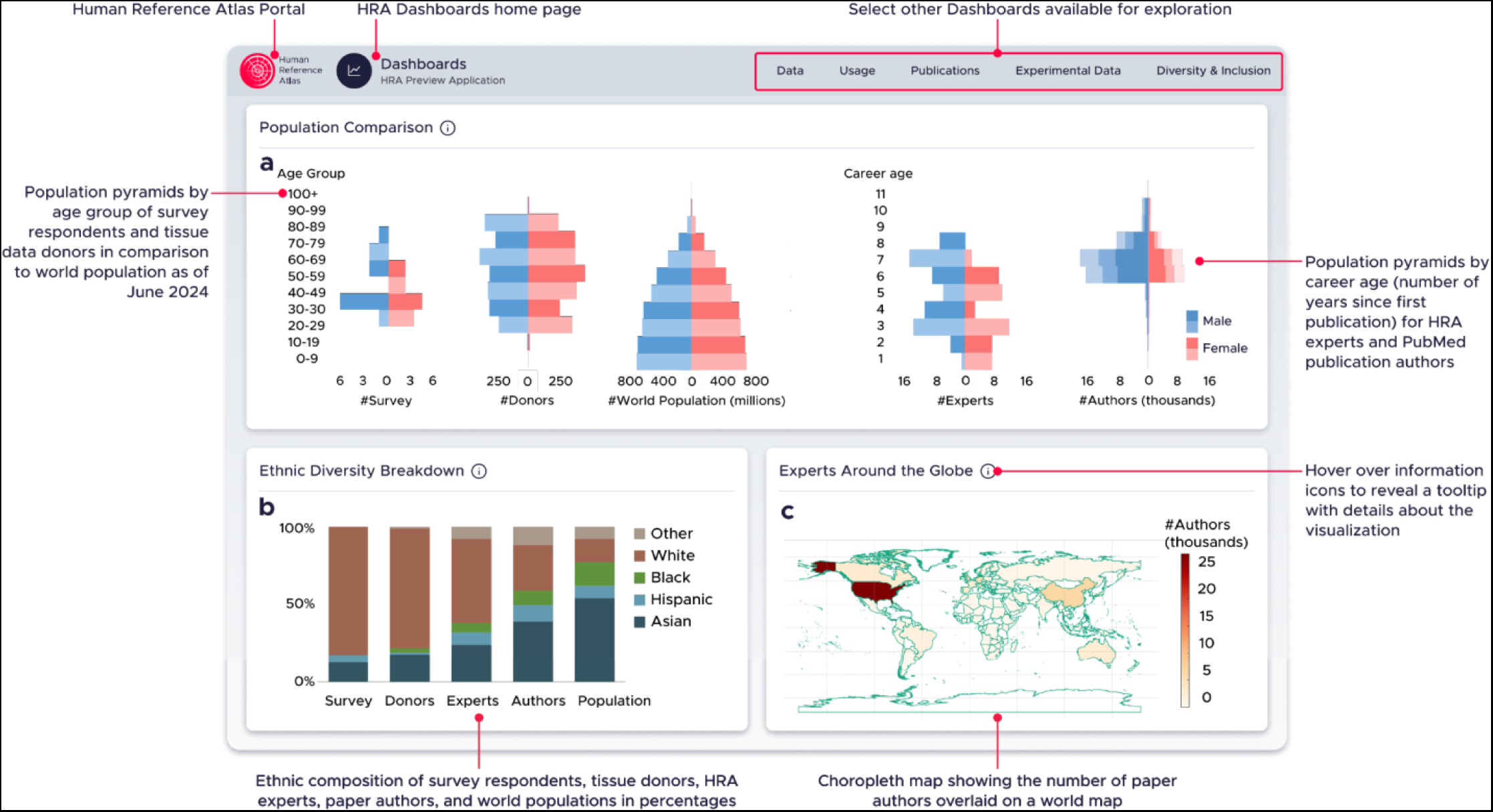
**HRA Equity Dashboard**

**Supplemental Figure 16:**
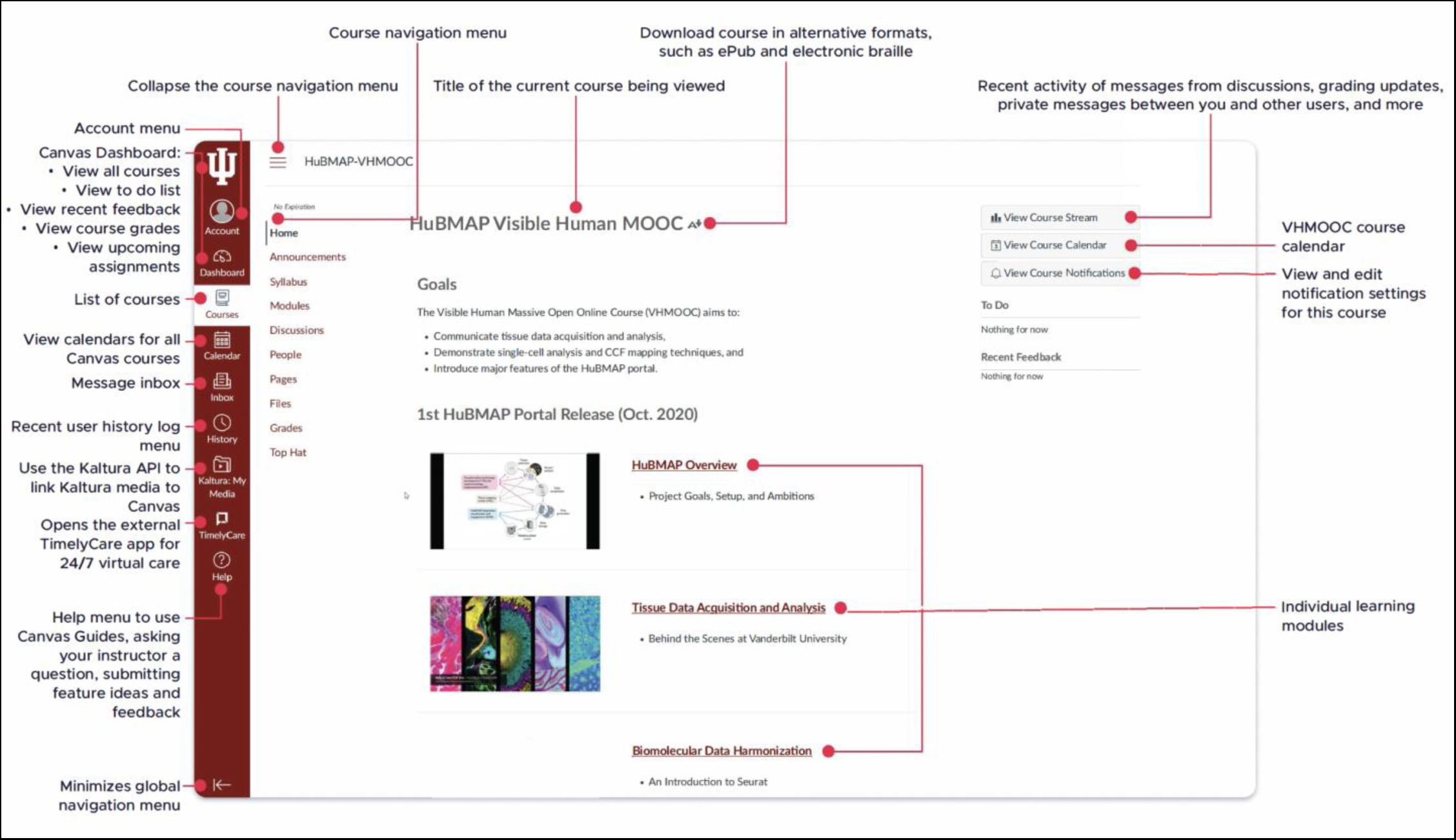
**Visible Human Massive Open Online Course (VHMOOC)**

**Supplemental Figure 17:**
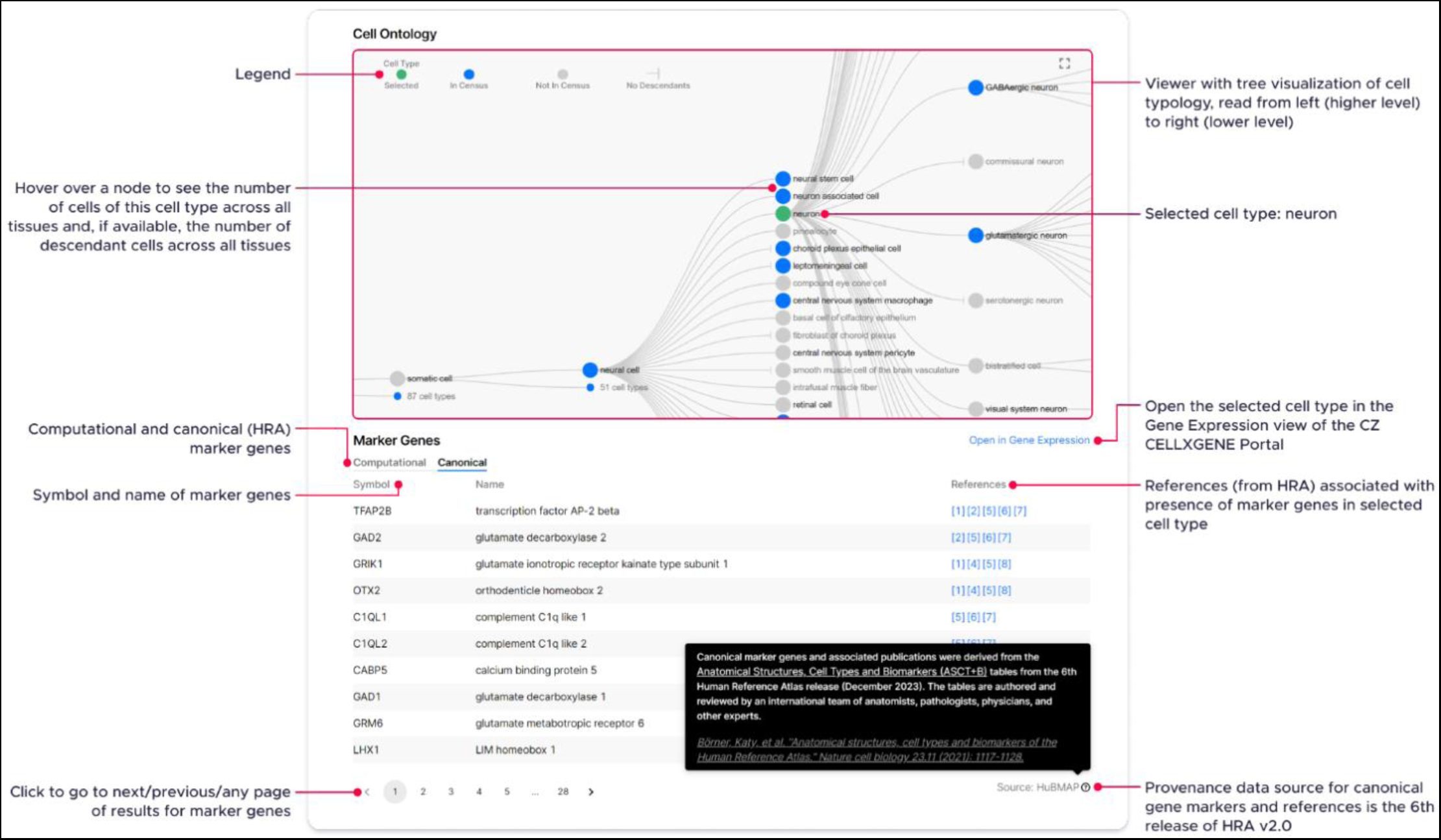
**CZ CellGuide Visualization of Cell Typology with ‘Canonical’ Marker Genes and ‘References’ from the HRA**

## Supplemental Tables

**Supplemental Table 1:** Experimental Data, HRA Data and Cell Type Annotation by Organ. Number of sc/snRNA-seq datasets per organ vs. number of 3D reference organs plus anatomical structure and unique cell types in ASCT+B tables vs. number of cell types that different cell type annotation tools can assign.

| Organ | Datasets with H5AD file | ASCT+B and 3D Reference Organs |  |  | Cell Type Annotation Tools |  |  |
| --- | --- | --- | --- | --- | --- | --- | --- |
|  |  | #AS in 3D (male+female) | #AS | #CT | Azimuth | CellTypist | popV |
| blood | 2,501 | none | 1 | 29 | 41 | 26 | 18 |
| blood vasculature | 3 | 244 | 1,054 | 10 | 0 | 0 | 14 |
| bone marrow | 131 | none | 1 | 47 | 43 | 39 | 14 |
| brain* | 656 | 210 | 188 | 625 | 0 | 0 | 0 |
| brain > motor cortex* | Part of 656 | 4** | 1 | 139 | 20 | 0 | 0 |
| brain > hippocampus* | Part of 656 | 12** | 4 | 45 | 0 | 20 | 0 |
| breast/mammary gland | 1 | 18 | 3 | 10 | 0 | 0 | 14 |
| eye | 192 | 96 | 39 | 55 | 0 | 0 | 27 |
| heart | 254 | 34 | 52 | 28 | 25 | 49 | 6 |
| kidney | 207 | 116 | 61 | 70 | 58 | 34 | 0 |
| large intestine | 140 | 22 | 54 | 58 | 0 | 0 | 16 |
| liver | 70 | 46 | 17 | 30 | 25 | 35 | 12 |
| lung | 531 | 141 | 54 | 74 | 78 | 71 | 36 |
| lymph node*** | 16 | 16 | 34 | 45 | 0 | 28 | 22 |
| pancreas | 14 | 11 | 31 | 30 | 11 | 9 | 14 |
| prostate gland | 34 | 18 | 13 | 19 | 0 | 0 | 13 |
| skin of body | 53 | 2 | 15 | 35 | 0 | 23 | 21 |
| small intestine | 177 | 23 | 39 | 34 | 0 | 0 | 16 |
| spleen | 39 | 12 | 37 | 59 | 0 | 32 | 22 |
| thymus | 47 | 6 | 18 | 50 | 0 | 0 | 21 |
| trachea | 29 | 8 | 20 | 17 | 0 | 0 | 18 |
| urinary bladder | 0 | 15 | 16 | 15 | 0 | 0 | 14 |
| uterus | 23 | 10 | 61 | 18 | 0 | 0 | 13 |
| **Total (sum, not unique)** | **5,118** | **1,064** | **1,810** | **1,542** | **301** | **366** | **331** |

\* Azimuth and CellTypist focus on the motor cortex and hippocampus, respectively. The 3D Reference Object and ASCT+B Table of the brain in the HRA focus on the entire organ.
\*\* Primary motor cortex is part of the precentral gyrus and hippocampus has three (head, body, tail) 3D reference objects in each hemisphere in male and female.
\*\*\* The 3D lymph node is generic and was placed in the mesenteric region as all HuBMAP data is from that region.

**Supplemental Table 2:**
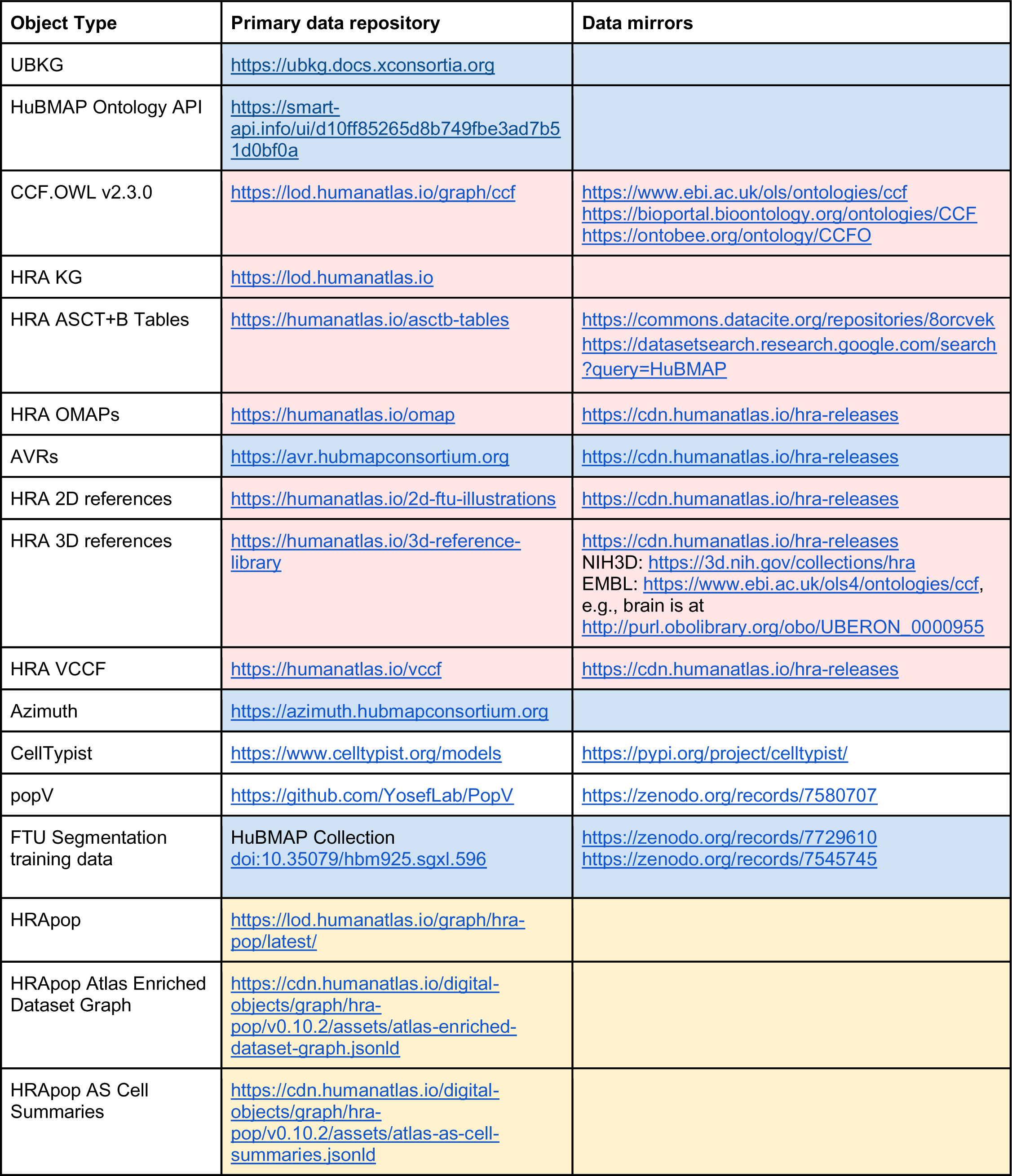
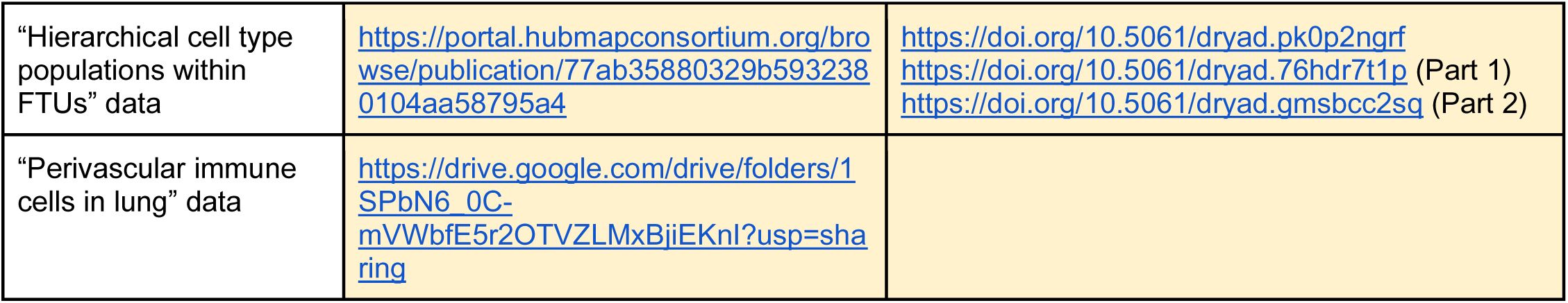
Primary and Secondary HRA Data Repositories. Color coding: Data used in HuBMAP Data Portal in blue, HRA Portal in red, in demonstration Previews in yellow, external code in white.

**Supplemental Table 3:** HRA Code Repositories. Color coding: Data used in HuBMAP Data Portal in blue, HRA Portal in red, in demonstration Previews in yellow, external code in white.

| Code Type and Name | GitHub Repository |
| --- | --- |
| **Data processing** |  |
| QuPath Manual Segmentation Tool (0.5.1) | https://qupath.github.io |
| Azimuth (0.4.6) | https://github.com/hubmapconsortium/azimuth-annotate |
| CellTypist (1.6) | https://github.com/Teichlab/celltypist |
| popV (0.9) | https://github.com/YosefLab/PopV |
| van Valen Tools | https://github.com/vanvalenlab?q=deep |
| van Valen's Cell Type Annotation (HuBMAP internal) | https://github.com/hubmapconsortium/deepcelltypes-hubmap |
| HuBMAP sc/snRNA-seq and CODEX pipelines | https://github.com/hubmapconsortium/salmon-rnaseq<br>https://github.com/hubmapconsortium/codex-pipeline |
| 3DCellComposer (1.2) | https://github.com/murphygroup/3DCellComposer |
| CytoSpatio (1.0.0) | https://github.com/murphygroup/CytoSpatio |
| PanelOptimizer | https://github.com/murphygroup/CODEXPanelOptimization |
| CellSegmentationEvaluator (1.5) | https://github.com/murphygroup/CellSegmentationEvaluator |
| FTU segmentation via Kaggle #1-2 Competitions | https://github.com/hubmapconsortium/pas-ftu-segmentation-pipeline<br>https://github.com/cns-iu/hra-multiftu-segmentation-pipeline |
| Cell Neighborhood Analysis | https://github.com/HickeyLab/Hierarchical-Tissue-Unit-Annotation |
| STELLAR | https://github.com/snap-stanford/stellar |
| STalign (1.0.1) | https://github.com/JEFworks-Lab/STalign |
| HRA Dataset Graphs Library (1.0) | https://github.com/hubmapconsortium/hra-rui-locations-processor |
| HRA Digital Object Processor | https://github.com/hubmapconsortium/hra-do-processor |
| HRA Workflows | https://github.com/hubmapconsortium/hra-workflows |
| HRA Workflows Runner | https://github.com/hubmapconsortium/hra-workflows-runner |
| HRA Multi-LOD Renderer (1.0.0) | https://github.com/cns-iu/hra-multi-lod |
| HRA AMAP | https://github.com/cns-iu/hra-amap |
| VCCF Visualizations | https://github.com/cns-iu/hra-vccf-cell-distance-visualizations |
| Vascular distance for 3D reconstruction of skin and spatial mapping of immune cell density | https://github.com/hubmapconsortium/vccf-visualization-2022 |
| EBI ASCT+B Table Validation (2024-07-11) | https://github.com/hubmapconsortium/ccf-validation-tools |
| HRA 3D Reference Organ Validation | https://github.com/hubmapconsortium/hra-ref-organ-validation |
| **API for data access** |  |
| HuBMAP Cells API | https://github.com/hubmapconsortium/cross_modality_query |
| HuBMAP Cells API (Python Client) (0.0.11) | https://github.com/hubmapconsortium/hubmap-api-py-client |
| HuBMAP Entity API (2.3.15) | https://github.com/hubmapconsortium/entity-api |
| HuBMAP Ingest API (2.3.17) | https://github.com/hubmapconsortium/ingest-api |
| HuBMAP Search API (3.3.12) | https://github.com/hubmapconsortium/search-api |
| API and SPARQL queries using the HRA-KG | https://github.com/hubmapconsortium/ccf-grlc |
| Ontology API for HuBMAP and SenNet applications (2.0.3) (extends UBKG API [2.1.4]) | https://github.com/x-atlas-consortia/hs-ontology-api<br>https://github.com/x-atlas-consortia/ubkg-api |
| HuBMAP UUID API (2.4.3) | https://github.com/x-atlas-consortia/uuid-api |
| HRA API (0.8.0) | https://github.com/x-atlas-consortia/hra-api |
| Tissue Block Annotation (1.0.0) | https://github.com/hubmapconsortium/hra-tissue-block-annotation |
| **User interfaces** |  |
| HuBMAP Consortium | https://hubmapconsortium.org |
| HuBMAP Data Portal (0.102.5) | https://github.com/hubmapconsortium/portal-ui,<br>https://portal.hubmapconsortium.org |
| HRA Portal | https://github.com/hubmapconsortium/hra-ui/tree/main/apps/humanatlas.io |
| Vitessce (3.4.6) | https://github.com/vitessce/vitessce |
| ASCT+B Reporter (2.8) | https://github.com/hubmapconsortium/hra-ui |
| EUI (3.8) | https://github.com/hubmapconsortium/hra-ui |
| RUI (3.8) | https://github.com/hubmapconsortium/hra-ui |
| FTU Explorer (0.5.0) | https://github.com/hubmapconsortium/hra-ui |
| HRA Organ Gallery (0.11.2) | https://github.com/cns-iu/hra-organ-gallery-in-vr |
| HRA Data Dashboard (0.1.0) | https://github.com/hubmapconsortium/hra-data-dashboard |
| VCCF Cell Distance Visualizations (0.1.0) | https://github.com/hubmapconsortium/hra-ui |
| Learn about the HRA through scrollytelling! | https://github.com/cns-iu/hra-scrollytelling |
| HRA Pilot Previews | https://github.com/hubmapconsortium/hra-previews |
| **Data / Ontologies** |  |
| HuBMAP Ontology (CCF v1.x) | https://github.com/hubmapconsortium/hubmap-ontology |
| Human Reference Atlas Knowledge Graph (HRA-KG) | https://github.com/hubmapconsortium/hra-kg |
| HRA Vasculature CCF | https://github.com/hubmapconsortium/hra-vccf |
| HRA Vocabulary (2.5.10) | https://github.com/hubmapconsortium/hra-vocab |
| HRApop (0.10.2) | https://github.com/x-atlas-consortia/hra-pop |
| HRAlit (0.5) | https://github.com/cns-iu/hra-literature |

## Notes

https://cns-iu.github.io/hra-construction-usage-supporting-information/

### Competing Interest Statement

Core authors:
In the past 3 years, RS has received compensation from Bristol-Myers Squibb, ImmunAI, Resolve Biosciences, Nanostring, 10X Genomics, Neptune Bio, and the NYC Pandemic Response Lab. RS is a co-founder and equity holder of Neptune Bio.
Sarah A Teichmann is a remunerated member of the Scientific Advisory Boards of Qiagen, Foresite Labs and Element Biosciences, a co-founder and equity holder of TransitionBio and EnsoCell Therapeutics, and a part-time employee of GlaxoSmithKline since January 2024.
HRA Team authors:
Bruce J Aronow: Nexstone Immunology, Uniquity, Advisors
Christopher Werlein: Speaker fee for Boehringer Ingelheim
Michael Snyder: Personalis, SensOmics, Qbio, January AI, Fodsel, Filtricine, Protos, RTHM, Iollo, Marble Therapeutics, Crosshair Therapeutics, NextThought and Mirvie, Jupiter, Neuvivo, Swaza, Mitrix, Yuvan, TranscribeGlass, Applied Cognition
Neil L Kelleher: Thermo Fisher Scientific, Proteinaceous, Integrated Protein Technologies, ImmPro

### Summary of Updates

We made changes throughout the paper to streamline the presentation while keeping biologist readers in mind.

