## Supplementary material for "Human BioMolecular Atlas Program (HuBMAP): 3D Human Reference Atlas Construction and Usage": ZIP file with 17 supplemental figures: 8 v3.12.2024.pdf

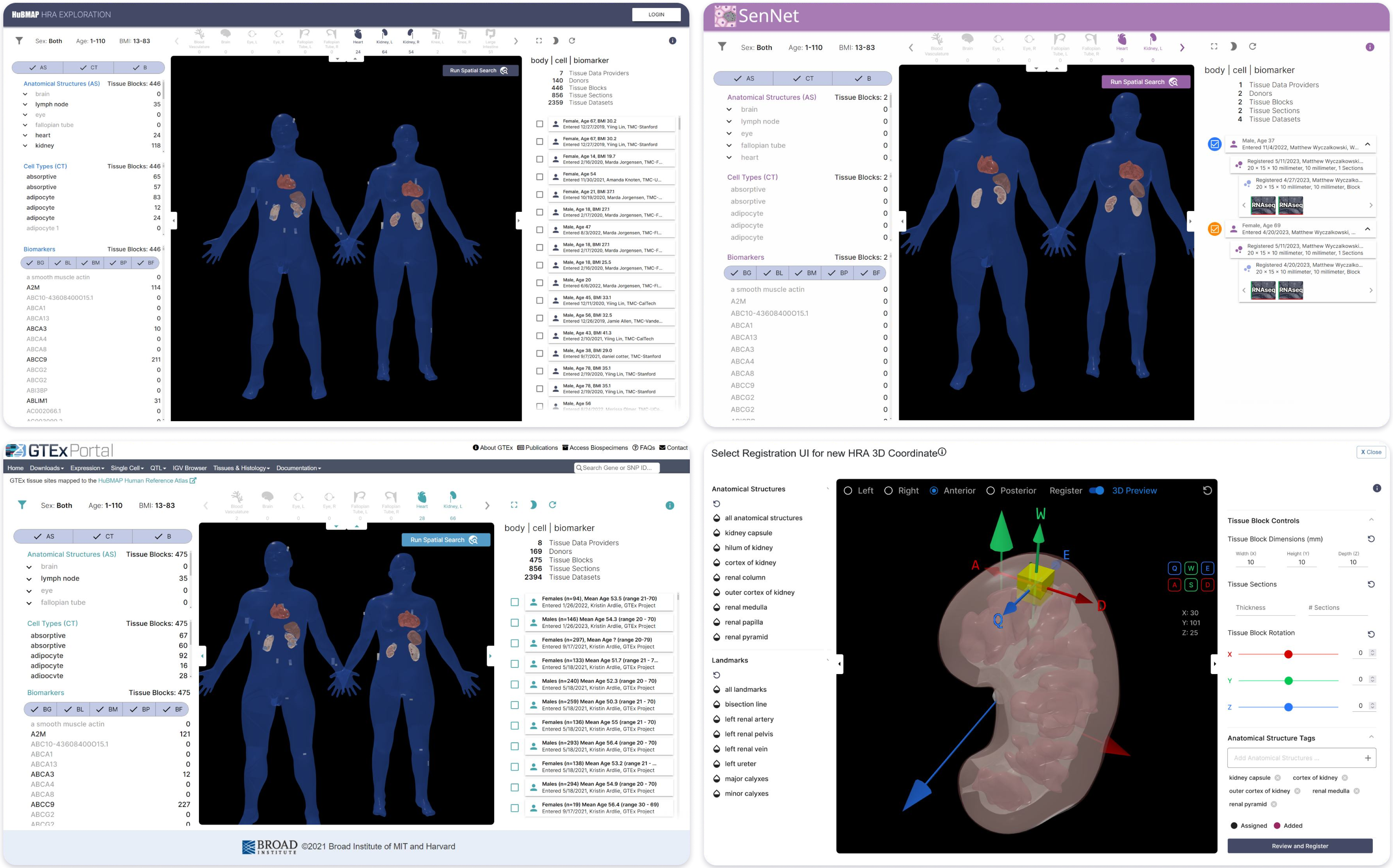


**Supplemental Figure 8: Customized, Branded Deployment of EUI in HuBMAP, SenNet, GTEx, and RUI in GUDMAP**
