## Supplementary material for "Human BioMolecular Atlas Program (HuBMAP): 3D Human Reference Atlas Construction and Usage": ZIP file with 17 supplemental figures: 11 v7.26.2024.pdf

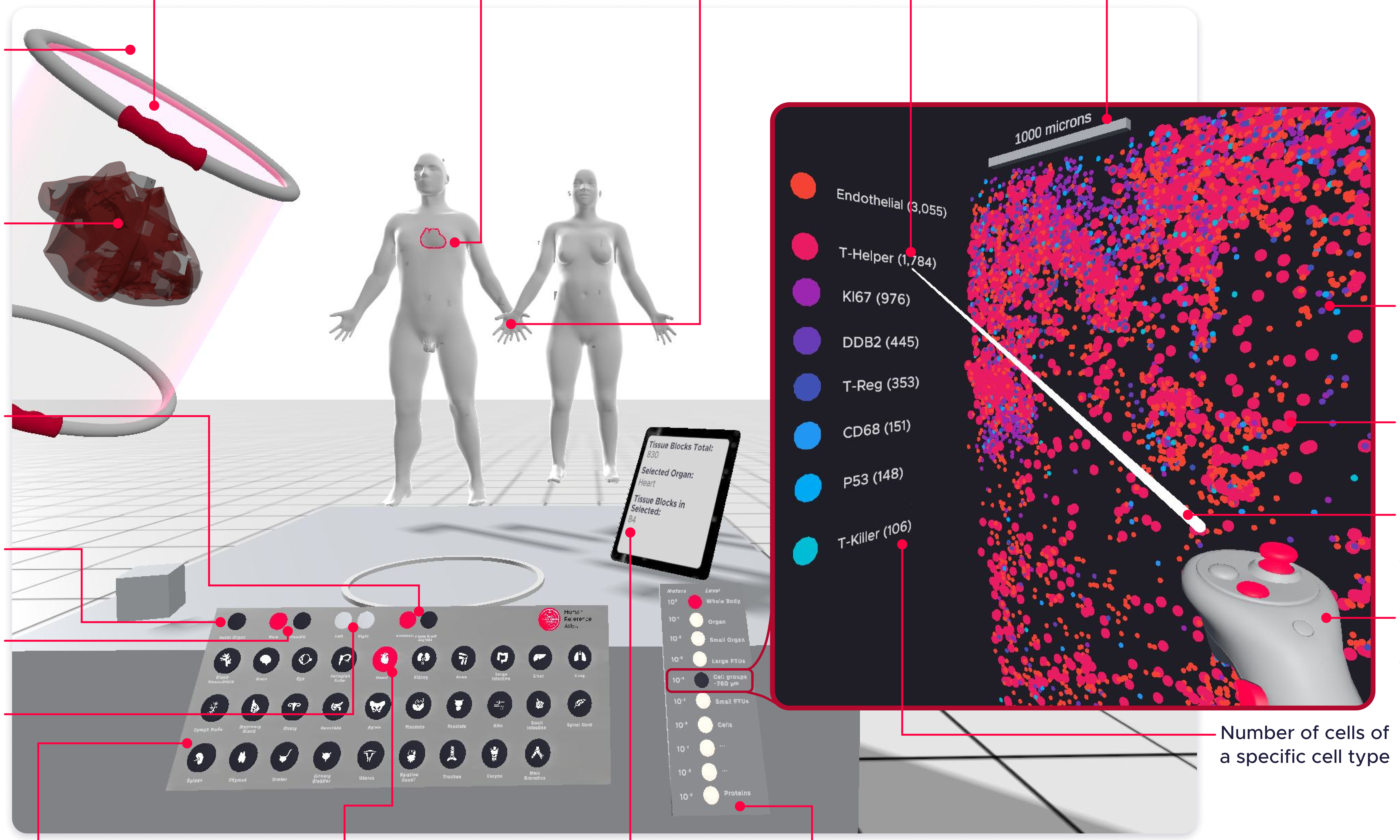


Credits for the single-cell data shown on the right side:
Source: Ghose S, Ju Y, et al. 3D reconstruction of skin and spatial mapping of immune cell density, vascular distance and effects of sun exposure and aging (2023)
DOI: 10.1038/s42003-023-04991-z

**Supplemental Figure 11: HRA Organ Gallery**
