## Supplementary material for "Human BioMolecular Atlas Program (HuBMAP): 3D Human Reference Atlas Construction and Usage": ZIP file with 17 supplemental figures: 14 v3.12.2024.pdf

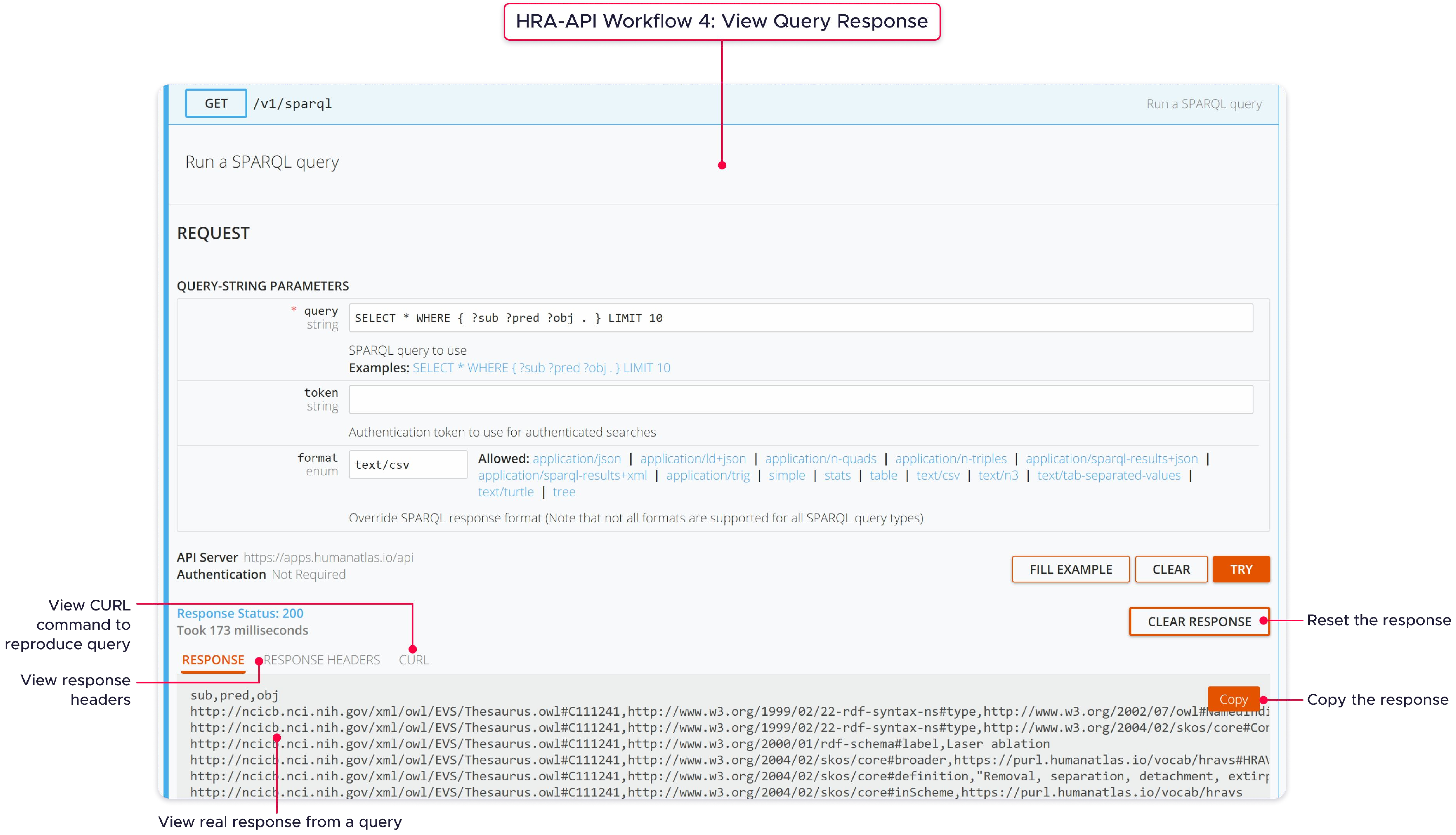


**Supplemental Figure 14: Human Reference Atlas Application Programming Interface: View Query Response**
