## Supplementary material for "Human BioMolecular Atlas Program (HuBMAP): 3D Human Reference Atlas Construction and Usage": ZIP file with 17 supplemental figures: 17 v3.12.2024.pdf

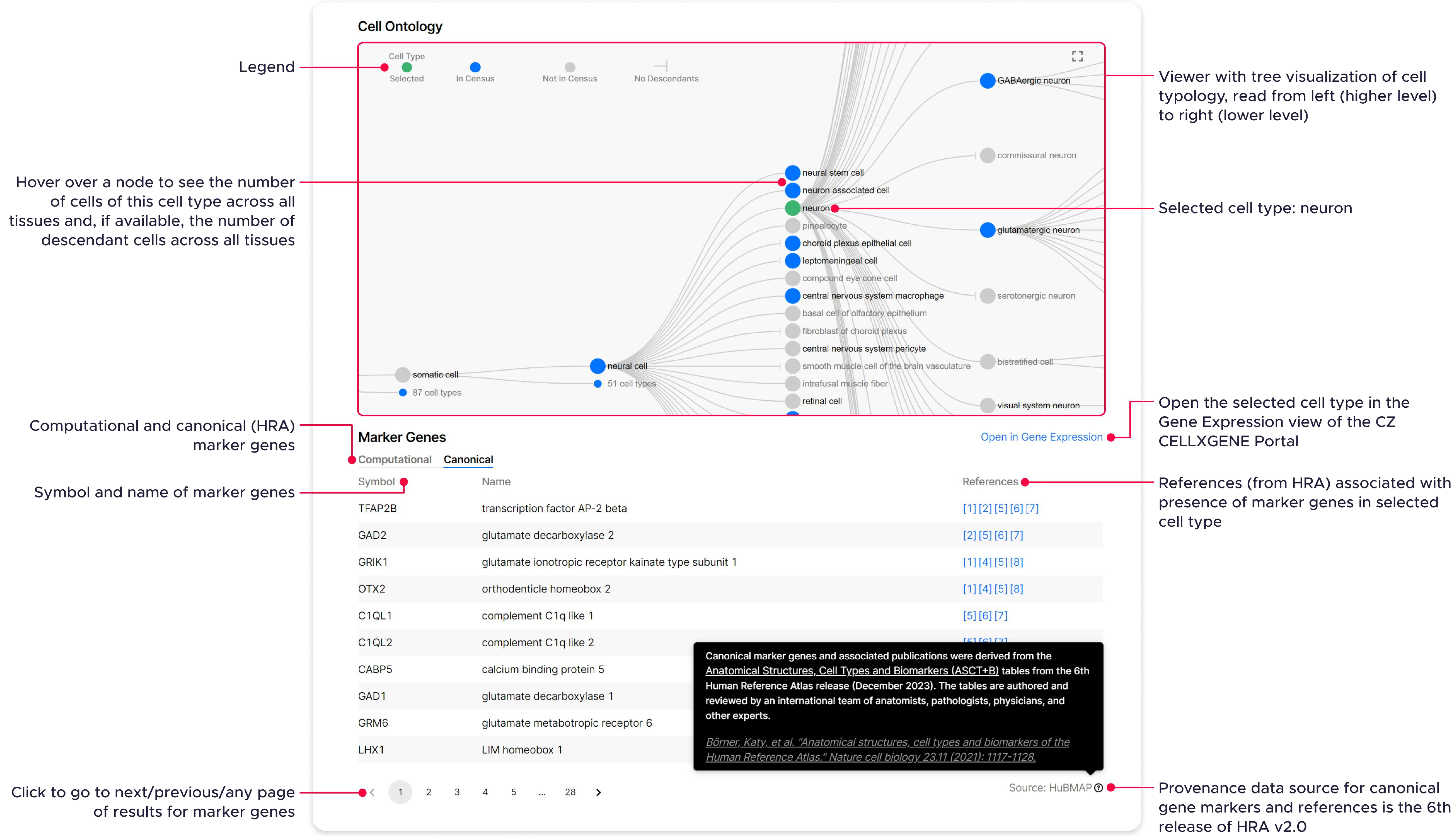


**Supplemental Figure 17: CZ CellGuide Visualization With ‘Canonical’ Marker Genes And ‘References’ From The HRA**
