## Supplementary figures and images for "Human BioMolecular Atlas Program (HuBMAP): 3D Human Reference Atlas Construction and Usage"

### 1 v3.12.2024.pdf

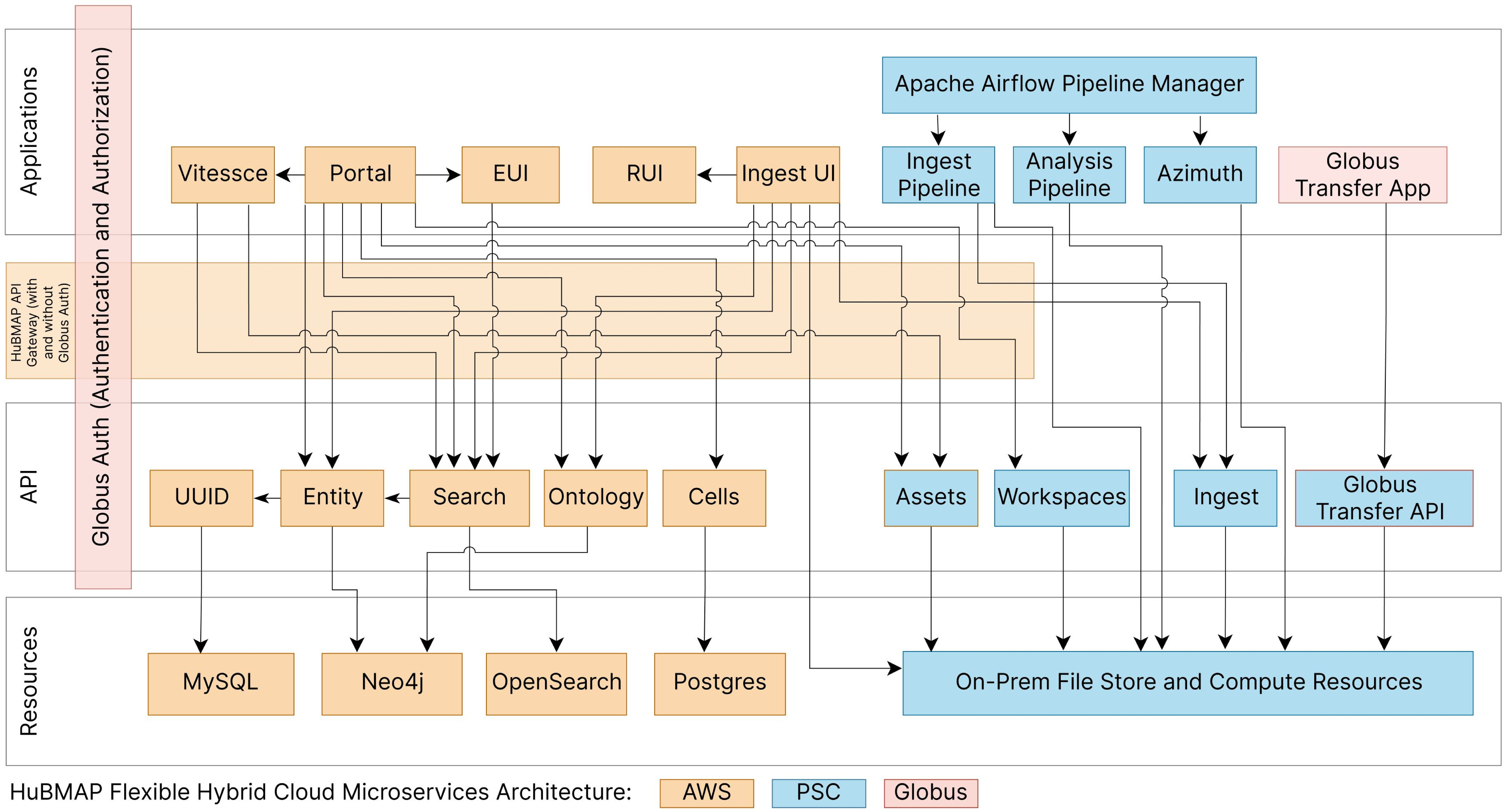


**Supplemental Figure 1. Hybrid Cloud Microservices System Architecture**

### 2 v5.20.2024.pdf

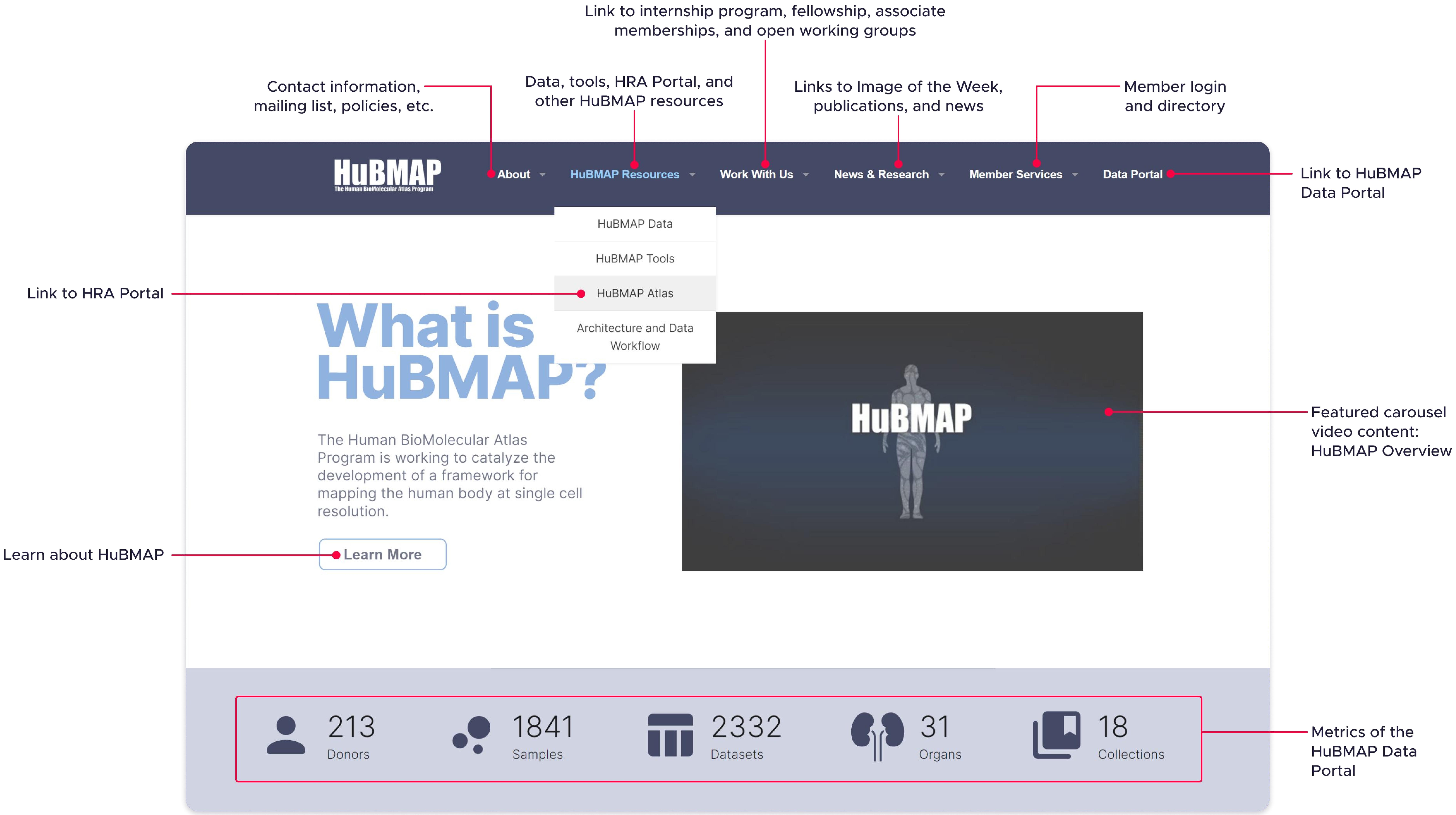


**Supplemental Figure 2. HuBMAP Consortium Website**

### 3 v3.12.2024.pdf

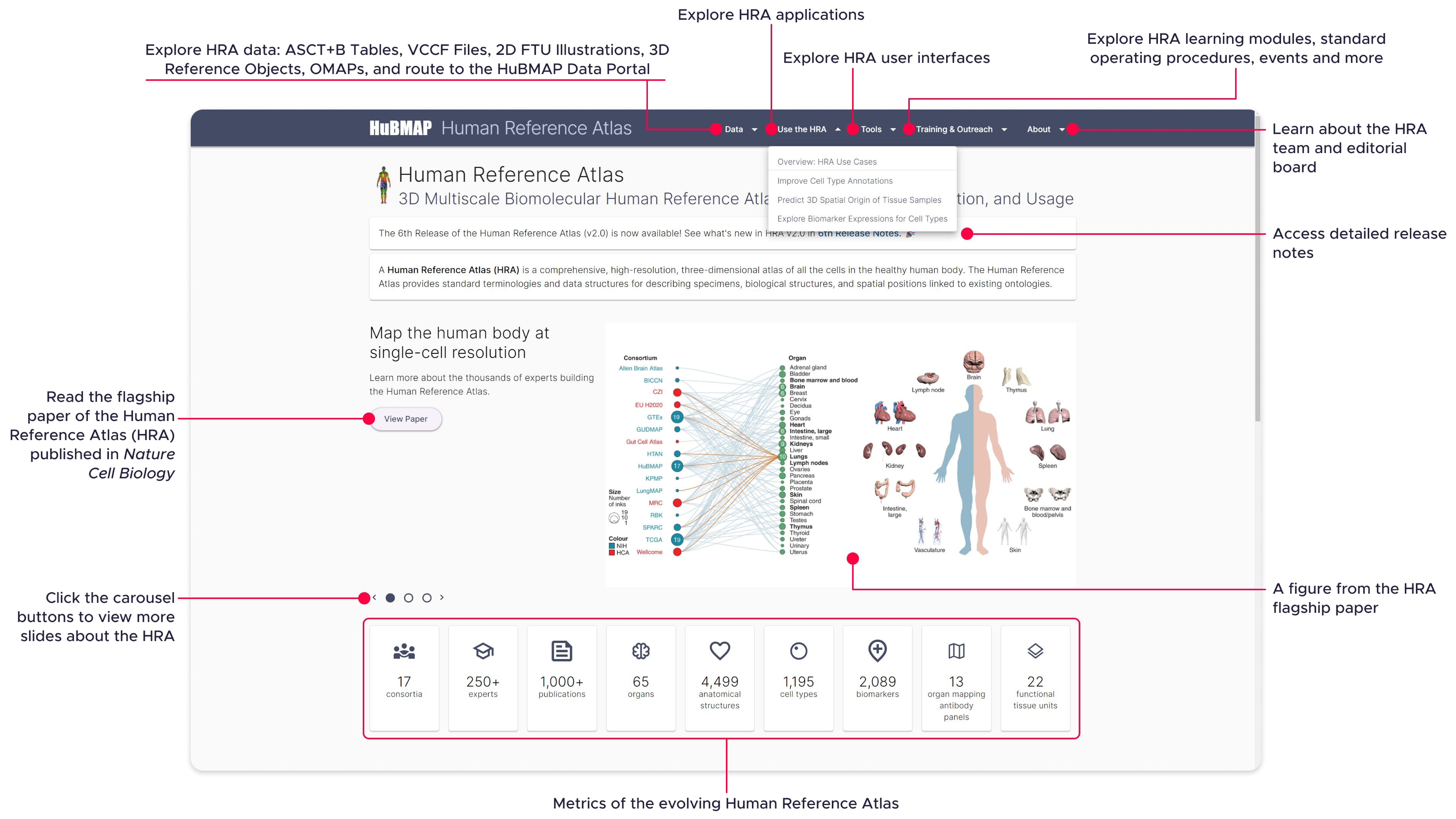


**Supplemental Figure 3. Human Reference Atlas Portal**

### 4 v3.12.2024.pdf

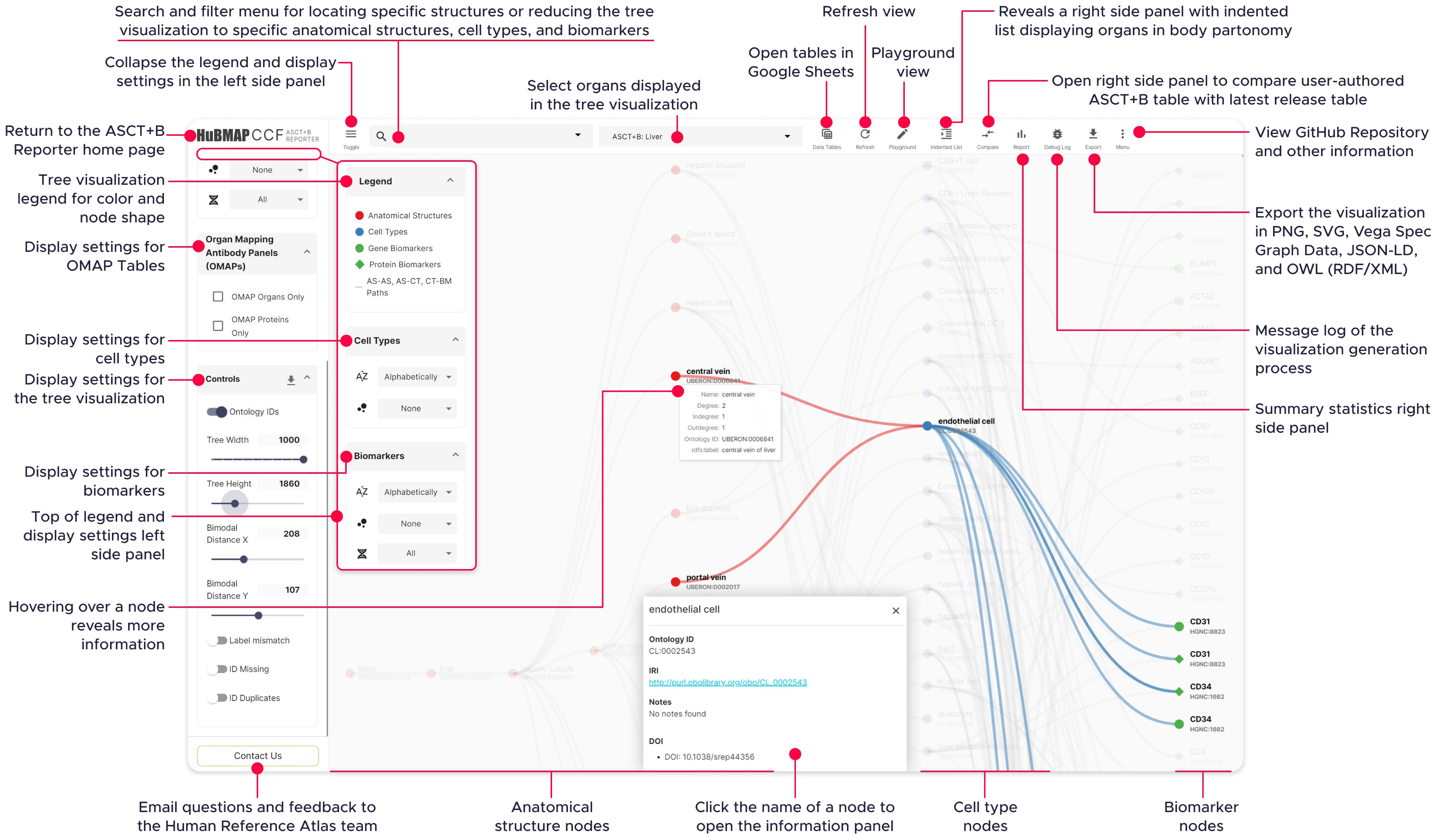


**Supplemental Figure 4: ASCT+B Reporter User Interface**

### 5 v3.12.2024.pdf

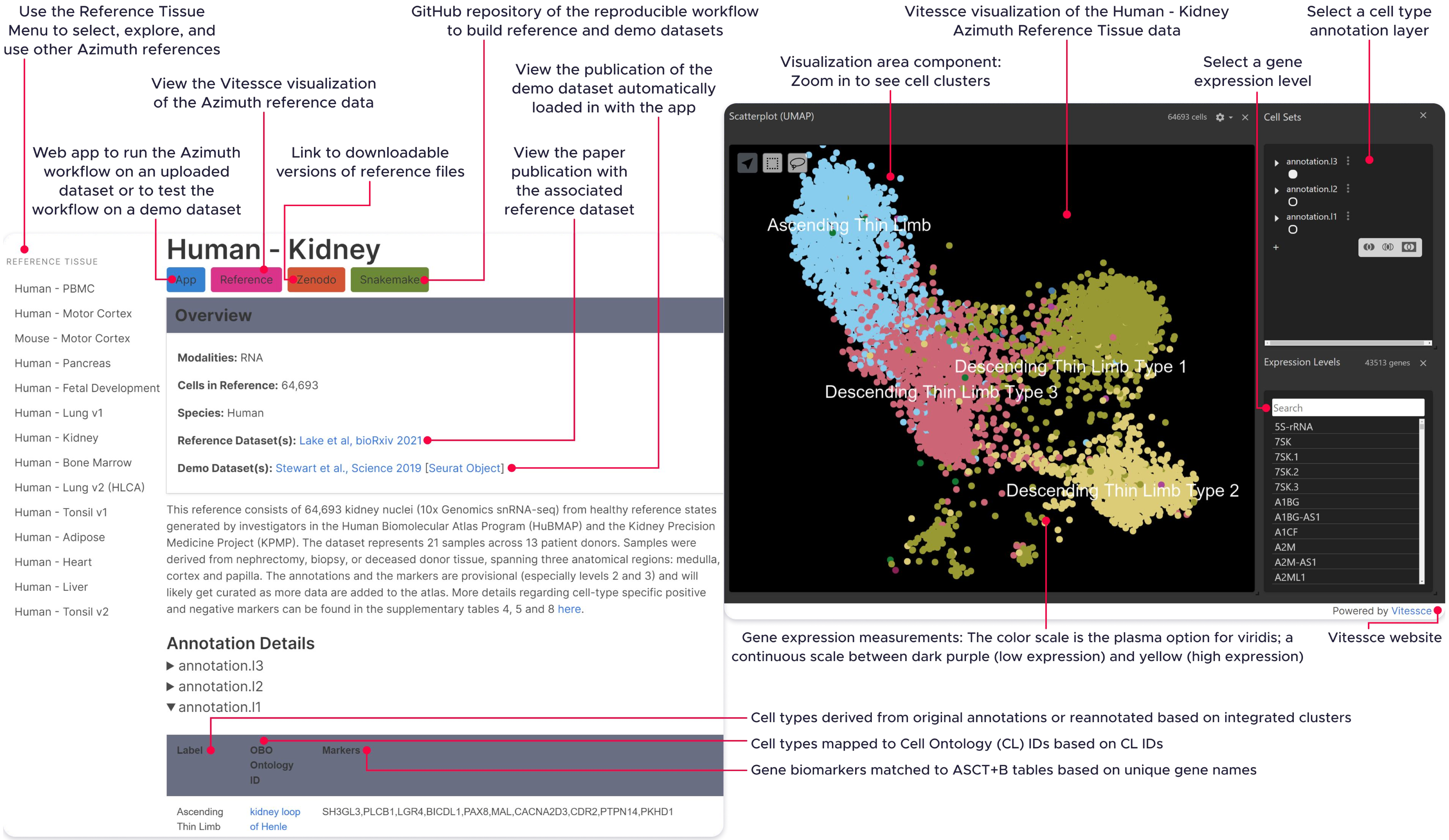


**Supplemental Figure 5. Azimuth Portal and Reference Explorer User Interface**

### 6 v3.12.2024.pdf

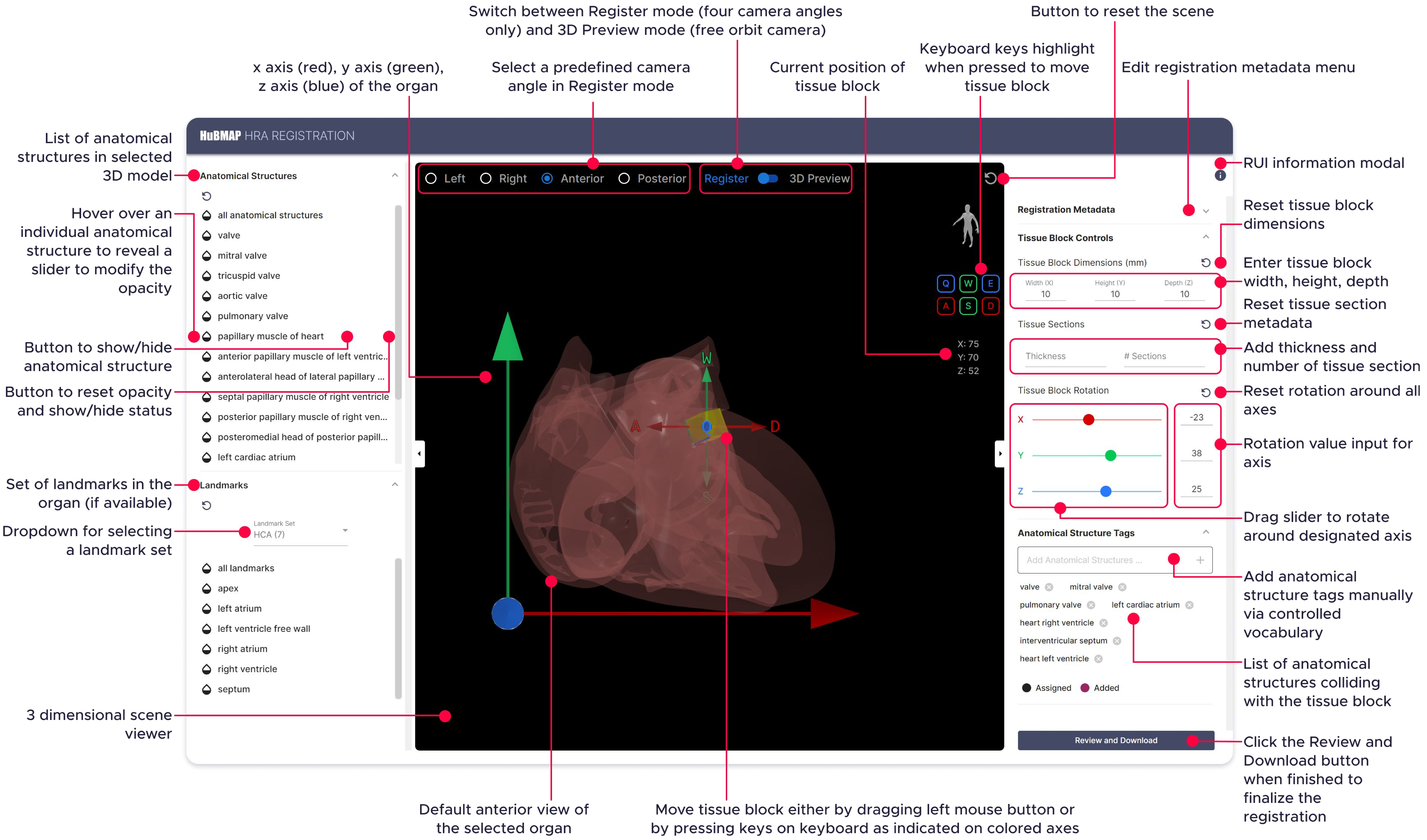


**Supplemental Figure 6: Registration User Interface (RUI)**

### 7 v3.12.2024.pdf

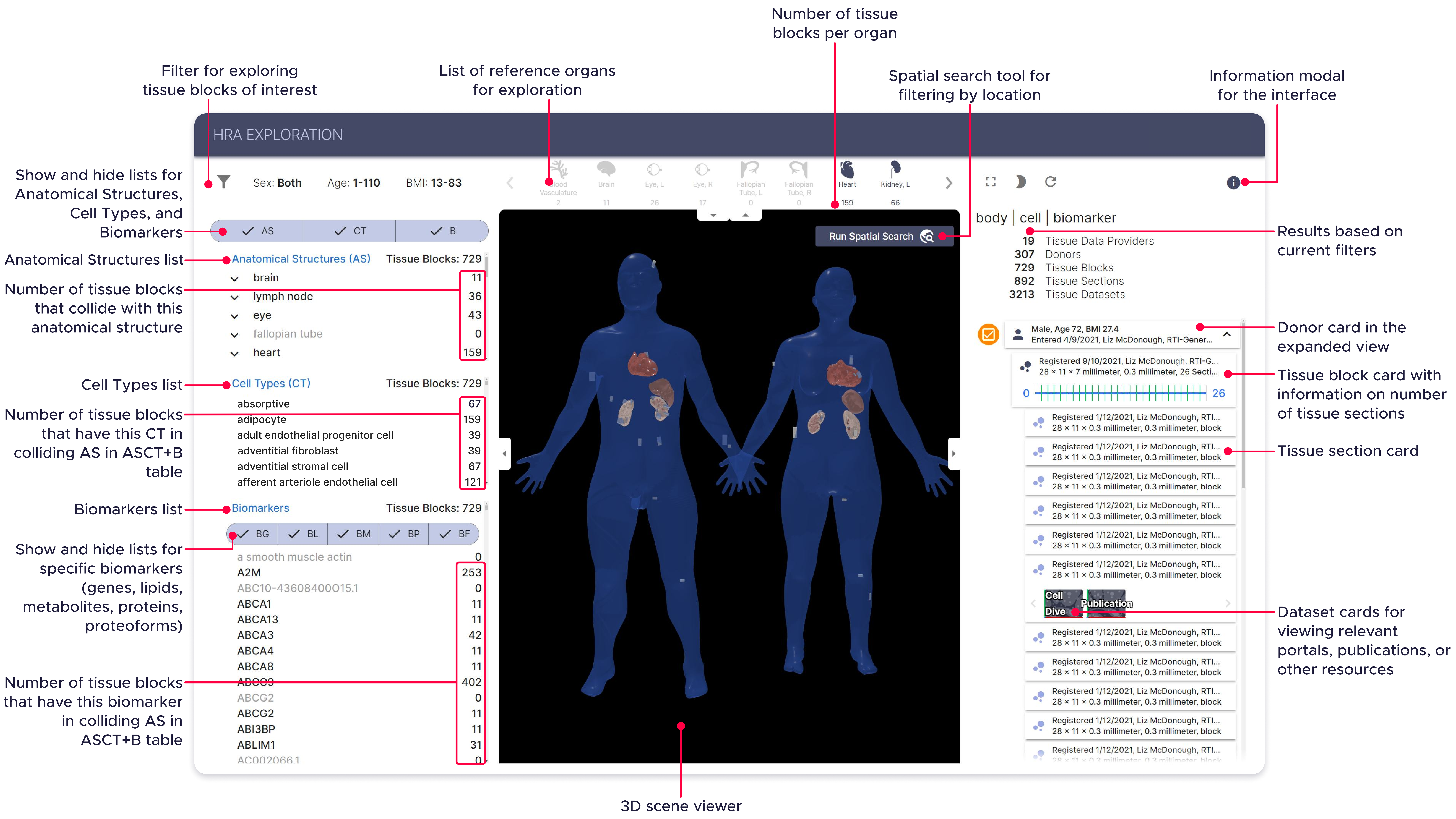


**Supplemental Figure 7: Exploration User Interface (EUI)**

### 9 v3.12.2024.pdf

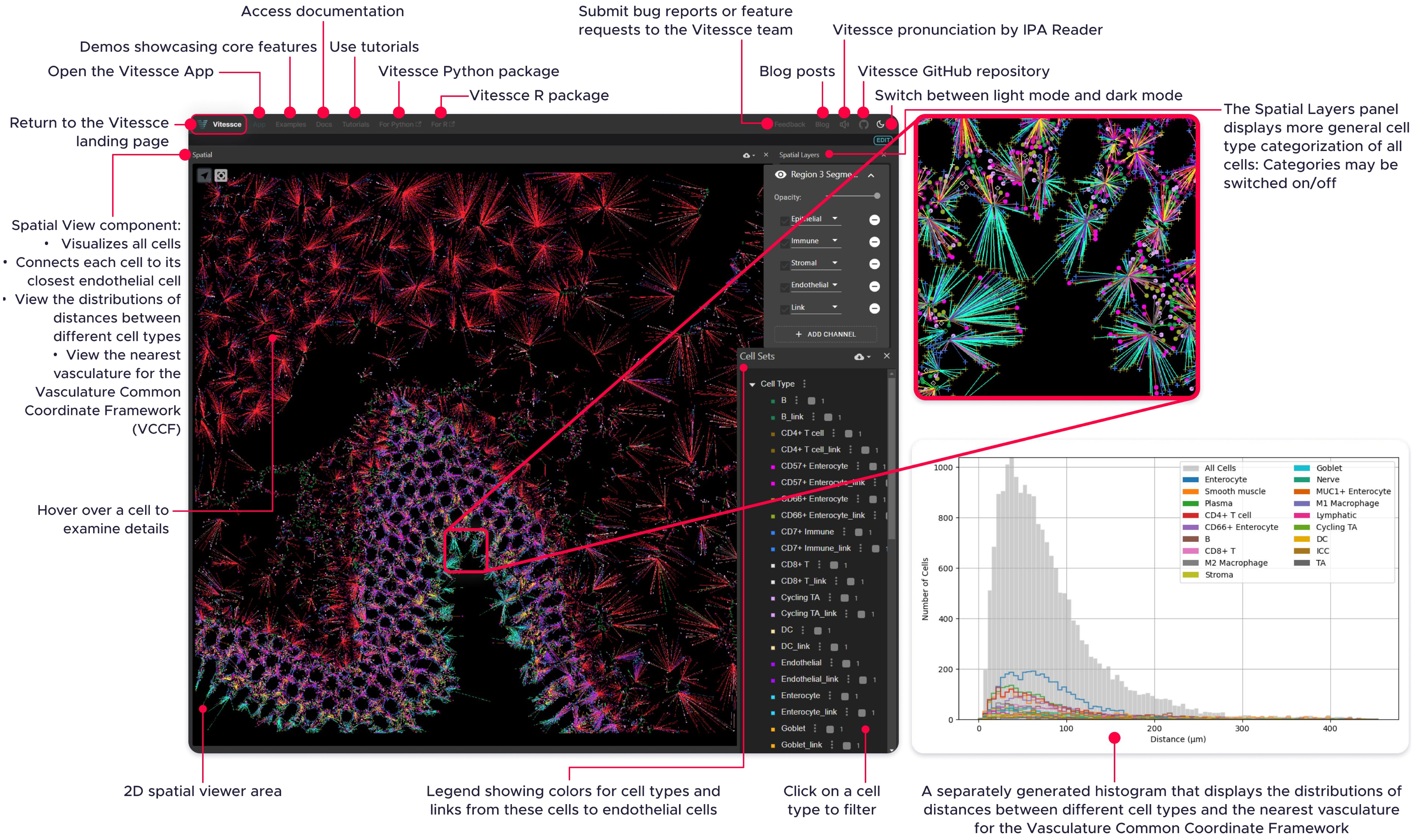


**Supplemental Figure 9: Vasculature Common Coordinate Framework Distance Visualizations**

### 10 v8.12.2024 v2.pdf

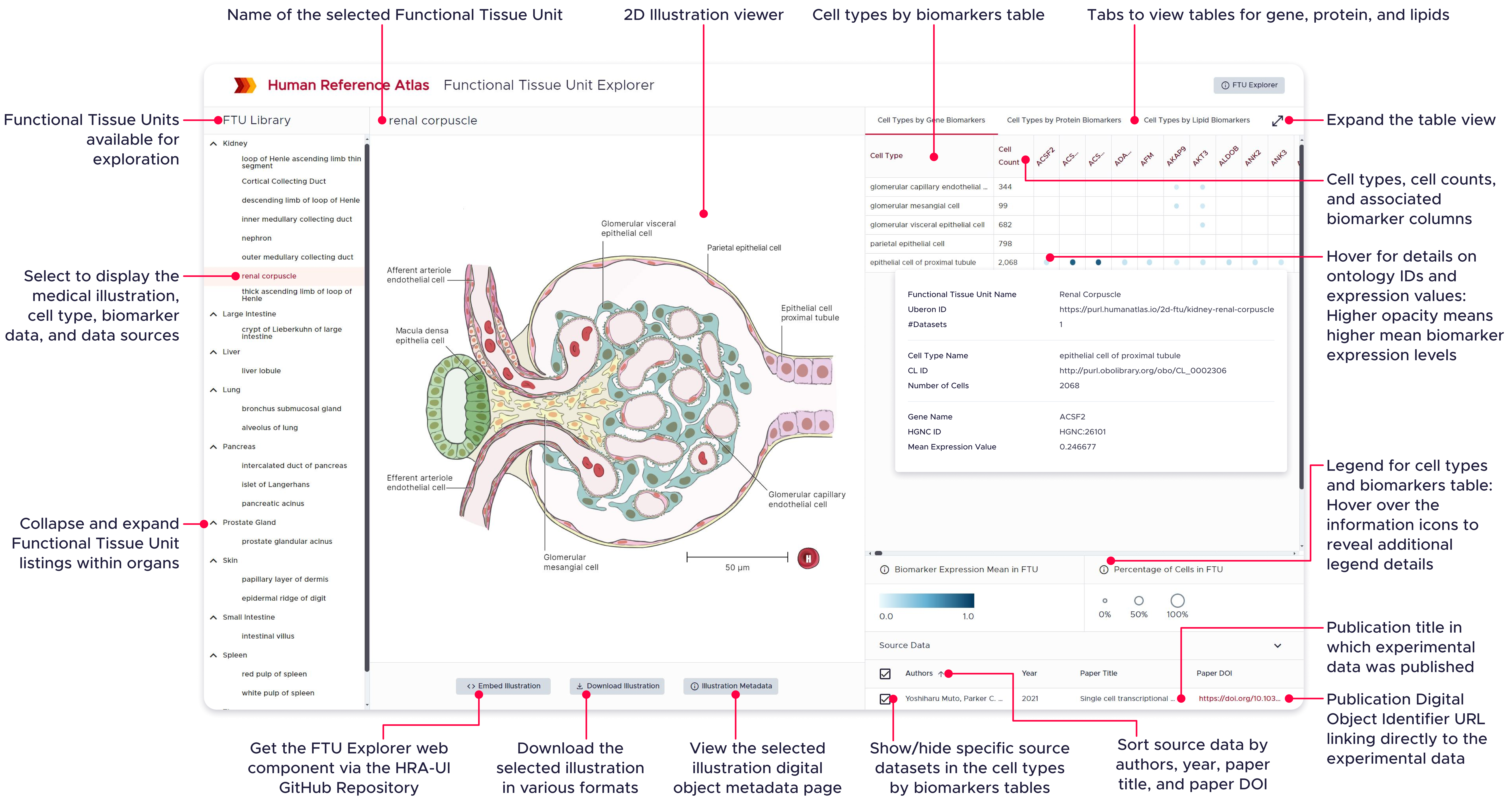


**Supplemental Figure 10: Interactive FTU Explorer**

### 15 v7.15.2024.pdf

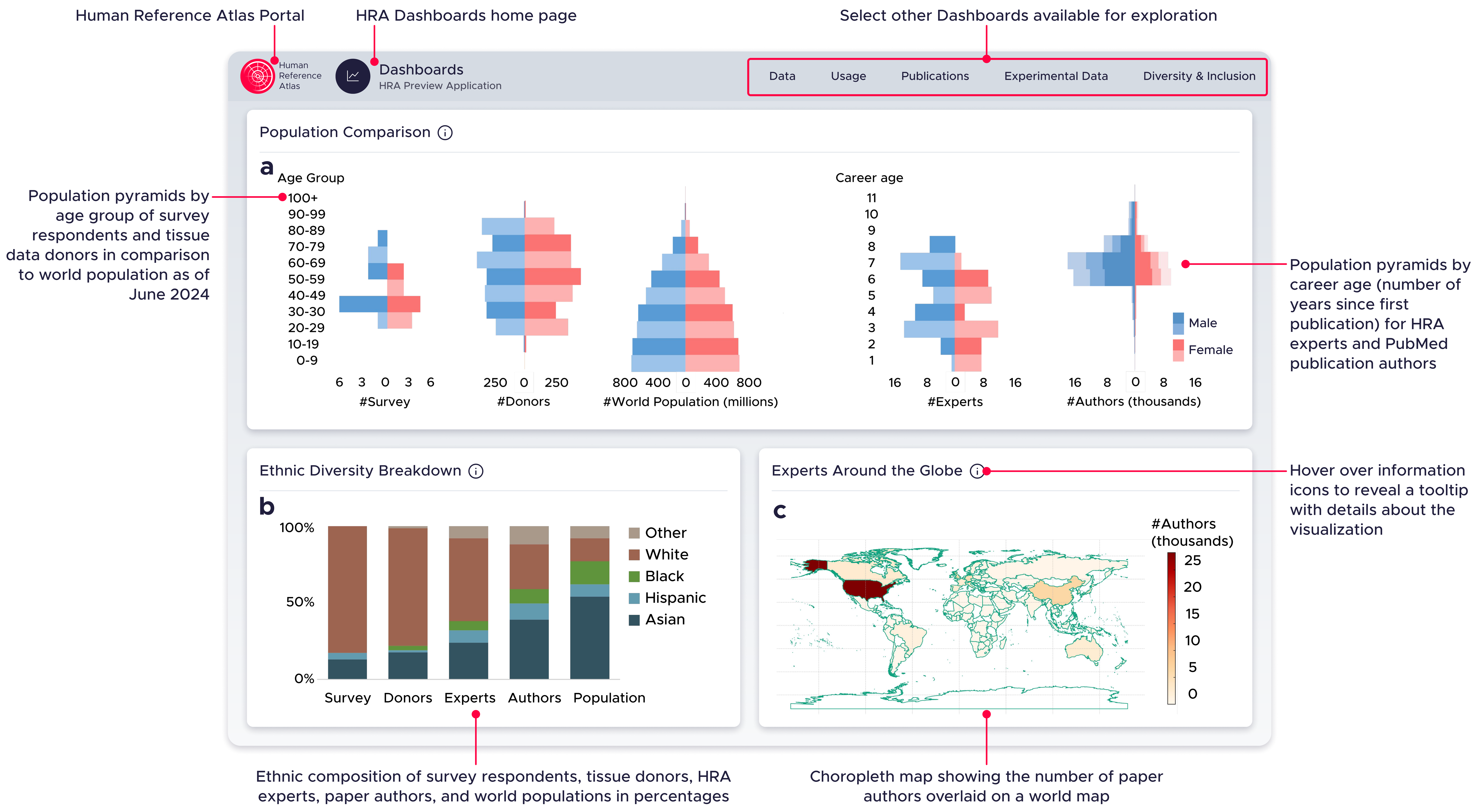


**Supplemental Figure 15. HRA Equity Dashboard**

### 16 v3.12.2024.pdf

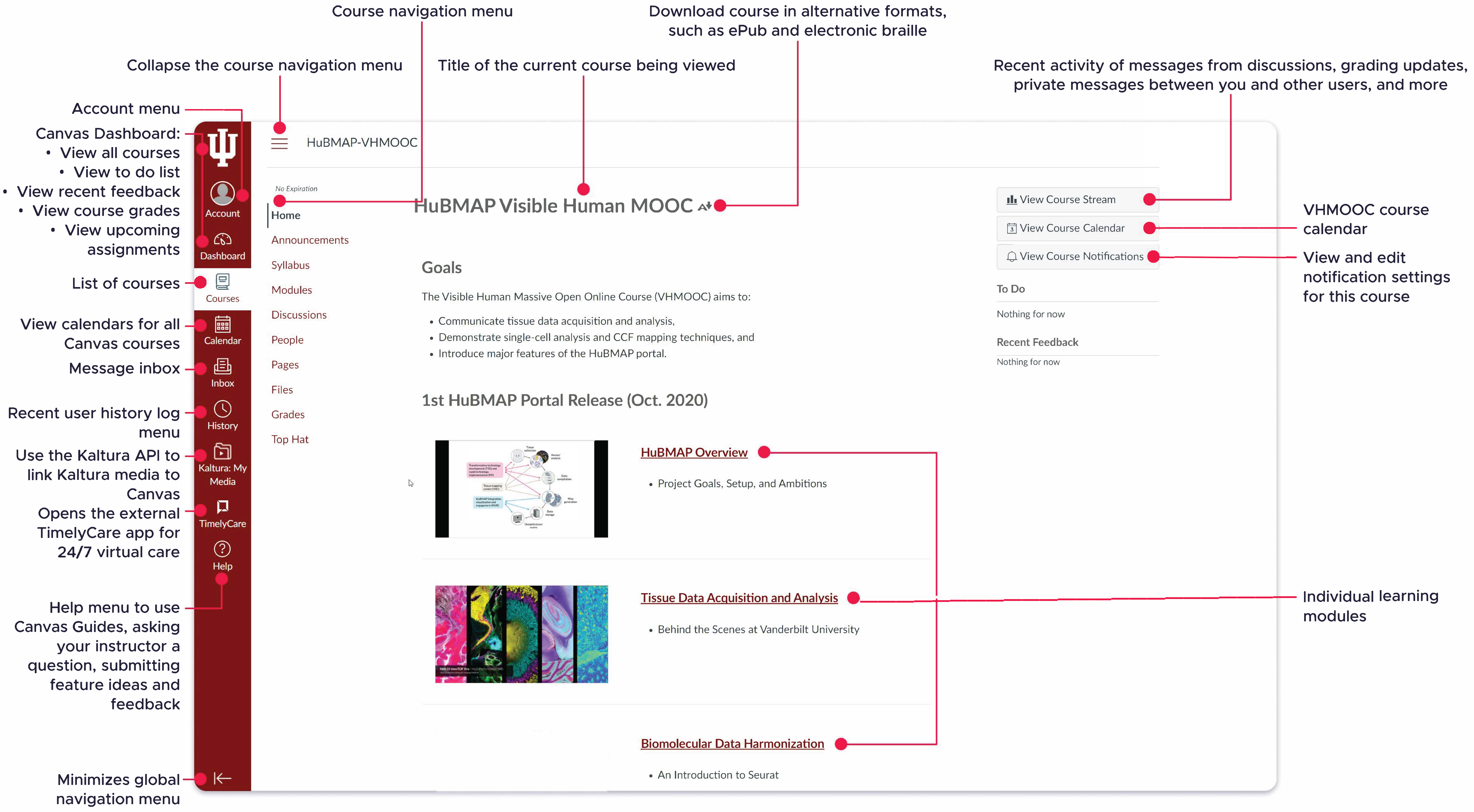


**Supplemental Figure 16: Visible Human Massive Open Online Course (MOOC)**
